# Computer-Vision Stabilized Intravital Imaging Reveals Lung Capillary Neutrophil Dynamics Crucial for Lung Host-Defense Function

**DOI:** 10.1101/2021.04.19.440486

**Authors:** Yoshikazu Tsukasaki, Kurt Bachmaier, Jagdish C. Joshi, Bhagwati Joshi, Peter T. Toth, Zhigang Hong, Saroj Nepal, Matthias Gunzer, Jaehyung Cho, Dolly Mehta, Chinnaswamy Tiruppathi, Sandra Pinho, Gary Mo, Jalees Rehman, Asrar B. Malik

## Abstract

Polymorphonuclear neutrophils (PMN) are highly dynamic innate immune cells which are essential for lung host defense. However, *in vivo* intravital imaging in moving organs such as the lung remains challenging due to motion artifacts. Here we describe a novel intravital imaging method with high-throughput analytical capability based on a computer vision stabilization algorithm, Computer-vision-Assisted STabilized intravital imaging (CASTii). The sub-micron precision of this approach enables analysis of compartmentalized intravital PMN dynamics. We quantified in real-time a novel patrolling function of lung intracapillary circulating PMN. We also describe the dynamics of intracapillary PMN pooling (marginated PMN pool) using direct imaging of PMNs. The pool was formed by repeated catch-and-release kinetics involving PMN deformation inside microvessels during the passage of PMNs in vessels. We observed rapid PMN recruitment into the lung tissue compartments from pooled PMNs in response to alveolar chemoattract exposure. In contrast, endotoxemia-induced intracapillary sequestration of PMN impaired PMN transmigration into the alveolar space and defective phagocytosis of live bacteria. Intravital imaging of PMN dynamics with CASTii provides fundamental insights into host-defense functions of lung capillary PMN.

## Introduction

The lung is constantly exposed to environmental pathogens requiring continuing maintenance of a sterile environment by immune cells such as polymorphonuclear neutrophils (PMNs)(Giacalone, Margaroli, Mall, & Tirouvanziam, 2020; Whitsett & Alenghat, 2015). PMNs in the lung are considered as the first defender cells that rapidly migrate into tissue and kill pathogens with defined temporal and spatial patterns optimized for lung host defense(Kolaczkowska & Kubes, 2013; Petri, Phillipson, & Kubes, 2008; Yipp et al., 2017). Such spatio-temporal functions of PMNs have been investigated using confocal and multiphoton fluorescence intravital imaging in living tissue with sub-micron/milliseconds scales(McDonald et al., 2008; Svoboda, Denk, Kleinfeld, & Tank, 1997; Yipp et al., 2017). Respiratory motion artifacts however severely limit image registration leading to analytical inefficiencies(Yipp et al., 2017). In the present studies we applied a computational solution to correct the motion artifacts. Our premise was that the ideal image-registration algorithm for lung intravital imaging should have the capability to correct both large movements and deformations with fast and accurate calculations. General image registration algorithms, such as cross-correlation methods (Dubbs, Guevara, & Yuste, 2016; Miri, Daie, Burdine, Aksay, & Tank, 2011) and phase-correlation methods with 2D Fourier transform (Parslow, Cardona, & Bryson-Richardson, 2014) cannot be used for this purpose because they fail to account for lung tissue deformations. Algorithms with block-wise rigid (Pnevmatikakis & Giovannucci, 2017) and non-rigid transform (Dunn, Lorenz, Salama, & Delp, 2014; Vercauteren, Pennec, Perchant, & Ayache, 2009) can be used but require extensive computation time with heavy calculations and are thus unsuited for physiological studies requiring rapid assessment of changes in PMN dynamics. In contrast to these methods, computer-vision is a feature-based method described herein extracts features of the image and tracks these along time frames of a movie with less information quantity(Lucas & Kanade, 1981; Shi & Tomasi, 1994).

Studies have reported extravascular resident PMN(Kreisel et al., 2010) and PMN marginalized pools(Doerschuk et al., 1987; Lien et al., 1987; Lien et al., 1990) in lungs under resting conditions(Yipp et al., 2017). Extravascular resident PMNs migrating in lung tissue were described by intravital imaging(Kreisel et al., 2010). Studies also showed PMN marginated pools(Doerschuk et al., 1987; Lien et al., 1987; Lien et al., 1990) was located in capillaries and the pools easily changed their size and dynamics in response to hemodynamic alterations in lungs(Bierman et al., 1952; Bierman et al., 1951; Summers et al., 2010). In endotoxemic conditions, PMNs in lungs migrate throughout lung tissue in response to endotoxin(Kreisel et al., 2010; Yipp et al., 2017). However, the dynamics of tissue-compartmentalized PMN in lungs and their relevance to host-defense function remain poorly understood due to limited imaging capabilities. In this study, we applied the computer-vision image registration technique for motion correction with sub-micron precision to lung intravital imaging and configured a high-throughput intravital imaging platform (CASTii: Computer-vision assisted intravital imaging). We found direct evidence of a marginated PMN pool in the intracapillary space but no evidence of extravascular tissue-resident PMNs in mice in the resting state. PMN pooling dynamics were essential for rapid PMN transmigration into the alveolar space in response to chemotactic gradients. In endotoxemia, PMNs lost their migratory capabilities in the intracapillary space over time in a tightly choreographed manner. This surprisingly compromised PMN transmigration in response to chemotactic agents and killing of pathogens in the alveolar space, thereby resulting in defective host defense function of PMNs.

## Results

### Physiologically respirating lung intravital imaging

The current intravital imaging methods in lungs require a creation of an optical window at which high pleural pressures are applied to mechanically constrain the mobility of respiring lungs(Looney et al., 2011; Murphy, Renninger, & Gossett, 1998; Rodriguez-Tirado et al., 2016). To understand the impact of mechanical arrest on the vascular structures and dynamics of blood cells in vessels, we first investigated the effects of suction pressure applied at the optical window. We observed that increased pressures artifactually enlarged the alveolar area, altered the shape of pulmonary vessels (Supplementary figure 1a, b), and itself caused the accumulation of PMN (Supplementary figure 1a, c), We did not detect these change at physiological pleural pressures (−13.6 to −20.4 cmH_2_O)(Murphy et al., 1998) (Supplementary figure 1a-c). We also evaluated the relationship of applied suction pressure to changes in lung motion using optical flow analysis with sub-pixel resolution and precision (Supplementary figure 2a-c). We observed significant lung motion displacement and deformation that were inversely correlated with the applied pressure (Supplementary figure 1d, e). In addition, lung displacement and deformation with suction pressures of −13.6 to −20.4 cmH_2_O (representing physiological pleural pressures) occurred in an uneven manner and varied among preparations and field of views (FOVs) (Supplementary figure 3 and supplementary movie 1).

### Computer-vision Assisted STabilized intravital imaging (CASTii) achieves pulmonary compartmentalized PMN imaging with high-throughput analysis capability

To correct the lung structural changes and motion artifacts described above, we developed Computer-vision Assisted STabilized intravital imaging (CASTii). This method is based on a computer vision image-registration algorithm that does not require unphysiological pressures to stabilize lungs. Image processing of CASTii consists of three distinct steps: (i) feature points detection, (ii) motion tracking, and (iii) motion correction involving averaging of respiratory cycles (Figure 1a and supplementary movie 2).

**Figure 1.**
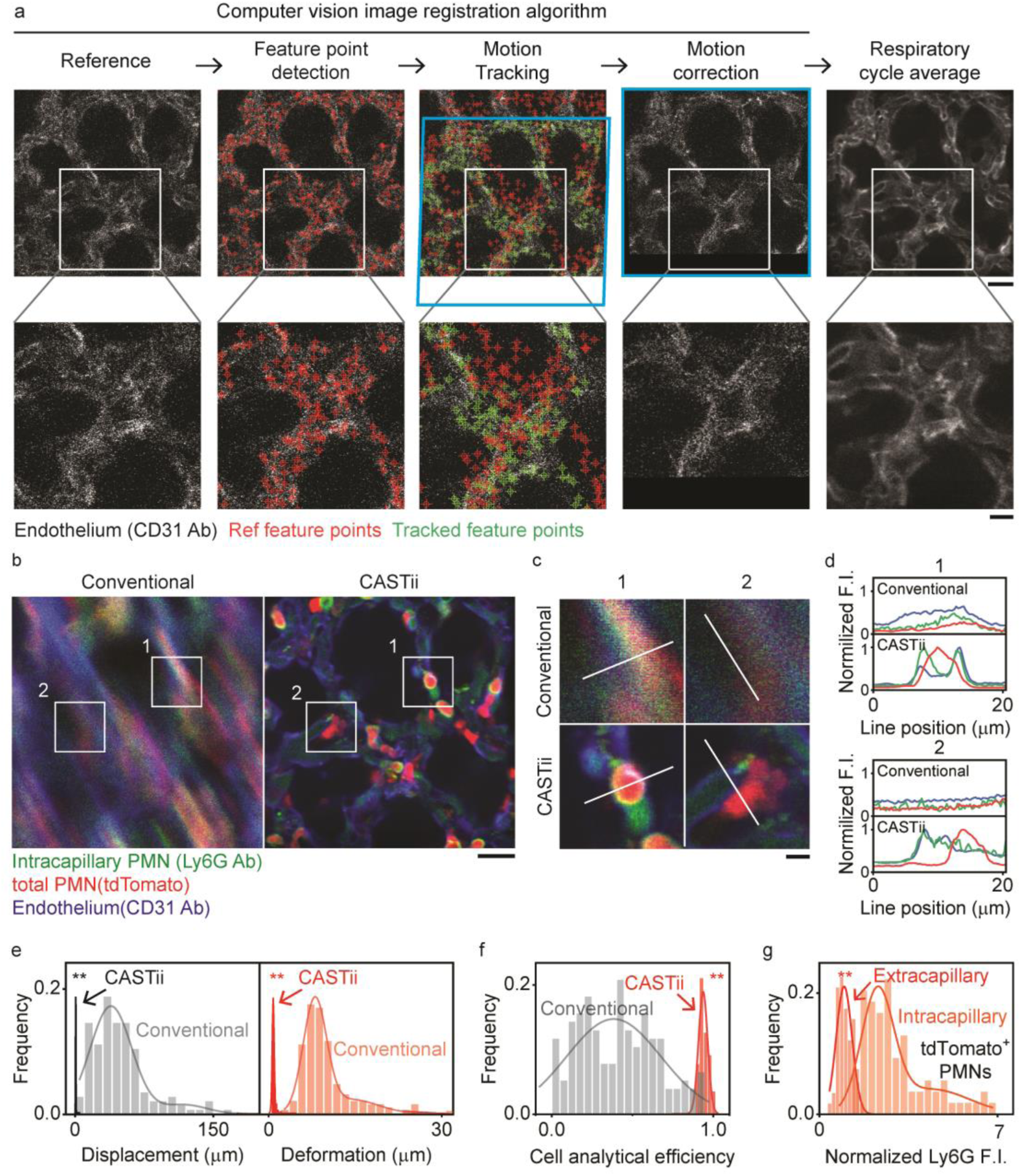
Lung intravital imaging in normally respiring lungs using CASTii. **a.** Flow chart of the stabilization processing of CASTii using an example set of lung images. Pulmonary microvessels are imaged; feature points are detected algorithmically in designated reference image (red crosses indicate the feature points). Feature points are tracked to calculate their deviation from reference (green crosses indicate tracked feature points). This estimated motion is corrected and followed by averaging over the respiratory cycle. Inset white boxes in the top row are magnified at the bottom row. Scale bar; 20 μm (Top) and 10 μm (Bottom). **b.** PMN imaging using CASTii after 4h of i.t. LPS demonstrates clear gain in precise stabilization. Left image: Conventional, right image: using CASTii. Scale bar; 20 μm. **c.** Magnified view from inset white boxes in b. Scale bar; 5 μm. d, Line profile analysis of PMNs and fine lung structures along white lines drawn in c. **e.** Quantification of motion displacement and deformation via peak-to-peak movements using conventional methods and CASTii; n=143 (FOV). **f.** Quantification of cell analytical efficiency with conventional methods and CASTii; n=143 (FOV). **g.** Compartmental PMN localization with Ly6G fluorescent intensity; n=58 (Cell, Extracapillary PMN) and n=77 (Cell, Intracapillary PMN). Statistical analysis was performed using two-sample t-test in e-g. All two-photon images are single z slices.

We acquired motion of lungs by a video-rate resonance two-photon microscope. Using the Shi and Tomasi method(Shi & Tomasi, 1994), we detected feature points, which are the ‘corners’ changing rapidly in intensity in pulmonary vascular structures in the reference image. We tracked the feature points from the reference image to the paired acquired image using Lucas Kanade optical flow analysis, the algorithm estimating the motion vector between frames(Lucas & Kanade, 1981). These feature point pairs were then used to calculate a homography matrix, which defines the geometric transformation of the paired images. This perspective transformation matrix was used for motion correction. Furthermore, to reduce the motion caused by respiratory cycle of the ventilator, we averaged frames corresponding to 1 respiratory cycle.

Next to determine the effectiveness of the stabilization method, we compared CASTii with current imaging methods, where only respiratory cycle averaging is used(Looney et al., 2011). Here we used Catchup mice(Hasenberg et al., 2015) in which PMNs genetically express the fluorescent protein tdTomato. We also injected i.v. fluorescent anti-Ly6G and anti-CD31 antibodies in these mice to visualize intravascular PMNs and endothelial cells, respectively. Pulmonary capillary endothelial cells were used to estimate lung motion and calculate homography matrices, and the transform matrices were then applied to all color channels for motion correction. Upon acquiring lung PMN images after i.t. instillation of LPS (i.t. LPS), we observed that the images acquired by conventional methods were blurry such that PMN and lung structures could not be clearly resolved (Figure 1b and supplementary movie 3). However, using CASTii we achieved markedly precise motion corrections and resolved fine PMNs and lung structural details over time (Figure 1b and supplementary movie 3). The stabilized images by CASTii also clearly resolved the endothelial monolayer lining of lung microvessels as well as PMNs in intracapillary and extracapillary spaces, where the double positive PMN showing tdTomato expression and anti-Ly6G antibody staining was localized in microvessels whereas single positive PMN with tdTomato expression was present only outside vessels (Figure 1c, d). These results show the ability to visualize PMN in lung micro-compartments (Figure 1c, d and supplementary figure 2f). Sub-cellular structures like PMN cell membrane were also visualized using the anti-Ly6G antibody (Figure 1c, d).

To quantitatively evaluate the precision of image registration, we next assessed lung motion displacement and deformation using CASTii. Without stabilization, we observed peak-to-peak displacement of 60 μm and deformation of 15 μm, whereas these parameters were markedly reduced by CASTii (Supplementary figure 2d). The motion displacement and deformation plots obtained using CASTii were 2D Gaussian fitted, and the image registration precision was calculated (Supplementary figure 2e). Since the standard deviation from 2D Gaussian fitting using our method was 0.15-0.19 μm (SE=0.002 μm), we were able to achieve image registration precision lower than optical resolution (theoretically 370 nm). We also statistically evaluated those parameters in the movies shown in this paper. Overall, the results showed marked improvement of motion displacement (peak-to-peak) by 64.5 times and deformation (peak-to-peak) by 12.8 times with sub-micron precision (Figure 1e and supplementary movie 4).

The impact of accurate image registration was next quantitatively assessed. The efficiency of cell analysis was evaluated as the efficiency of cell detection using an automated cell counting algorithm (see Materials and Methods). Only 5% of movies showed >90% cell detection efficiency with conventional methods, while almost all movies had >90% cell detection efficiency with CASTii, achieving high throughput cell analytical efficiency with greater than 20-fold improvement by CASTii (Figure 1f and supplementary movie 4). Cell localization was evaluated as the relationship between PMN localization (i.e., in/outside of vessel) and their reactivity to intravascular antibodies. >90 % intracapillary PMNs showed significantly high Ly6G antibody signal compared to extracapillary PMNs (Figure 1g), indicating that accurate image registration enabled analysis of compartmentalized PMN populations.

We also compared the present computer vision image registration algorithm with the existing algorithms (Supplementary table1 and supplementary movie 5). The diffeomorphic demons image registration algorithm, as an example of the existing intensity-based methods, was better than the feature-based methods in terms of motion correction accuracy, but its computational run time was 21 times slower (Supplementary table1 and supplementary movie 5). We also compared the transform algorithms (Supplementary table1 and supplementary movie 5). Translational transform failed to correct for the distortions caused by lung tissue motion, while perspective transformation achieved similar motion correction accuracy to non-rigid transformation as well as diffeomorphic demons. Based on these results, we adopted perspective transform into our image registration algorithms which achieved the fastest precise motion correction among these algorithms.

### Defective PMN transit through lung vasculature induced by endotoxemia

We assessed PMN localization and morphology in mouse lungs in the resting state and after endotoxin challenge induced by either i.p. or i.t. LPS. We observed PMN in lung microvessels in the basal state, 15 cells/298 μm square FOV (Figure 2a, b). PMN accumulation in lungs observed under basal conditions was not due to experimental artifact such as the surgery needed to insert the optical window, as confirmed by post-mortem imaging of lungs from mice injected with cell specific antibodies (Supplementary figure 4a-d). PMN number in lungs markedly increased over time after either i.t. or i.p. LPS injection (Figure 2a, b **and** supplementary figure 5a, b).

**Figure 2.**
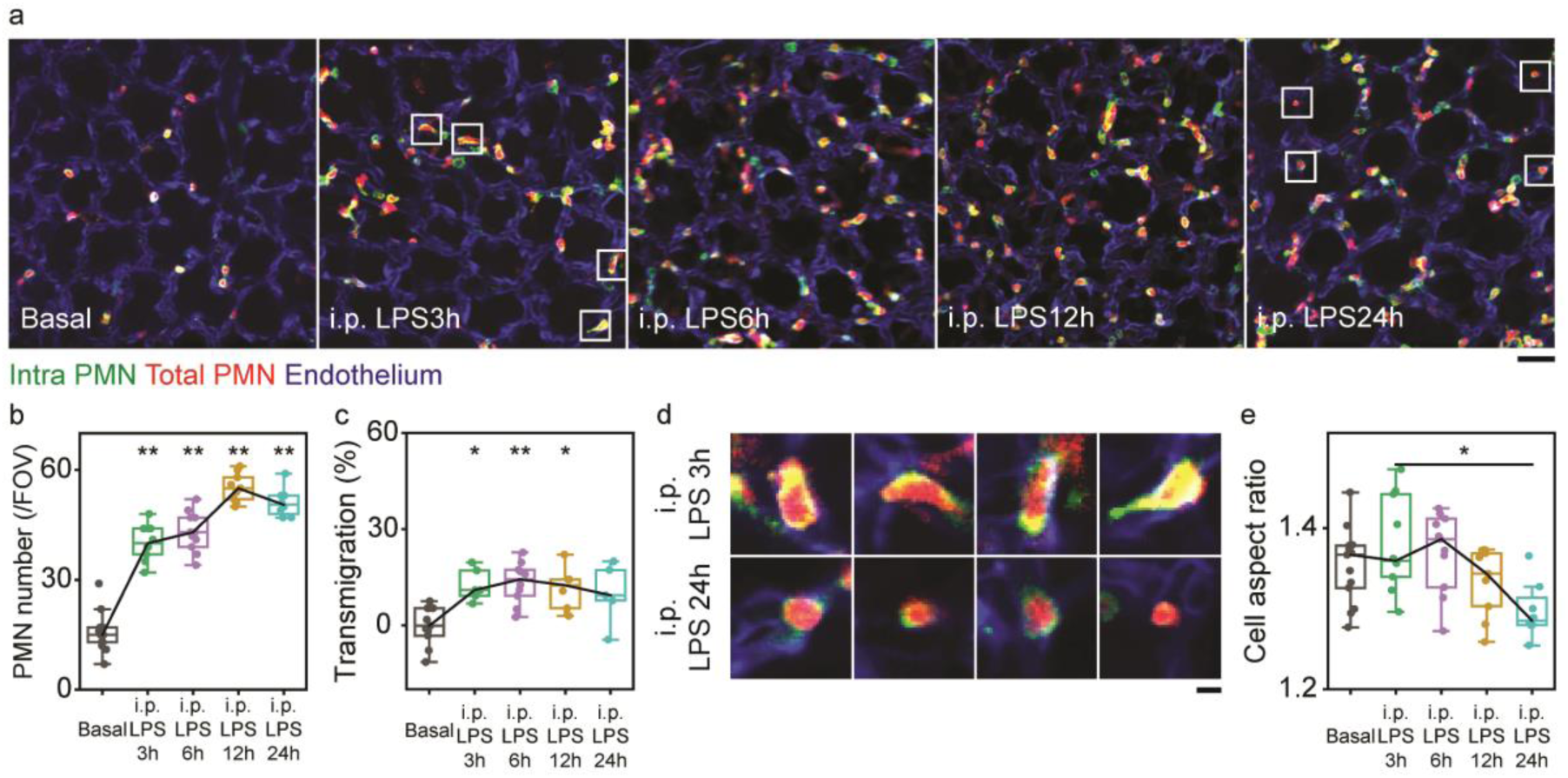
PMN morphological changes and lung localization in endotoxemia. **a.** Two-photon fluorescence images of PMNs and lung microvessels at different time points after i.p. LPS challenge. 10-seconds movies were averaged for each image. Scale bar; 40 μm. **b.** Quantitative analysis of PMN number per FOV; n=13 (FOV) for basal, n=10 (FOV) for i.p. 3h LPS, n=11 (FOV) for i.p. 6h LPS, n=9 (FOV) for i.p. 12h LPS, n=8 (FOV) for i.p. 24h LPS. **c.** Quantitative analysis of PMN transmigration; n=203 (cell) in 11 FOVs for basal, n=234 (cell), n=473 (cell) in 6 FOVs for i.p. 3h LPS, n=504 (cell) in 10 FOVs for i.p. 6h LPS, n=497 (cell) in 6 FOVs for i.p. 12h LPS, n=379 (cell) in 5 FOVs for i.p. 24h LPS. **d.** Magnified PMN morphology after i.p. 3h LPS and i.p. 24h LPS. Inset white boxes in a are magnified. Scale bar; 5 μm. **e.** Quantitative analysis of cell aspect ratio of PMN; n= 6092 (cells) in 13 FOVs for basal, n= 10841 (cell) in 8 FOVs for i.p. 3h LPS, n= 13784 (cell) in 10 FOVs for i.p. 6h LPS, n= 12277 (cell) in 8 FOVs for i.p. 12h LPS, n= 9243 (cell) in 8 FOVs for i.p. 24h LPS. Statistical analysis was performed using one-way ANOVA Tukey test for b, c and e; *P<0.05, **P<0.01. All two-photon images are single z slices.

We next quantitatively distinguished between intravascular and extravascular PMNs (Figure 1b-d **and** supplementary figure 2f **and** 5c) determined PMN transmigration across pulmonary microvessels (Figure 2c **and** supplementary figure 5d). Transmigration was defined as extravascular PMN number/total PMN number. We observed that almost all PMN were present in the intravascular space in the resting state with no evidence of tissue resident PMN; however, 25-30% of PMN were transmigrated at 4h after i.t. LPS whereas only 10-15 % PMN were observed at this time in extravascular space after i.p. LPS (Figure 2c and supplementary figure 5d).

The cell aspect ratio (defined as σ_large_/σ_small_) decreased over time after i.p. LPS (Figure 2d, e) but not after i.t. LPS (Supplementary figure 5e), indicating that PMN transitioned from migratory to non-migratory phenotype during endotoxemia but not in response to localized i.t. LPS. Interestingly, the cell aspect ratio of extravasated PMN had significantly higher values as compared with intravascular PMN (Supplementary figure 5g), indicating a distinct morphological change of PMN in the parenchymal tissue. In a control experiment with i.t. PBS and i.p. PBS and without surgery to insert optical windows, we observed no significant differences as compared to basal conditions (Supplementary figure 4a-d).

We next studied the behavior of circulating PMN in lungs by determining PMN dynamics at rest. PMN showed “catch and release” type of dynamics in the basal state (Figure 3a, b **and supplementary movie 6**); that is, PMN were briefly arrested in larger diameter microvessels (∼10 μm) at their point confluence with smaller diameter microvessels, and then squeezed into narrower ∼5 μm diameter vessels through PMN deformation (as reflected by increased cell aspect ratio) (Figure 3b-d). To study the kinetics of PMN in lung microvessels, we quantified the cell number/FOV over time using automatic cell counting based on the 2D Gaussian fitting algorithm(Yildiz et al., 2003), a method resistant to noise by using the surrounding image information. In the basal state, the change of cell number slowly fluctuated due to PMN turnover in FOV, indicating that PMN repeatedly entered into and exited from the FOV (Figure 3e). Autocorrelation analysis(Negwer et al., 2018; Pack et al., 2014) showed the mean dwell time (duration of PMN residence in FOV) value was 36 seconds with single exponential fitting of autocorrelation curve in the basal state (Figure 3f).

**Figure 3.**
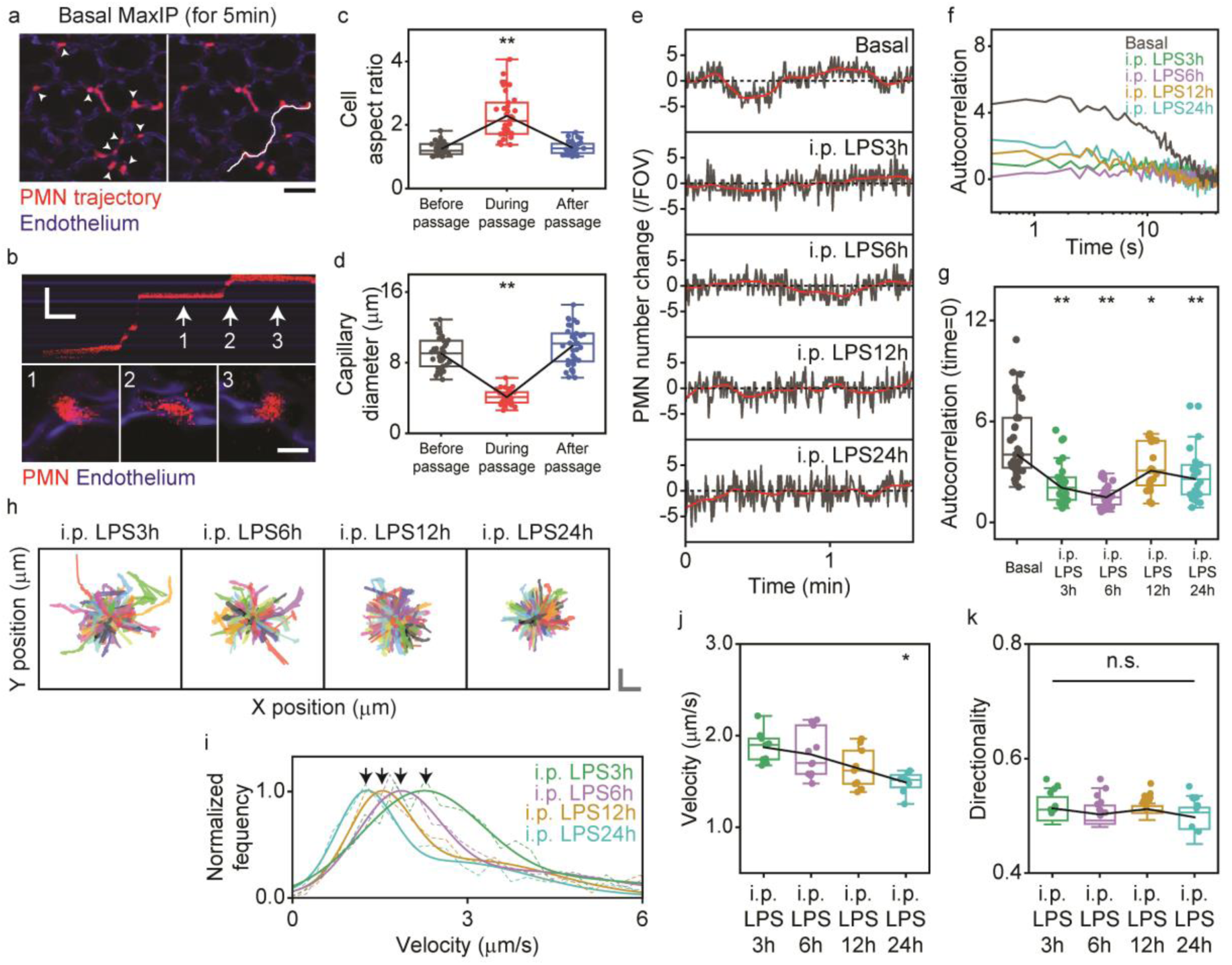
PMN dynamics at rest and in response to endotoxemia in respiring lungs. **a.** PMN trajectory at baseline. Maximum intensity projection (MaxIP) over 5 min is shown as well as PMN trajectory in white (right panel); lung microvessels in which transiently trapped PMNs are labeled (arrow heads). Scale bar; 40 μm. **b.** Kymograph showing PMN passage in lung microvessels. The white trajectory line was examined via kymograph to demonstrate the “catch and release” kinetics of PMNs. Scale bar; 50 μm (vertical) and 10 s (horizontal). Morphology of PMNs at 3 time points (1,2,3) within the vessel, as indicated by white arrow, is shown at the bottom. Scale bar; 10 μm. **c.** Quantitative analysis of cell aspect ratio during PMN passage; n=35 (cell) in 12 FOVs. **d.** Quantitative analysis of diameter of lung microvessels during PMN passage. **e.** Alterations in total number of PMNs within a FO over time recorded under a variety of conditions. **f.** Autocorrelation analysis of PMN number fluctuations at baseline and after i.p. LPS challenge. PMN number fluctuations over 2.1 min are used for autocorrelation analysis. **g.** Quantification of autocorrelation values at time lag of 0; n=33 (data set) in 11 FOVs for Basal, n=30 (data set) in 11 FOVs for i.p. 3h LPS, n=27 (data set) in 10 FOVs for i.p. 6h LPS, n=27 (data set) in 9 FOVs for i.p. 12h LPS, n=24 (data set) in 8 FOV for i.p. 24h LPS. **h.** Trajectory of PMN migration during endotoxemia induced by i.p. LPS. Scale bar; 10 μm (vertical) and 10 μm (horizontal); n=304 (cell) for i.p. 3h LPS, n=329 (cell) for i.p. 6h LPS, n=384 (cell) for i.p. 12h LPS, and n=307 (cell) for i.p. 24h LPS. **i.** Changes in velocity distribution over time after i.p. LPS. Arrowheads indicate the first peak obtained by double Gaussian fitting; n=898 (cell) for i.p. 3h LPS, n=692 (cell) for i.p. 6h LPS, n=805 (cell) for i.p. 12h LPS, and n=1171 (cell) for i.p. 24h LPS. **j.** Quantitative analysis of PMN velocity at different times after i.p. LPS. **k.** Quantitative analysis of directionality of PMN motility at different times after i.p. LPS; n=10841 (cell) in 8 FOVs for i.p. 3h LPS, n= 13784 (cell) in 10 FOVs for i.p. 6h LPS, n= 12277 (cell) in 8 FOVs for i.p. 12h LPS, n= 9243 (cell) in 8 FOVs for i.p. 24h LPS. Statistical analysis was performed using one-way repeated measures ANOVA Tukey test in c and d, and one-way ANOVA Tukey test in g, j, k; *P<0.05, **P<0.01. All two-photon images are single z slices.

We next followed PMN dynamics post i.t. and i.p LPS and analyzed the results in a similar manner (Figure 3e, f **and** supplementary figure 6a, b **and supplementary movie 7, 8**). We observed that i.t. LPS induced persistent slow fluctuations of PMN numbers (Supplementary figure 6a); also the autocorrelation curves and autocorrelation values at time=0 were similar in shape and values to the basal state (Supplementary figure 6b, c). Following i.p. LPS, however, PMN number fluctuations were reduced compared to the basal state (Figure 3e) and autocorrelation curves were shifted to lower values and autocorrelation values at time=0 were also decreased (Figure 3f, g). These data show that PMN circulating dynamics are maintained post-i.t. LPS whereas PMN showed markedly defective motility in lung microvessels post-i.p. LPS, as compared to controls (Supplementary figure 4e).

We also determined PMN migration dynamics in lungs following endotoxin challenge using the automated cell tracking method (Figure 3h **and** supplementary figure 6d **and supplementary movie 9**). The peak of PMN velocity distributions calculated from PMN migration trajectories were significantly decreased after both i.t. LPS (from 1.8 μm/s to 1.5 μm/s) and i.p. LPS (from 1.9 μm/s to 1.5 μm/s) (Figure 3i, j **and** supplementary figure 6e, f). While the directionality of PMN migration was reduced post i.t. LPS (Supplementary figure 6g), it did not significantly change post i.p. LPS (Figure 3k). Thus, changes in migration velocity and directionality were prominent with i.t. LPS as compared to i.p. LPS which significantly restricted the movement of PMN in microvessels. These findings show that the route of LPS administration differentially alters PMN trafficking, and as a result may have a different role in mediating inflammatory lung injury(Matute-Bello, Frevert, & Martin, 2008).

We next determined PMN migration velocity and directionality of the extravasated PMN as compared with intravascular PMN (Supplementary figure 5f, h). Both parameters were significantly reduced in extravascular PMN indicating that PMN transmigration into tissue decreases both their velocity and directionality, and PMN motility was entirely different in blood microvsessels as compared to extravascular tissue.

### Determinative role of PMN margination in lung microvessels in facilitating PMN recruitment into lung tissue

PMN form a large marginated pool in lung microvessels (Cooper, Bizios, & Malik, 1985; Doerschuk et al., 1987; Hogg, 1987; Kolaczkowska & Kubes, 2013; Peters, 1997). Using CASTii, we found that margination was the result of continuous cycles of “catch and release” of PMN as they traversed in microvessels as the result of cell deformation (Figure 3a-d). As the lung PMN marginated pool size can be altered by pulmonary vasoconstriction(Bierman et al., 1952), we assessed the effects of norepinephrine-induced pulmonary vasoconstriction. Intravenous injection of norepinephrine significantly and rapidly decreased PMN number in the pool while injection of control PBS had no effect (Figure 4a-c **and supplementary movie 10**). PBS or norepinephrine did not affect capillary diameters (Supplementary figure 7a) consistent with norepineprine-induced vasoconstriction occurring in small pulmonary arteries (Dawson, Bronikowski, Linehan, & Hakim, 1983; Grimm, Dawson, Hakim, & Linehan, 1978). PMN pool size was defined as stationary PMN/total PMN in lung vessels, where total PMN consisted of stationary PMN and dynamic PMN; the value is expressed as μ-2σ/μ+2σ from mean μ and standard deviation σ of PMN cell number fluctuation (Figure 4b). PMN pool size was reduced by norepinephrine from 75% to 67% (Figure 4d). Autocorrelation analysis^22,23^ (used to evaluate the effects of norepinephrine on PMN dwell time) also showed that the autocorrelation curve did not change with PBS whereas it shifted to the left with norepinephrine, indicating reduction of the dwell time (Figure 4e, f). Measurements of dwell times by single exponential fitting of autocorrelation curves showed that the dwell time after norepinephrine decreased markedly from 36 to 7 seconds while no change was seen with PBS (Figure 4g), demonstrating marked plasticity of lung’s marginated PMNs in response to pulmonary hemodynamic changes.

**Figure 4.**
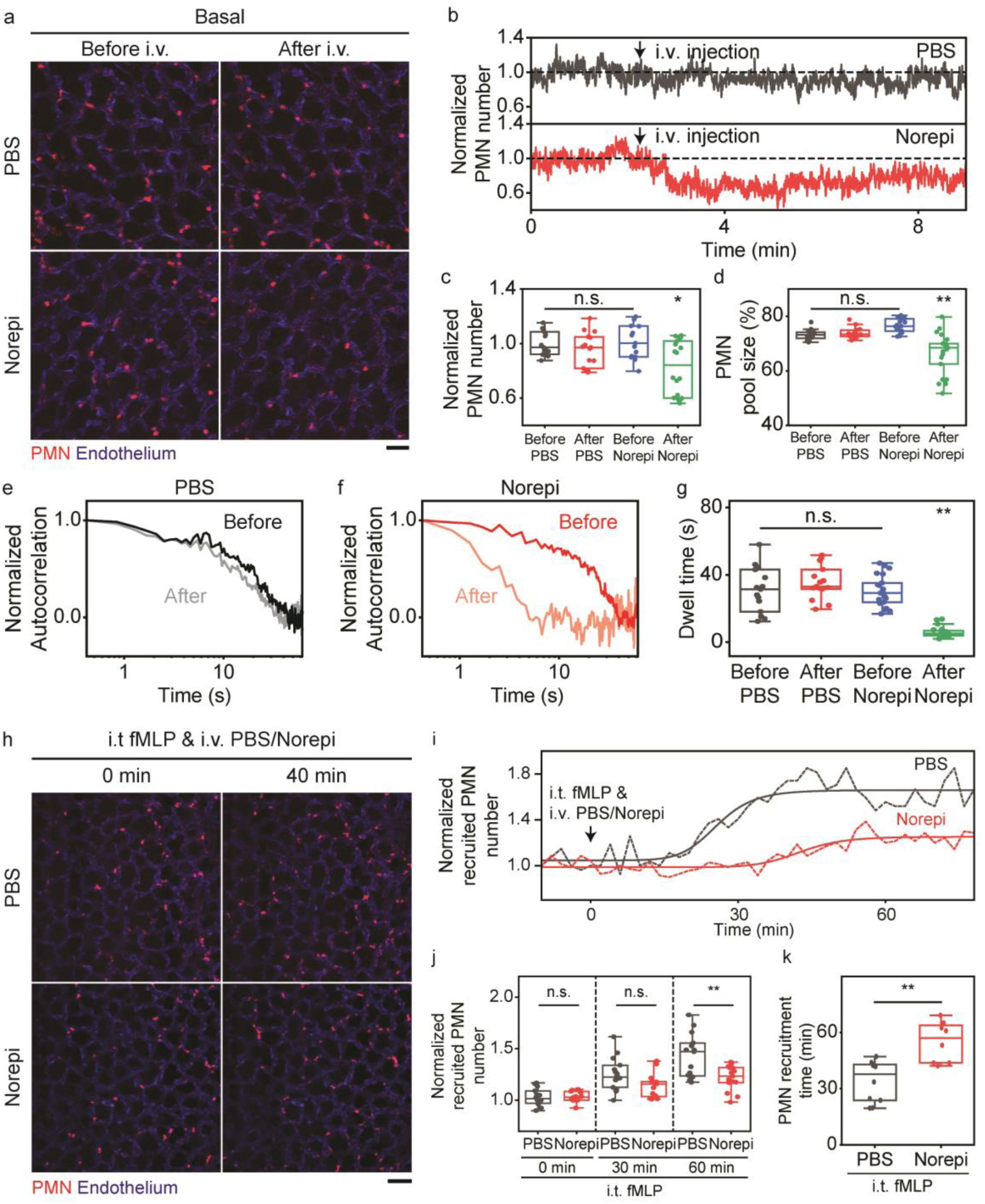
Dynamics of marginated PMN pool in lung vessels. **a.** Two-photon fluorescence images of PMNs and pulmonary microvessels before and after i.v. injection of PBS or norepinephrine (Norepi). Scale bar; 40 μm. **b.** PMN number changes in response to i.v. injection of PBS or norepinephrine. **c. d.** Quantitative analysis of PMN number (c) and PMN pool size (d) before andafter i.v. injection of PBS/norepinephrine (Norepi). PMN pool size is defined as ratio of stationary and total PMN (see text); n=15 (data set) in 5 FOVs for before PBS, n=15 (data set) in 5 FOVs for after PBS, n=15 (data set) in 7 FOVs for before Norepi, n=18 (data set) in 7 FOVs for after Norepi. **e. f.** Autocorrelation analysis of PMN number fluctuations before abd after i.v. injection of PBS (e) or norepinephrine (Norepi) (f). **g.** Quantitative analysis of dwell time from autocorrelation curves; n=15 (data set) in 5 FOVs for before PBS, n=15 (data set) in 5 FOVs for after PBS, n=21 (data set) in 7 FOVs for before Norepi, and n=21 (data set) in 7 FOVs for after Norepi. **h.** Two-photon fluorescence images of PMNs and pulmonary microvessels after i.t. inhalation of fMLP and i.v. injection of PBS/norepinephrine (Norepi). Scale bar; 60 μm. **i.** Recruitment of PMNs in response to i.t. inhalation of fMLP and i.v. injection of PBS or norepinephrine (Norepi). Recruited PMNs are defined as PMNs remaining in FOV for more than 4 min. **j.** Quantitative analysis of PMN recruitment after i.t. inhalation of fMLP and i.v. injection of PBS or norepinephrine (Norepi). n=15 (FOVs) in i.t. fMLP i.v. PBS or Norepi. **k.** Quantitative analysis of PMN recruitment time. Recruitment time was determined by using logistics fitting only when the raising phase appeared; n= 10 (FOVs) in i.t. fMLP i.v. PBS and n=8 (FOVs) in i.t. fMLP i.v. Norepi. Statistical analysis was performed using one-way ANOVA Tukey test in c and d and g and j and two-sample t-test in k; *P<0.05, **P<0.01. All two-photon images are single z slices.

We next tested the hypothesis PMN marginated pool normally present in lung microvessels facilitates PMN recruitment into lung tissue in an on-need basis in response to airway infection and release of chemoattractants in the airspace. Studies were made by instilling the chemoattractant fMLP into airways. We observed rapid lung tissue PMN accumulation in response to i.t. fMLP whereas PMN accumulation was significantly attenuated and reduced following reduction in the marginated pool induced by i.v. norepinephrine (Figure 4h-k **and supplementary movie 11**). PMN accumulation was not seen in control experiments without airway application of fMLP (Supplementary figure 7b, c). The recruited PMN numbers, defined as the number of PMN remaining in the same FOVs for at least 4 min, were significantly different between i.v. PBS and norepinephrine (Figure 4i, j). The PMN accumulation times quantified by logistics function fitting were also significantly altered by reductions in PMN dwell times with norepinephrine (Figure 4i, k). These data together show the essential role of normal PMN margination in microvessels in promoting PMN recruitment efficiency into lung tissue.

### Defective transmigration of lung vascular sequestered PMN impairs bacterial phagocytosis in airspace

Based on the time-dependent reduction in PMN velocity and increased PMN sequestration in lung microvessels at 24h following induction of endotoxemia post i.p. LPS challenge (Figure 3h-j), we addressed the migratory potential of these PMN; i.e., ability of long-term lung vascular sequestered PMNs to transmigrate into lungs. These studies were made by establishing a chemoattractant gradient by instilling fMLP directly into airways. We observed that PMN motility increased after i.t. fMLP administered at 3h post i.p. LPS, whereas surprisingly no change was seen after i.t. fMLP gradient was established at 24h post i.p. LPS (Figure 5a, b **and supplementary movie 12**). Thus, the long period of endotoxemia had paralysed the ability of PMN to transmigrate. Analysis of PMN velocity, cell aspect ratio, and directionality of PMN migration showed significant changes after i.t. fMLP between the 3h and 24h post i.p LPS conditions (Figure 5c). In control experiments, these parameters showed no significant changes after i.t. PBS (Supplementary figure 7d). We observed PMN transmigration in response to fMLP gradient in the 3h i.p. LPS group whereas PMN transmigration was severely impaired in the 24h i.p. LPS group (Figure 5d). PMN transmigration increased by 2.5 times after applying the fMLP gradient in the i.p. LPS3h condition whereas PMN transmigration did not increase after i.t. PBS control (Figure 5e). After 24h of i.p. LPS, we observed failure of PMN to transmigrate in response to the same fMLP gradient (Figure 5e). The transmigrated PMNs at post 3h LPS demonstrated slower migration velocities, higher cell aspect ratios, and reduced directionality in the tissue consistent with previous observation (Supplementary figure 5f-h) as compared to intracapillary PMN (Figure 5f). These results showed that prolonged 24h lung vascular PMN sequestration induced by endotoxemia interfered with the ability of PMN to transmigrate into airspace.

**Figure 5.**
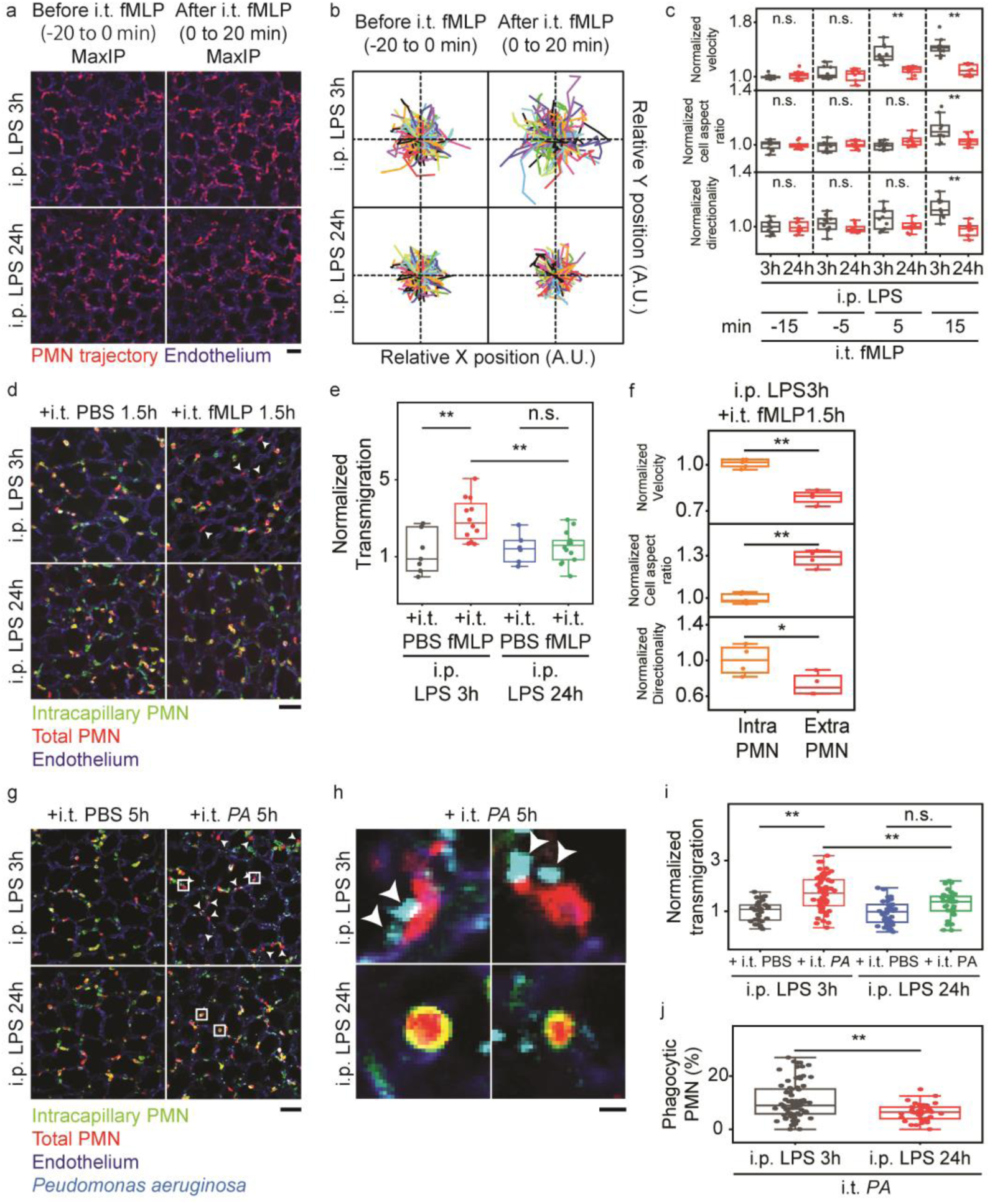
Time dependent defects in PMN transmigration and phagocytosis in lungs post-endotoxemia. **a.** Two-photon fluorescent images of PMN trajectory and pulmonary microvessels before and after i.t. fMLP in mice challenged i.p. LPS for 3h or 24h. Results show maximum intensity projection over 20 min as PMN trajectory. Scale bar; 40 μm. **b.** Trajectory of PMN migration before and after i.t. fMLP in i.p. 3h vs. 24h LPS in mice; n=178 (cell) for before i.t. fMLP at i.p. 3h LPS condition, n=216 (cell) for after i.t. fMLP at i.p. 3h LPS condition, n=261 (cell) for before i.t. fMLP at i.p. 24h LPS condition, and 240 (cell) for after i.t. fMLP at i.p. 24h LPS condition. **c.** Quantitative PMN velocity, cell aspect ratio, and directionality before and after i.t. fMLP at i.p. 3h or 24h LPS; n=900-1200 (cell) in 10 FOVs for i.p. 3h LPS, n=1300-1800 (cell) in 10 FOVs for i.p. 24h LPS. **d.** Two-photon fluorescent images of PMNs and pulmonary microvessels after 1.5 h of i.t. PBS or fMLP at i.p. 3h or 24h LPS. Arrows indicate transmigrated PMNs. Scale bar; 40 μm. **e.** Quantitative analysis of transmigration of PMN after 1.5 h of i.t. PBS or fMLP at i.p. 3h or 24h LPS; n=432 (cell) in 7 FOVs for i.t. PBS at i.p. 3h LPS, n=697 (cell) in 12 FOVs for i.t. fMLP at i.p. 3h LPS, n=132 (cell) in 6 FOVs for i.t. PBS at i.p. 24h LPS, n=582 (cell) in 14 FOVs for i.t. fMLP at i.p. 24h LPS. **f.** Changes in velocity, cell aspect ratio and directionality of PMN migration between intravascular and extravascular compartments after 1.5 h of i.t. fMLP at i.p. 3h LPS; n=426 (cell) in 5 (data set) and 20 FOVs for intravascular PMN (Velocity), n=376 (cell) in 4 (data set) and 20 FOVs for extravascular PMN (Velocity), n=479 (cell) in 5 (data set) and 20 FOVs for intravascular PMN (Cell aspect ratio), n=331 (cell) in 4 (data set) and 20 FOVs for extravascular PMN (Cell aspect ratio), n=255 (cell) in 4 (data set) and 20 FOVs for intravascular PMN (Directionality), n=281 (cell) in 4 (data set) and 20 FOVs for extravascular PMN (Directionality). **g.** Two-photon fluorescent images of PMNs and pulmonary microvessels after instillation of *Pseudomonas aeruginosa* (*PA*) for 5 h of i.t. vs. i.t. PBS control post challenge with i.p. 3h or 24h LPS. Arrows indicate transmigrated PMNs. Scale bar; 40 μm. **h.** Magnified *PA*-phagocytosing PMN after 5 h of i.t. *PA* in i.p. 3h LPS mice. Inset white boxes in g are magnified. Scale bar; 4 μm. **i.** Quantitative analysis of transmigration of PMN after 5h of i.t. PBS or *PA* in response to i.p. 3h or 24h LPS; n=1932 (cell) in 30 FOVs for i.t. PBS at i.p. 3h LPS, n=3759 (cell) in 59 FOVs for i.t. *PA* at i.p. 3h LPS, n=2700 (cell) in 36 FOVs for i.t. PBS at i.p. 24h LPS, n=2059 (cell) in 31 FOVs for i.t. *PA* at i.p. 24h LPS. **j.** Changes in phagocytosis by PMN of PA after 5 h of i.t. *PA* in mice challenged with i.p. 3h or 24h LPS; n=4171 (cell) in 77 FOVs and n=2941 (cell) in 42 FOVs. Statistical analysis was performed using one-way ANOVA Tukey test in c, e and i and two-sample t-test in f and j; *P<0.05, **P<0.01. All two-photon images are single z slices.

Next, to use a translationally relevant model of lung injury, we challenged the mice with live *Pseudomonas aeruginosa* (*PA*) bacteria i.t. to address the bacterial killing function of the transmigrated PMNs. In *PA* i.t. infected lungs, PMN failed to transmigrate at 24h after i.p. LPS challenge but not at 3h (Figure 5g). The transmigration in response to *PA* was increased by 1.3 fold after i.p. LPS 3h when compared with 24h (Figure 5i).

These observations were confirmed using whole lung analysis of PMNs (Supplementary figure 8a, b). We also observed that the transmigrated PMNs effectively phagocytosed *PA* within the airspace at 3h in contrast to transmigrated PMN residing in microvessels for 24h after i.p. LPS challenge (Figure 5h). Quantification of phagocytic activity (the number of PMN colocalized with *PA*)/(total PMN number in tissue) showed a significantly decrease from 11% (i.p. LPS 3h) to 6% (i.p. LPS 24h) (Figure 5j) indicating that extravasated PMNs present at 3h post LPS actively phagocytosed *PA* as compared to extravasated PMNs present at 24h post LPS. The difference of bacterial burden was detectable using the described transmigration analysis (Supplementary figure 8c). Lung injury with lung edema was also more severe 24h after i.p. LPS injection than after 3h of i.p. LPS (Supplementary figure 8d). These data demonstrate that defective PMN transmigration because of prolonged PMN sequestration in lung microvessels induced by prolonged endotoxemia resulted in impairment in PMN transmigration and *PA* phagocytic capacity.

## Discussion

Here we describe a novel stabilized intravital imaging method based on a computer vision image registration algorithm, CASTii. This method enabled visualization of fine lung structures such as microvessels and PMN traversing them and tissue-compartmentalized PMN trafficking in real-time with sub-micron image registration in normally respiring mouse lungs. We demonstrated (i) the method’s utility in intravital lung imaging in respiring lungs and ability to provide continuous high-resolution images that allowed high-throughput quantitative assessment of tissue-compartmentalized PMNs’ dynamics in lung microvessels and extravasated PMNs; (ii) for the first time via direct PMN visualization the functional role of the previously described PMN marginated pool in the lung vasculature(Cooper et al., 1985; Doerschuk et al., 1987; Hogg, 1987; Kolaczkowska & Kubes, 2013; Peters, 1997) in rapidly facilitating PMN migration into lung tissue as a requirement for efficient PMN bactericidal function, and (iii) failure of PMN transmigration during severe endotoxemia-induced PMN sequestration in microvessels, which induced defective phagocytosis of bacteria and contributed to impaired host-defense function of lungs.

We visualized and quantified the pooling of PMN in lung microvessels. We showed that the marginated pool constituted up to 75% of total PMNs in resting mouse lungs, a value in range of previous determinations(Cooper et al., 1985; Doerschuk et al., 1987; Hogg, 1987; Kolaczkowska & Kubes, 2013; Peters, 1997). There are however controversies about the variation of PMN pooling size (40-90 %) (Cooper et al., 1985; Doerschuk et al., 1987; Hogg, 1987; Kolaczkowska & Kubes, 2013; Peters, 1997). These differences may reflect architecture of the pulmonary microvessels among specieis and the PMN mechanics traversing the microvessels as well as methods of quantifying the PMN pool. Our approach involving direct visualization of neutrophils by intravital imaging complements prior methods which relied on radioactive measurements to characterize the size of the marginated pool as well as the transit time of neutrophils(Doerschuk et al., 1987; Summers et al., 2014). Another concern is that priming of PMN might affect PMN circulatory dynamics (Doerschuk et al., 1987; Summers et al., 2014); however, this possibility was excluded by imaging post-mortem mouse lungs in which the size of the pool was found to be similar values determined in live mice by the described *in vivo* imaging method. We demonstrated that the pool was the result of the dwell time of PMNs traversing the lung microvessels in which PMNs displayed repeated cycles of “catch and release” as the cells entered microvessels of different diameters. We observed that PMN reduced their cell size while traversing lung microvessels ∼5 μm in diameter and slowed significantly in larger diameter microvessels (∼10 μm). The rounding and squeezing of PMN in these vessels is consistent with the ability of PMNs to change shape and contract in the lung microcirculation on the basis of their actin cytoskeletal machinery(Sarantos et al., 2008).

We showed that the lung vascular marginated PMN pool was not a static feature of microvessels. PMN dwell times shortened in response to pulmonary arterial vasoconstriction induced by norepinephrine injection(Dawson et al., 1983; Grimm et al., 1978), and PMN pool size concomitantly decreased. This is in agreement with evidence that increased blood velocity shortened the dwell time of PMNs and decreased the pool size(Bierman et al., 1952). The question arises about the function of lung vascular marginated PMN pool. We demonstrated that the effectiveness of recruitment of PMNs into tissue was dependent on the pool of PMNs in lung microvessels. We observed that a larger PMN pool enabled rapid mobilization of PMNs into lung tissue in response to establishing a fLMP chemotactic gradient in airways. Thus, margination of PMNs in lung microvessels serves to efficiently recruit PMNs into lung tissue, a function essential for host-defense of lungs through PMN mediated bacterial killing(Ley et al., 2018). Importantly, recruitment of the marginated pool greatly facilitated the phagocytosis of live bacteria in the airspace.

High throughput compartmentalized intravital imaging of lung PMNs also revealed unique PMN dynamics in lungs in response to endotoxemia(Abraham, Carmody, Shenkar, & Arcaroli, 2000; Grommes & Soehnlein, 2011). While PMNs in the early phase of endotoxemia (3h after i.p. LPS) remained motile in lung microvessels and showed characteristic transmigration into lung tissue in response to a fMLP chemotactic gradient setup in the airspace and with instillation live *PA* bacteria in airways, protracted endotoxemia seen at 24h post LPS leading to persistent sequenstation of PMN severely impaired PMN transmigration into tissue. This PMN migratory paralysis significantly impaired phagocytosis of the airway-instilled *PA*. The increased number of PMNs migrating into tissue at 3h post LPS more effectively phagocytosed instilled *PA* as compared to the defective PMN transmigration into tissue at 24h post LPS coupled to impaired phagocytosis of live bacteria. Thus, the defective PMN transmigration seen in our studies helps to explain the impaired innate immune response in Acute Respiratory Distress Syndrome (ARDS) patients(Tavares-Murta et al., 2002) and difficulty in resolving pneumonia infection in septic patients with long-term PMN sequestration (Han & Mallampalli, 2015; Matthay et al., 2019).

Our results are consistent with a key pathogenic role of lung vessel sequestered PMNs in endotoxemia(Han & Mallampalli, 2015; Hellebrekers, Vrisekoop, & Koenderman, 2018; Matthay, Ware, & Zimmerman, 2012; Matthay et al., 2019; Ware & Matthay, 2000) since impaired transmigration prevented elimination of the bacterial pathogens. Our results also suggest that restoration of defective PMN transmigration phenotype may rescue the immune defects seen in patients with sepsis, pneumonia, and ARDS(Han & Mallampalli, 2015; Matthay et al., 2019). We also demonstrated that intravascular sequestration of PMNs during prolonged endotoxemia in lung microvessels due to hyper-adhesivity of the activated endothelium(Schmidt et al., 2012), markedly impaired PMN transmigration and bacterial killing. This might explain the intractability of severe inflammatory lung as in ARDS with preponderance of PMN sequestered in lung microvessels that are unable to transmigrate and inappropriately positioned in vessels where they induce vascular injury(Han & Mallampalli, 2015; Matthay et al., 2019).

Despite the usefulness of CASTii in imaging of lungs as described above, there are still possible limitations. One is that the lung motion may exceed more than half of the field of view, thus reducing the area of lung available for the imaging analysis. Z movement and deformation in the setting of a touchless preparation without a thoracic window also cannot be corrected. Instrumental active feed-back of tissue motion with real-time tracking could obviate this issue(Griffiths et al., 2020; Karagyozov, Mihovilovic Skanata, Lesar, & Gershow, 2018). While the perspective transformation using a homography matrix is well-suited for lung intravital imaging, it would not suffice in situations with local tissue deformations. Solutions such as introducing non-rigid image registration algorithms(Crum, Hartkens, & Hill, 2004) is an option to address tissue deformations as shown in Supplementary table1 and supplementary movie5. Another limitation relates to our use of a short-term experimental optical window that required opening of the pleural cavity prior to imaging. This limited the imaging duration to 3-4h in our studies; however, implanting permanent optical windows(Entenberg et al., 2018) could overcome this limitation. Another possible concern is that excitation laser distortions due to refraction between airspace and tissue may reduce image resolution in deep lung tissue. However, mprovements incorporating adaptive optics may correct the wave front corresponding distortion, and provide deeper lung tissue imaging(Ji, Milkie, & Betzig, 2010).

In summary, we describe a stabilized intravital imaging with computer-vision algorithm, CASTii, which corrects for lung motion artifacts at sub-micron precision and thus achieves high-throughput and lung compartmentalized PMN dynamic imaging. We defined a highly active patrolling function of PMNs in mouse lungs in the basal state and tissue recruitment of PMNs dependent on the size of vascular pool of marginated PMNs. We showed that the lung marginated PMN pool has a previously unrecognized role in rapidly mobilizing PMN into tissue required for efficient phagocytosis and bacterial killing. These observations support an essential role of lung microvessel marginated PMNs in serving an important host-defense function through their ability to rapidly mobilize into the airspace.

## Materials and Methods

### Mice

Mice were housed and bred under specific pathogen-free conditions at the University of Illinois at Chicago (UIC) Animal Care Facility and all experiments conformed to ethical principles and guidelines approved by the UIC Institutional Animal Resources Center animal usage committee. Male and female mice between 8 and 16 weeks of age were used for experiments. C57BL/6N was purchased from Charles River Laboratories and Catchup mice were obtained from a Material Transfer Agreement with University Duisburg– Essen(Hasenberg et al., 2015). For intravital PMN imaging with both of endogenous labeling by tdTomato expression and exogenous antibody labelling, homogeneous Catchup mice, where the neutrophil-specific locus Ly6G was modulating with a knock-in allele expressing Cre recombinase and the fluorescent protein tdTomato and tdTomato with a Cre-activatable CAG promoter was inserted into the ROSA26 locus(Hasenberg et al., 2015), were cross-bred with C57BL/6N to generate heterogeneous Catchup mice. Heterogeneous Catchup mice were used for all experiments and analysis.

### Endotoxemia model of lung injury

Mice were injected with a single intraperitoneal dose of 5mg/ml LPS (Lipopolysaccharides from Escherichia coli O111: B4, L2630, Lot#026M4022V; Sigma-Aldrich) dissolved in PBS at 7.5-10 mg/kg body weight and lungs were imaged after 3, 6, 12, 24 hours. Mice were also inhaled aerosol of 1mg/ml LPS 5 ml (Lipopolysaccharides from Escherichia coli O55: B5, L2880, Lot#028M4094V; Sigma-Aldrich) dissolved in PBS or 0.8 mg/ml fMLP 2.5 ml (N-Formyl-Met-Leu-Phe, F3506, Lot#MKCC5355; Sigma-Aldrich), where 2 mg fMLP was dissolved in 50 μl DMSO and then in 2.5 ml PBS, using nebulizer connected with inhalation box (Compressor nebulizer, MODEL: NE-C30; OMRON). Lungs were imaged after 2, 4 hours for LPS and after 1.5 hours for fMLP inhalation. *Pseudomonas aeruginosa* (*PA*) constitutively expressing GFP (GFP-PA01) was prepared as previously described^(Joshi et al., 2020)^ and 1 x 10^7^ CFU in 40 µl of PBS was intratracheally sprayed using micro aerosol spray (Liquid PenWu Device, BJ-PW-M; Biojane). Lungs were imaged after 5 hours of *PA* solution injection. Biosafety level 2 room was used for all the experiments of *PA*, instructions and precautions were followed as directed by the safety rule of UIC Biologic Resources Labratory.

### Whole lung analysis

Lung single cell suspensions were prepared as described(Joshi et al., 2020). Briefly, lung tissues were minced and enzymatically digested with 1 mg/mL collagenase A (Roche) for 50 min at 37°C. Digested tissue was forced through metal canula and passed through a 75-mm nylon filter to obtain single-cell suspensions. The red blood cells were lysed using lysis buffer and cell suspensions were washed with FACS buffer. Samples were analyzed using CytoFLEX S Flow Cytometer (Beckman Coulter) and data were analyzed by using Kaluza Analysis software (Beckman Coulter). Viable bacterial burdens in the lung single cell suspensions were determined by plating serial dilutions onto *Pseudomonas* isolation agar (LB), followed by incubation at 37°C overnight. Excised lungs were used to determine the lung wet/dry weight ratio as described previously(Joshi et al., 2020). First, wet-weight was taken thereafter lung were kept for complete dry in the oven at 55-60°C overnight and dry lung weight measured followed by lung wet/dry weight ratio was determined to quantify lung edema.

### Lung intravital imaging of PMNs and microvascular structures

Mice were injected with ketamine (10 mg/ml) and xylazine (2.5 mg/ml) at 40-80 mg/kg body weight (for ketamine) and 10-20 mg/kg body weight (for xylazine). Intravenous injection via jugular vein of Brilliant Violet 421-labeled Ly6G antibody (25 μg/mice) (1A8, Biolegend), SeTau647(SETA BioMedicals)-labeled CD31 antibody (25 μg/mice) (390, Biolegend) and Fluorescein-Dextran (250 μg/mice) (D1823, Thermofisher Scientific) were performed to stain PMNs and lung microvascular structures, respectively, before surgery. Surgical methods for gaining access to the lung and the design of an optical window are based on Looney et al.(Looney et al., 2011). A resonance-scanning two photon microscope (Ultima Multiphoton Microscopes, Bruker Fluorescence Microscopy, Middleton WI; formerly Prairie Technologies) with an Olympus XLUMPlanFL N 20x (NA 1.00) and Immersion oil (Immersol W (2010); Carl Zeiss) were employed to collect multi-color images (Dichroic mirror; 775 long pass filter (775 LP; Bruker), IR blocking filter; 770 short pass filter (770 SP; Bruker), Emission filter; 460/50 nm for Brilliant Violet 421 (FF01-450/70-25; Semrock), 525/50 nm for Fluorescein (Bruker), 595/60 nm for tdTomato (Bruker) and 708/75 nm for SeTau647 (FF01-708/75-25; Semrock)) with 960 nm excitation (Laser power: 9.1-18.8 mW/cm^2^ at back aperture of objective) at video rate. For long-term lung intravital imaging, movies with z stacks (25 μm depths at 5 μm step) were acquired and the z plane images with correct focus was extracted by cross-correlation analysis. During imaging we applied 13.6-20.4 cmH_2_O of suction to gently immobilize the lung using Vacuum Regulators (AMVEX), where the optical window with a coverslip was sealed with high vacuum silicone grease (DOW CORNING; Sigma Aldorich) to avoid air leak and keep stable suction, and mice were ventilated using Mouse Vent G500 (Kent Scientific) with a tidal volume of 10μl air per gram of mouse weight or a target inspiratory pressure of 14-15 cmH_2_O when nebulizer unit (Aeroneb Lab Nebulizer Unit, small VMD; Kent Scientific) was connected, a respiratory rate of 138 breaths per minute, and a positive-end expiratory pressure of 3 cm H_2_O and 0.5-1.5% Isoflurane was delivered continuously to maintain anesthesia via ventilator and heart rate of mice was monitored by using ECG/EKG analysis system with needle electrode (ADInstruments). Objectives, immersion oil, mice stage and lung optical windows were kept at 27°C using a heater to avoid thermal drift and to stably maintain lung physiological condition. Imaging data was generated using Bruker Ultima In Vivo Multiphoton Microscopy System at the Fluorescence Imaging Core at Research Resources Center at University of Illinois at Chicago.

### Drug treatment during lung intravital imaging

Catheter was inserted into jugular vein and sutured into place using Micro Cannulation System (Braintree Scientific). 10 μg/ml 150 μl or 30 μg/ml 50 μl norepinephrine ((-)-Norepinephrine, A7257, Lot#SLBW7427; Sigma-Aldrich), where 10 mg norepinephrine was dissolved in 200 μl 0.5 M HCl and then in 10 ml PBS for 1 mg/ml norepinephrine solution, was intravenously injected via catheter during lung intravital imaging. 1 mg/ml fMLP 1 ml or 0.5 ml was intrathecally inhaled via ventilator with nebulizer unit during lung intravital imaging.

### Computer vision image registration algorithm of CASTii

The code of a computer vision image registration algorithm of CASTii is written in LabVIEW with Python integration kit (National Instruments) and Enthought Canopy (Enthought).

#### Algorithm: Feature point detection

Reference image is first smoothed via a gaussian blurring filter (Kernel; 27 pixels) to remove pixilation artifacts while preserving tissue structural information. Feature points are detected in the reference image with Shi and Tomasi method (LabVIEW’s IMAQ corner Detector VI) (Kernel; 3, Pyramid level; 1), which detects ‘corners’ that change rapidly in intensity(Shi & Tomasi, 1994). This search is set to allow 500 feature points per image.

#### Algorithm: Pairwise motion tracking and correction

We then attempt to track the feature points from reference image to paired image using Lucas Kanade optical flow analysis (LabVIEW’s IMAQ Optical Flow (LKP) VI) (Pyramid level; 6, Window size; 30)(Lucas & Kanade, 1981). Prior to optical flow analysis, the intensity of the paired image is normalized to that of reference image to reduce tracking error. Normalized intensity at paired image at a given pixel I_np_ (x_i_, y_j_) are acquired by I_np_ (x_i_, y_j_)= σ_r_ (I_p_ (x_i_, y_i_)-μ_p_)/σ_p_+ μ_r_, where I_p_ (x_i_, y_j_) is intensity at paired image at a the pixel (i, j), σ_r_ and σ_p_ are standard deviation of reference and paired image intensity of all pixel, respectively, μ_r_ and μ_p_ are the overall mean intensity of reference and paired image, respectively. We also run the optical flow analysis in reverse from paired image to reference image, and discard feature point pairs that are calculated to be further than 10 pixels. These remaining feature point pairs are used to calculate a 3X3 homography matrix, which defines the geometric transformation applied to the paired image, through OpenCV’s PerspectiveTransform function, in order to compute the stabilized image.

#### Algorithm: Motion correction with multiple reference images

In lung intravital imaging, it is sometimes difficult to perform pairwise motion tracking against a single reference image due to excessive respiratory cycle motion and deformation that prevents tracking and focusing. To overcome the issue, we use a multiple reference approach. Master reference image is collected as the highest image intensity from first 14 frames of movie. 7 reference images from the master reference, corresponding to half respiratory cycle, are extracted and pairwise motion tracking is performed between images. Finally, homography matrices of 7 reference images against master reference image is calculated. A reference image is chosen as tracking error, which is the ratio of discard feature point pairs and all feature point pairs, become < 20% and homography matrices of all images against master reference image are calculated and used for motion correction. For multi-channel lung intravital imaging, a series of homography matrices of reference channel is also used for motion correction of other channels.

### Comparison of computer vision image registration algorithms with the existing intensity-based image registration

We used Matlab’s imregdemons function (Image Processing toolbox) as intensity-based image registration algorithm to compare with our algorithm. The lung imaging movies with 20 frames were stabilized with imregdemons function with described parameter set as shown in Supplementary table1. Remaining motions after stabilization were evaluated by optical flow analysis and standard deviation of the remaining motion displacement were quantified. To compare among transforms, we used OpenCV’s AffineTransform function as translation transform and Matlab’s fitgeotrans and imwarp function (Image Processing toolbox) as non-rigid transform with described parameter set as shown in Supplementary table1. Computational time per this paper was estimated based on movies with total 25,596,000 frames used for paper’s analysis. Additional color channel was assumed to be half computational time.

### Quantitative image analysis

More than 3 mice data were used for statistical analyses.

#### Lung motion analysis

Lung motion was acquired at video-rate and motion vectors were estimated by optical flow analysis. The motion displacement vector v_di_ between from frame i and i+1 was expressed by v_di (i, i+1),j_ = (x_i+1,j_-x_i,j_, y_i+1,j_-y_i,j_), and the deformation vector v_de_ was expressed by v_de (i, i+1),j_ = ((x_i+1,j_-x_i,j_)-Σ_j_(x_i+1,j_-x_i,j_)/k, (y_i+1,j_-y_i,j_) -Σ_j_(y_i+1,j_-y_i,j_)/k), where x_i,j_ and y_i,j_ are the coordinates of feature points at frame i and paired feature point j and paired feature point number k. The motion displacement and deformation position were plotted as Σ_i_ v_di(i, i+1),j_ and Σ_i_ v_de(i, i+1),j_ along time frame i and the distance of peak-to-peak of displacement and deformation was calculated.

#### PMN dynamics analysis

CASTii-stabilized movies were directed to an automated cell counting and tracking software, G-count (G-angstrom), which is based on 2D gaussian fitting algorithm(Yildiz et al., 2003). In brief, a region of interest (ROI) was raster-scanned over each frame to identify cells based on whether 2D gaussian fitting could converge. By repeating this process over movie, we acquired cell number per FOV and quantitative information of candidate cells in the movie, including XY position, XY spreading by XY standard deviation and fluorescent intensity. Using the information, cell motions were tracked when the XY position in current frame was within a ROI from the previous frame. Automatic tracking allows us to acquire quantitative dynamics information such as cell trajectory, velocity and directionality(Suraneni et al., 2012). These data were further analyzed by customized LabVIEW programs. Transmigration occurrences were calculated by transmigrated PMN number (tdTomato positive cell-Ly6G positive cell number) / total PMN (tdTomato positive cell) number. Cell aspect ratios were calculated by σ_large_/σ_small_, where σ_large_ was larger standard deviation and σ_small_ was smaller standard deviation out of two standard deviation parameters obtained from 2D gaussian fitting. PMN number and transmigration occurrence were quantified by using averaged images of 10-seconds movies. Cell aspect ratios were quantified using 5-minuites movies. Path lengths of cell migration, p, was calculated by p=Σ_i=1∼j-1_((x_i+1_-x_i_)^2^+(y_i+1_-y_i_)^2^)^1/2^ and displacement of cell migration, d, was calculated by d= (x_j_-x_0_)^2^+(y_j_-y_0_)^2^, where x_i_ and y_i_ are the coordinates at frame i and j is final frame number(Suraneni et al., 2012). Directionalities were calculated by d/p(Suraneni et al., 2012).

#### Autocorrelation analysis

Data sets over 2.1 min (totaling 300 data points) containing total cell number in an FOV was used for autocorrelation analysis with unbiased normalization. Autocorrelation curve was fitted by single exponential with exp(-(2k)t), where k is turnover rate and t is time^22,23^. Dwell time was calculated by 1/k.

### Code availability

The codes of computer vision stabilization algorithm are available at a dedicated Github repository [https://github.com/YoshiTsukasaki/elife].

## Acknowledgements

The authors would like to acknowledge helpful discussion with Dr. M. R. Looney (UCSF) and valuable suggestion from Dr. J. M. Farahany and Dr. K. H. Yamada and experimental help from Dr. K. Zhu, Dr. J. Chen, Dr. X. P. Gao, G. Liu and D. Wang. Research reported in this publication was supported by NIH grants P01HL060678, P01HL077806, R01HL090152, R01HL125350, R01HL045638, R01HL149300, R01HL152515 and T32HL007829.

## Author Contributions

Y.T., J.R. and A.B.M. designed the research;

Y.T. performed the experiments and the majority of the analyses;

J.C.J., B.J. and D.M. helped with the live *Pseudomonas* culture and the imaging in lung; K.B, J.C and C.T. provided input on the experimental design

P.T. helped with the experimental design and optimization of the two-photon imaging system;

Z.H. and S.N. assisted with development of the surgical methods;

M.G. developed the mice of neutrophil lineage labeling;

S.P. G.M, and A.B.M. reviewed the data analysis;

Y.T., J.R. and A.B.M. wrote the initial draft of the paper. All authors reviewed and revised the manuscript.

---

**Supplementary table 1.** Comparison of Computer Vision Image registration algorithms with existing image registration algorithms and comparison of transforms in computer vision image registration algorithms. Remaining motion after stabilization and computational time are compared between intensity-based and feature-based image registration algorithms. Furthermore, these are also compared among different transforms in feature-based image registration. Remaining motion after stabilization indicates the standard deviation of motion displacement in stabilized lung movie analyzed by optical flow analysis; remaining motion after stabilization (n=3), computational time (n=3). Red text shows the algorithm we adopted. Bold text indicates best spec among algorithms. ^†^ Computational time was estimated based on movies with total 25,596,000 frames used for paper’s analysis. Additional color channel was assumed to be half computational time.

| Methods | Algorithm | Parameter set | | Motion correction accuracy ($\mu\text{m}$) | Computational time/20 frames (s) | Estimated computational time/25,596,000 frames<sup>†</sup> (days) |
| --- | --- | --- | --- | --- | --- | --- |
| Intensity-based image registration | Diffeomorphic demons<sup>12</sup> | Pyramid Levels 6, Iteration Number 100, Accumulated Field Smoothing 2 | | **0.520 ± 0.038** | 61.602 ± 0.700 (21.5) | 625.636 |
| Feature-based image registration | Feature point detection, Motion tracking, Translation transform | Feature point detection, Motion tracking (Feature point number 500, Pyramid level 6, Max. Iteration number 20, Window size 20-30, Displacement threshold 0.1) | - | 1.253 ± 0.147 | **2.802 ± 0.065 (0.97)** | 28.457 |
|  | Feature point detection, Motion tracking, Perspective transform |  | - | **0.618 ± 0.061** | **2.870 ± 0.043 (1)** | **29.148** |
| | Feature point detection, Motion tracking, Non-rigid transform | | Geometric transformation fit (Polynomial 4) | **0.563 ± 0.052** | 4.785 ± 0.041 (1.67) | 48.597 |

**Supplementary figure 1.**
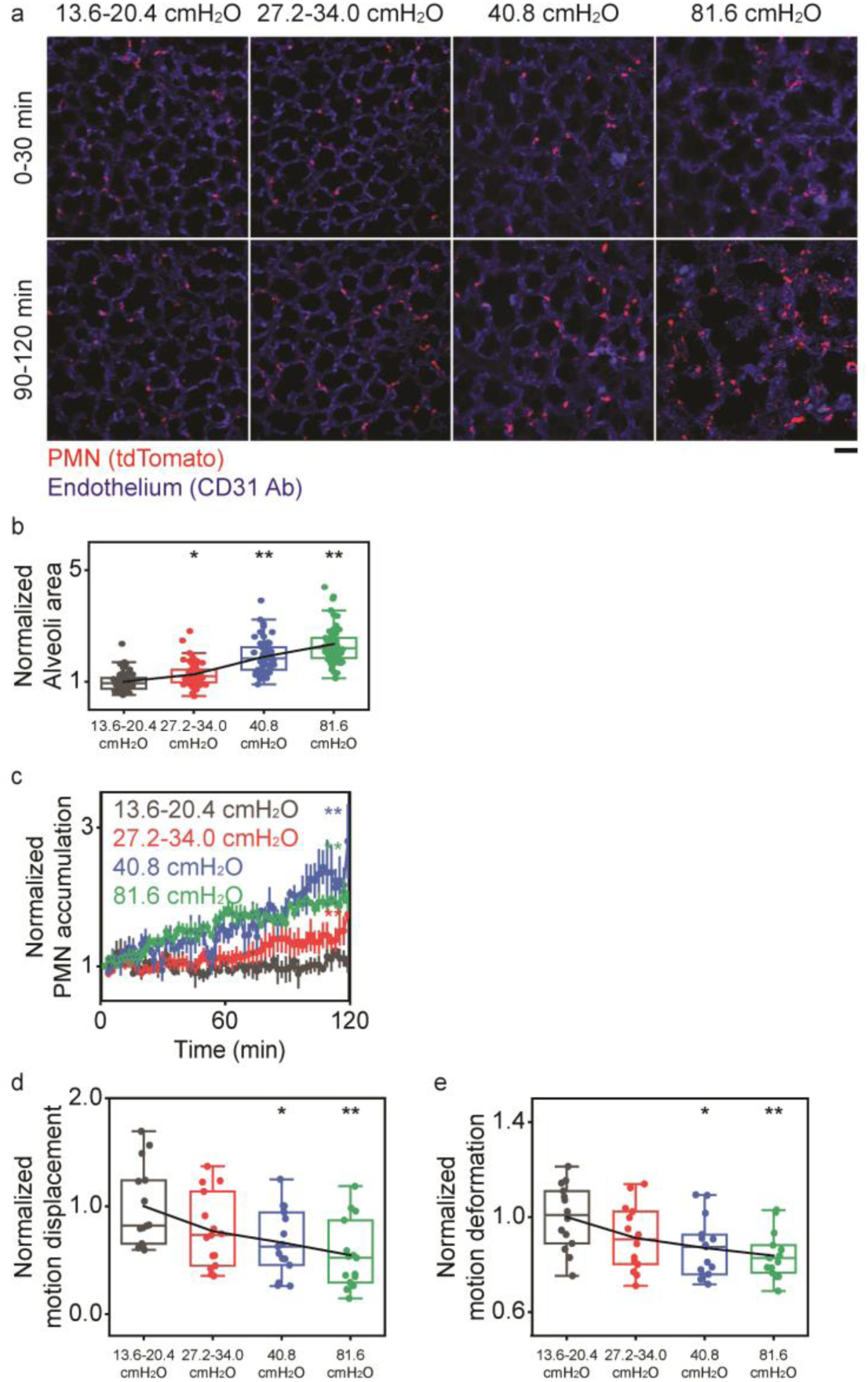
Lung Injury and Motion Artifacts During Lung Intravital Imaging using Mechanical Stabilization of Lungs. **a.** PMN accumulation as well as alveolar structural alterations in response to applied suction pressure at optical window. Scale bar; 40 μm. **b.** Increase in alveoli area in response to suction pressure; n=60 (alveoli), 5 FOVs for each suction pressure. **c.** Quantification of PMN accumulation over time; n=3 (FOV) for pressure of 13.6-20.4 cmH_2_O, n=3 (FOV) for pressure of 27.2-34.0 cmH_2_O, n=4 (FOV) of 40.8 and 81.6 cmH_2_O pressure. **d. e.** Normalized motion displacement (d) and deformation (e) in relation to suction pressure; n=14 (FOV) for 13.6-20.4 and 27.2-34.0 cmH_2_O and n=15 (FOV) for other pressures. Statistical analysis was performed using one-way ANOVA Tukey test in a, c-e; *P<0.05, **P<0.01. All two-photon images are single z slices.

**Supplementary figure 2.**
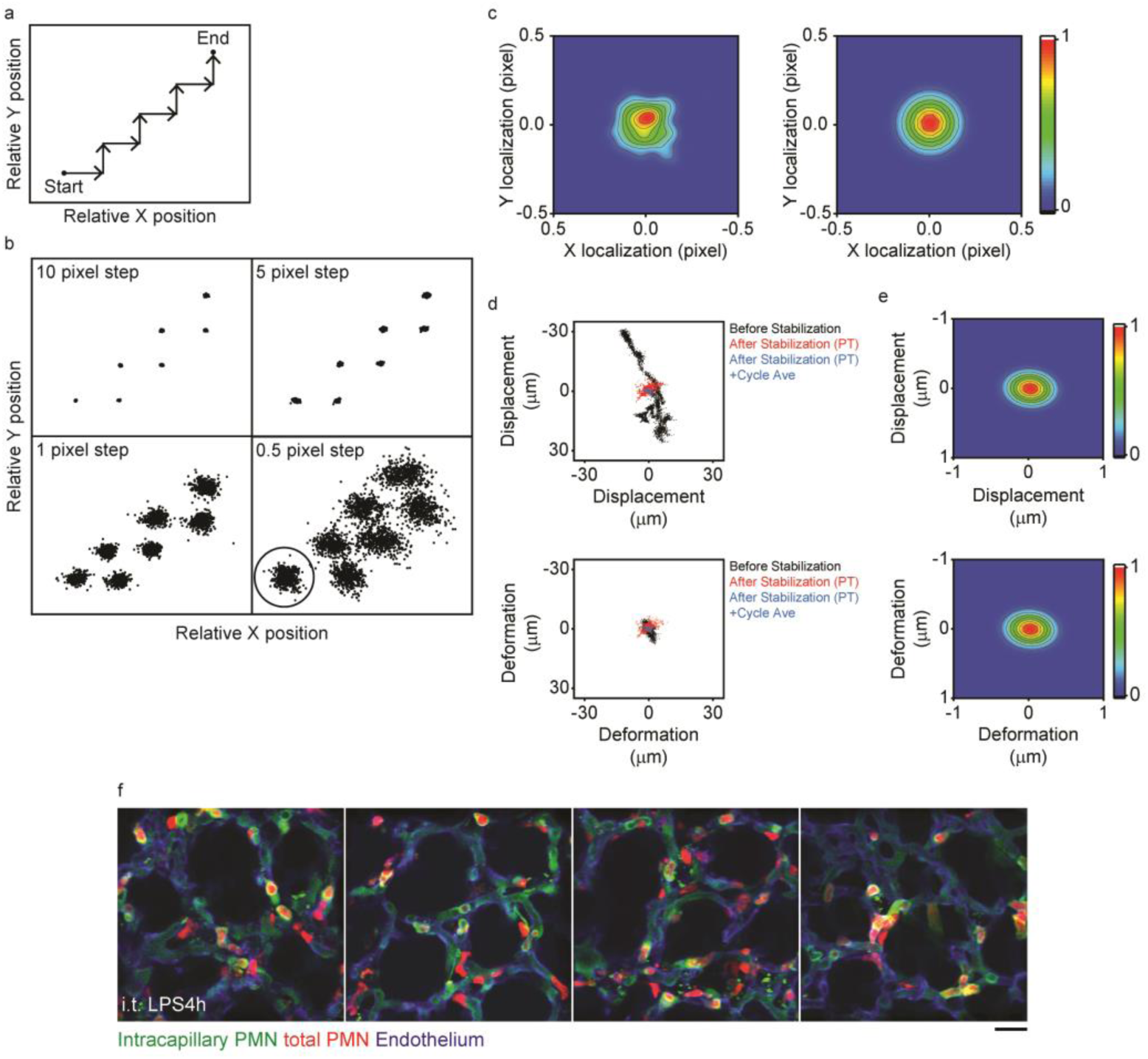
Resolution and Precision of CASTii Algorithm. **a.** Schematic drawing of experimental procedures. While images of pulmonary vessels (labeled by anti-CD31 antibody) are acquired by video-rate without ventilation, the microscope stage is moved in stepwise manner. **b.** Demonstration of effective motion tracking by optical flow analysis with a variety of step size motions. Motion tracking of feature points is performed with these points using optical flow analysis; n=3150 (Feature points) for 10-pixel step, n=3374 (Feature points) for 5 pixel step, n=3437 (Feature points) for 1 pixel step, n=3416 (Feature points) for 0.5 pixel step. **c.** Identification of precision of motion tracking. Plots from 0.5-pixel step circle (n=488, left) is analyzed by 2D Gaussian fitting, resulting in σ=0.092 pixel (right). **d.** Scatter plots of lung motion with displacement and deformation; n=4500 (Feature points) for before stabilization, for after stabilization (XY), for after stabilization (PT), for after stabilization (PT) + respiratory cycle averaging. XY indicates XY alignment algorithm, where each of averaged XY motion displacement was horizontally moved back, and PT indicates perspective transform algorithm for motion correction. **e.** The precision of the CASTii. Displacement and deformation plot after stabilization by CASTii + respiratory cycle averaging were analyzed by 2D Gaussian fitting, resulting in σ_x_=0.19 μm, σ_y_=0.14 μm for displacement and deformation, respectively. **f.** Representative compartmentalized PMN imaging using CASTii after 4h of i.t. LPS. Scale bar; 20 μm.

**Supplementary figure 3.**
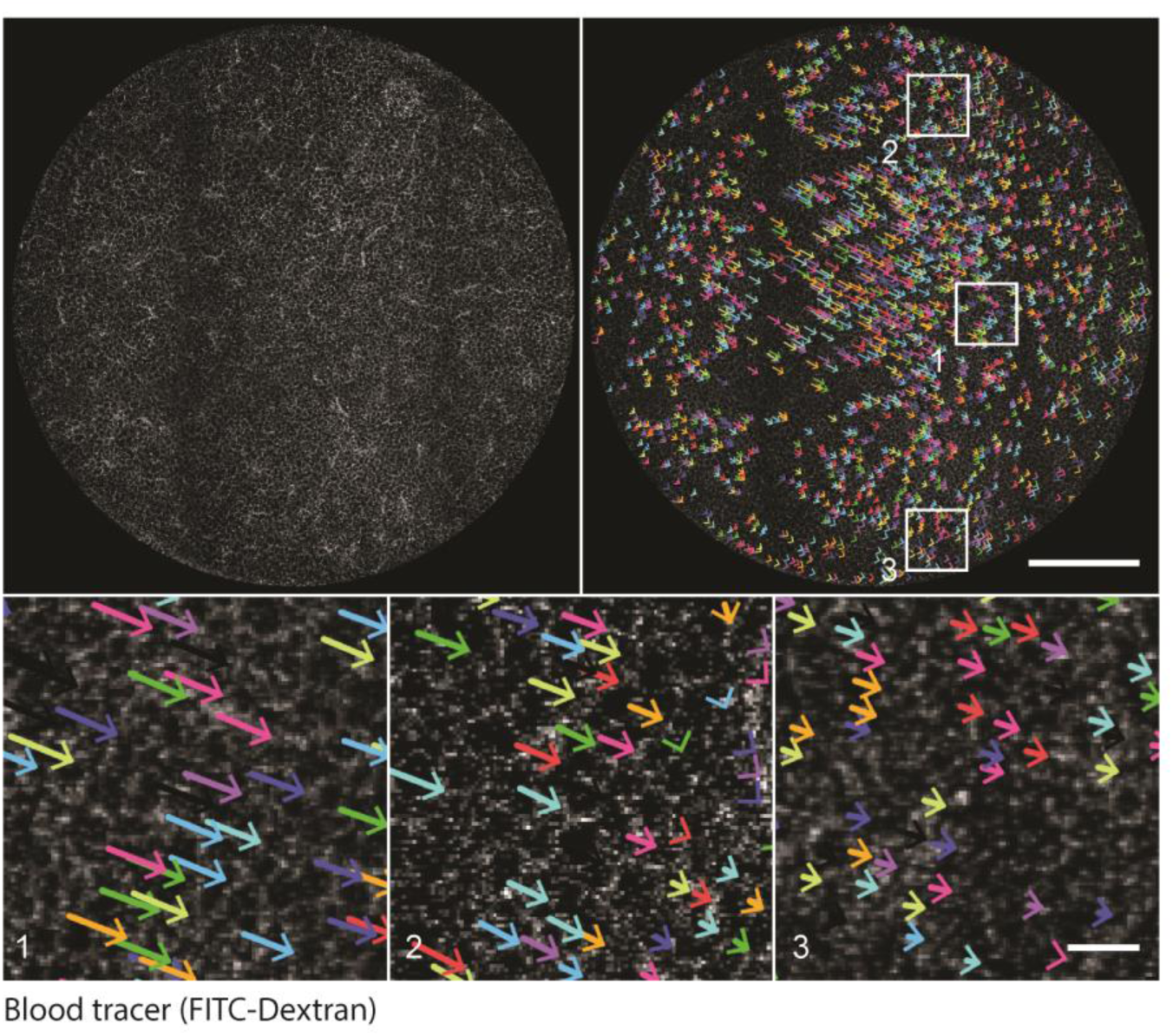
Motion Artifacts Within Lung Optical Window. Image of pulmonary microvessel structure (Top left). Image of pulmonary microvessels with motion vector with colored arrow (Top right). Numbered, inset white boxes in top right are magnified (Bottom). Scale bar; 1 mm (Top) and 100 μm (Bottom).

**Supplementary figure 4.**
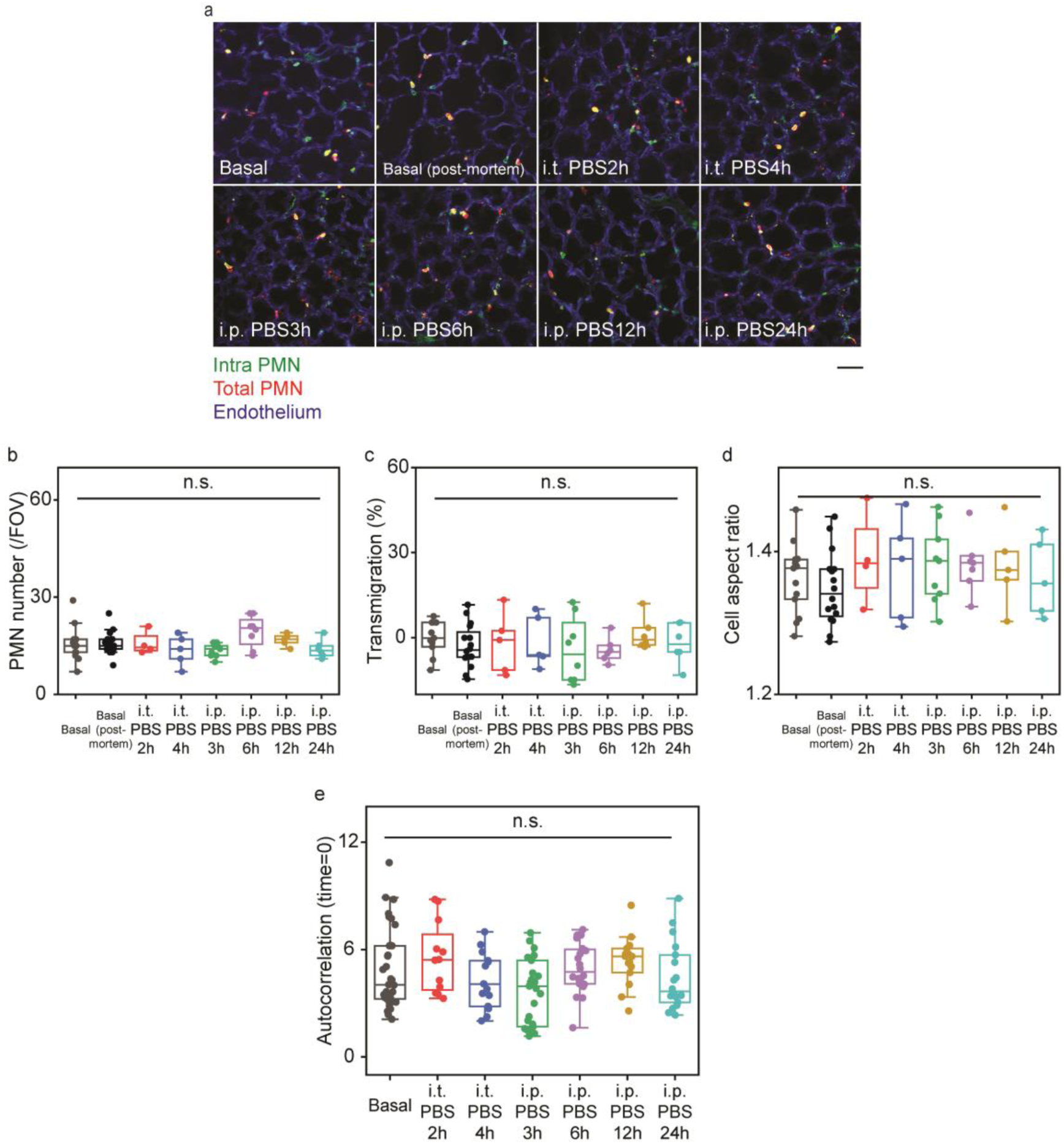
PMN Localization, Cell Number, Morphology and Autocorrelation Analysis During Control Condition in Respiring Lungs. **a.** Two-photon fluorescent images of PMNs and pulmonary microvessels under a variety of control experimental conditions. Basal (post-mortem) indicates lung in basal condition after euthanizing mice without surgery to insert optical windows. 10-seconds movies were averaged for each image. Scale bar; 40 μm. **b.** Quantitative analysis of PMN number per FOV; n=13 (FOV) for basal, n=16 (FOV) for basal (post-mortem), n=4 for i.t. PBS2h, n=5 (FOV) for i.t. PBS4h, n=9 (FOV) for i.p. PBS3h, n=18 (FOV) for i.p. PBS6h, n=5 (FOV) for i.p. PBS12h, n=6 (FOV) for i.p. PBS24h. **c.** Quantitative analysis of PMN transmigration; n=203 (cell) in 11 FOVs for basal, n=252 (cell) in 16 FOVs for basal (post-mortem), n=63 (cell) in 5 FOVs for i.t. PBS2h, n=68 (cell) in 5 FOVs for i.t. PBS4h, n=123 (cell) in 8 FOVs for i.p. PBS3h, n=185 (cell) in 6 FOVs for i.p. PBS6h, n=84 (cell) in 6 FOVs for i.p. PBS12h, n=74(cell) in 6 FOVs for i.p. PBS24h. **d.** Quantitative analysis of cell aspect ratio of PMNs; n= 6092 (cell) in 13 FOVs for basal, n= 252 (cell) in 16 FOVs for basal (post-mortem), n= 2337 (cell) in 4 FOVs for i.t. PBS2h, n= 2336 (cell) in 5 FOVs for i.t. PBS4h, n= 3859 (cell) in 9 FOVs for i.p. PBS3h, n= 4640 (cell) in 8 FOVs for i.p. PBS6h, n= 2762 (cell) in 5 FOVs for i.p. PBS12h, n= 1958 (cell) in 6 FOVs for i.p. PBS24h. **e.** Quantitative analysis of autocorrelation value at time lag of 0 in a variety of control experimental conditions; n=33 (data set) in 11 FOVs for basal, n=12 (data set) in 4 FOVs for i.t. PBS2h, n=15 (data set) in 5 FOVs for i.t. PBS4h, n=27 (data set) in 9 FOVs for i.p. PBS3h, n=24 (data set) in 8 FOVs for i.p. PBS6h, n=15 (data set) in 5 FOVs for i.p. PBS12h, n=18 (data set) in 6 FOVs for i.p. PBS24h. Statistical analysis was performed using one-way ANOVA Tukey test in b-e. All two-photon images are single z slices.

**Supplementary figure 5.**
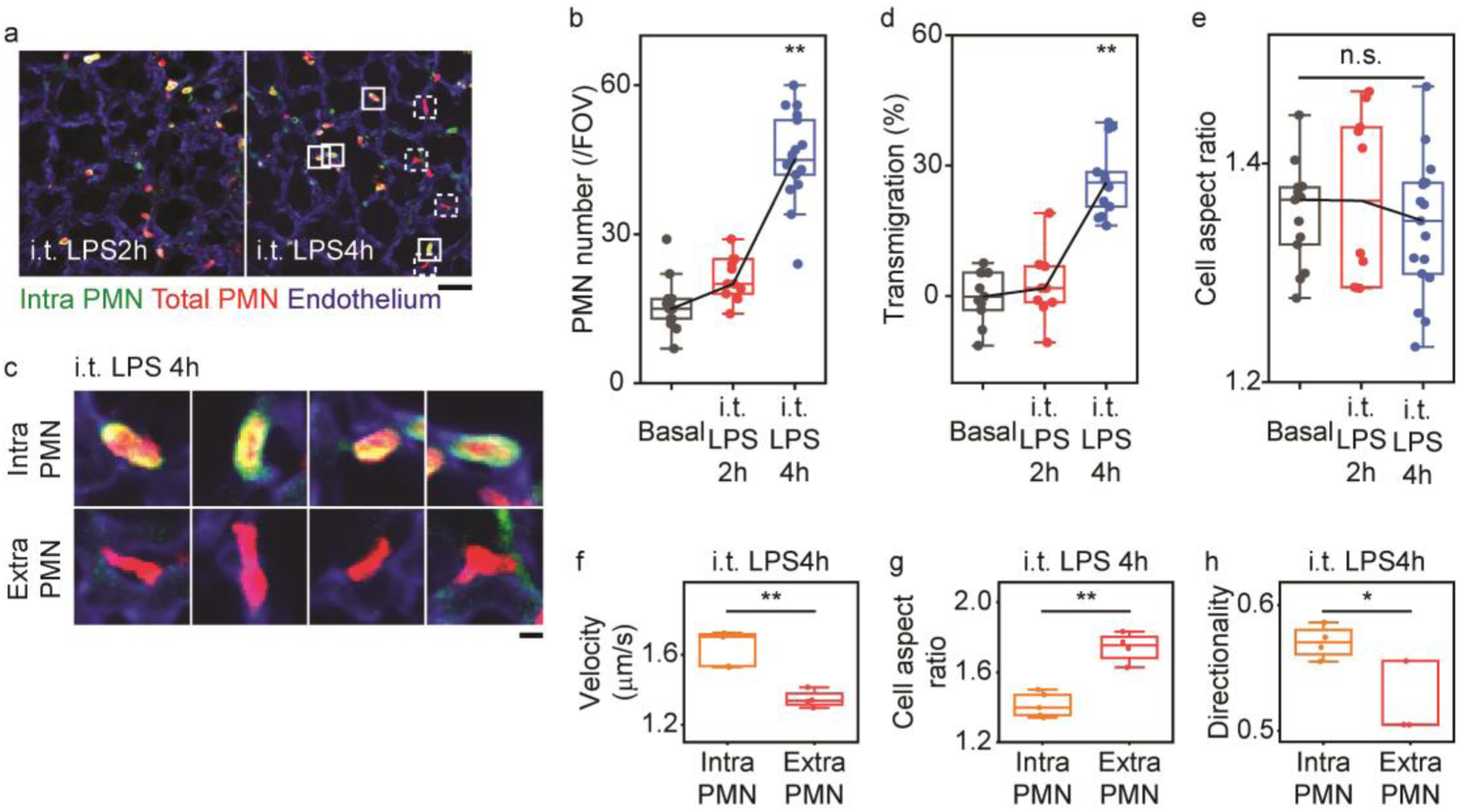
PMN Localization, Morphology and Dynamics in Response to intra-tracheal (i.t.) LPS Challenge. **a.** Two-photon fluorescent images of PMNs and pulmonary microvascular structure at different times after i.t. LPS condition. Scale bar; 40 μm. **b.** Quantitative analysis of PMN number per FOV; n=13 (FOV) for Basal, n=11 (FOV) for i.t. LPS2h, n=17 (FOV) for i.t. LPS4h. **c.** Magnified images of intravascular PMN and extravascular PMN at i.t. LPS4h. Inset solid and dotted white boxes are magnified. Scale bar; 5 μm. **d.** Quantitative analysis of PMN transmigration; n=203 (cell) in 11 FOVs for basal, n=234 (cell) in 10 FOVs for i.t. LPS2h, n=773 (cell) in 14 FOVs for i.t. LPS4h. **e.** Quantitative analysis of cell aspect ratio of PMN; n= 6092 (cell) in 13 FOVs for basal, n= 5391 (cell) in 10 FOVs for i.t. LPS2h, n= 15554 (cell) in 16 FOVs for i.t. LPS4h. **f, g, h**. Change of velocity (f) and cell aspect ratio (g) and directionality (h) of PMN migration between intravascular and extravascular PMN after 4h of i.t. LPS; n=471 (cell) in 5 (data set) and 16 FOVs for intravascular PMN (Velocity), n=302 (cell) in 4 (data set) and 16 FOVs for extravascular PMN (Velocity), n=317 (cell) in 5 (data set) and 16 FOVs for intravascular PMNs (Cell aspect ratio), n=269 (cell) in 4 (data set) and 16 FOVs for extravascular PMN (Cell aspect ratio), n=365 (cell) in 4 (data set) and 16 FOVs for intravascular PMN (Directionality), n=242 (cell) in 3 (data set) and 16 FOVs for extravascular PMN (Directionality). Statistical analysis was performed using one-way ANOVA Tukey test in b, d, and e, and two-sample t-test for f-h; *P<0.05, **P<0.01. All two-photon images are single z slices.

**Supplementary figure 6.**
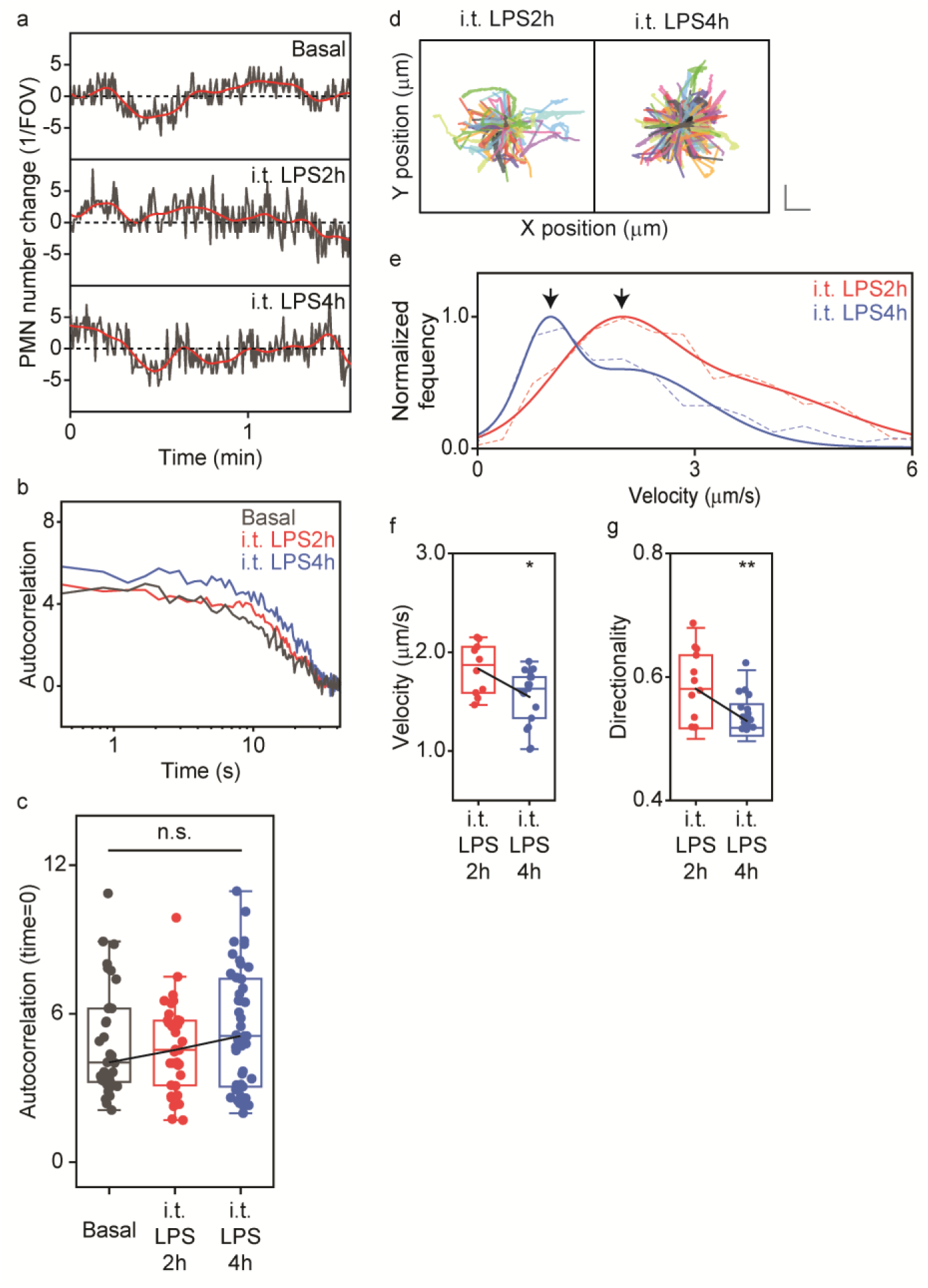
PMN Dynamics in Lungs Following intra-tracheal (i.t.) endotoxin. **a.** Total number of PMN within a FOV changes over time after i.t. LPS. **b.** Autocorrelation analysis of PMN number fluctuation at basal state and after i.t. LPS. PMN number fluctuations over 2.1 min are used for autocorrelation analysis. **c.** Quantification of autocorrelation value at time lag of 0; n=33 (data set) in 11 FOVs for basal, n=33 (data set) in 11 FOVs for i.t. LPS2h, n= 51 (data set) in 17 FOVs for i.t. LPS4h. **d.** Trajectory of PMN migration after i.t. LPS. Scale bar; 10 μm (vertical) and 10 μm (horizontal); n=351 (cell) for i.t. LPS2h, n=344 (cell) for i.t. LPS4h. **e.** Changes of velocity distribution over time after i.t. LPS. Arrowheads show the first peak obtained by double Gaussian fitting; n=503 (cell) for i.t. LPS2h, n=1068 (cell) for i.t. LPS4h. **f, g.** Quantitative analysis of velocity (f) and directionality (g) of PMN motility at different times after i.t. LPS; n= 5391 (cell) in 10 FOVs for i.t. LPS2h, n= 15554 (cell) in 16 FOVs for i.t. LPS4h. Statistical analysis was performed using one-way ANOVA Tukey test in c and two-sample t-test in f and g; *P<0.05, **P<0.01.

**Supplementary figure 7.**
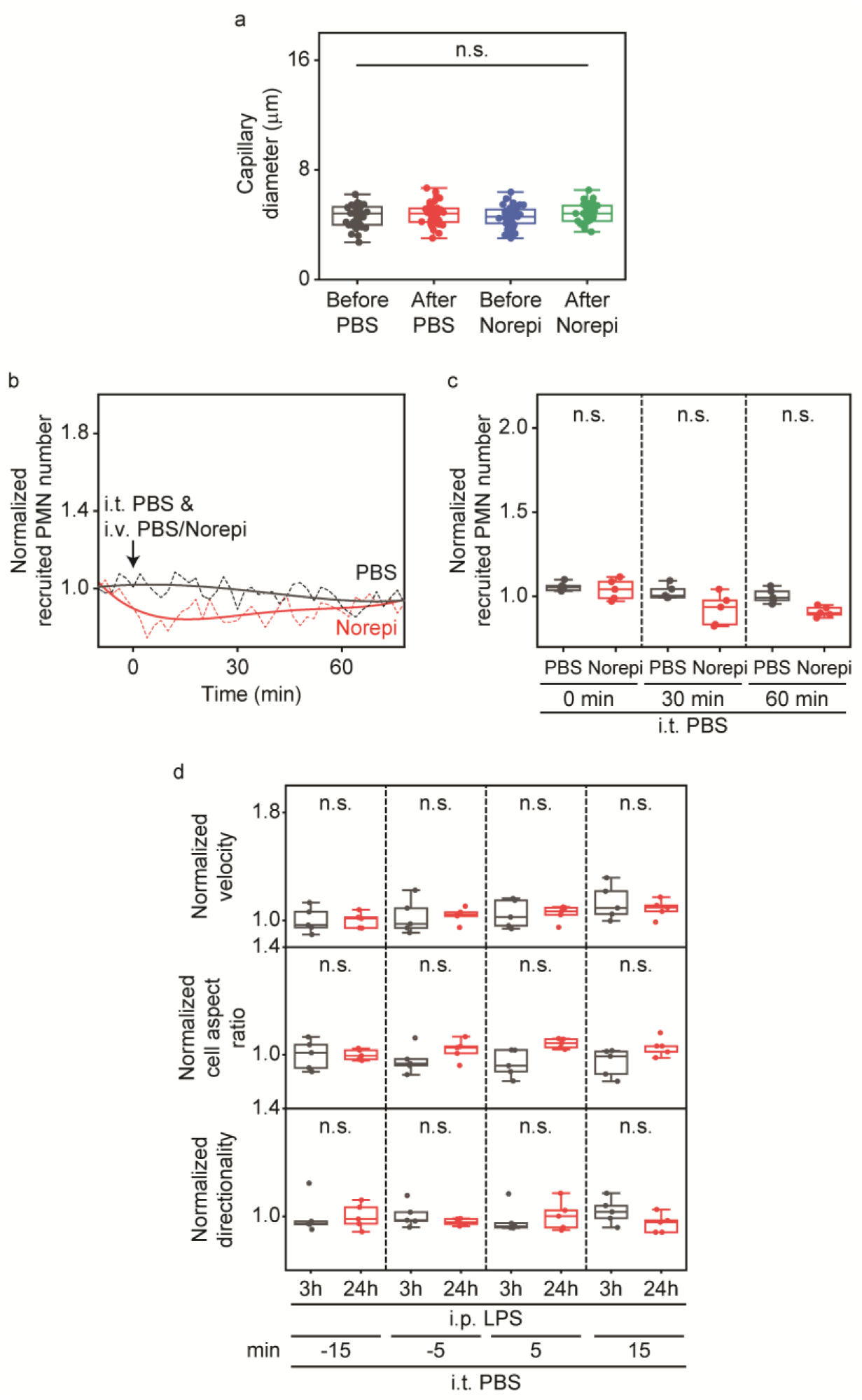
Capillary Diameters in Response to PBS or Norepinephrine and PMN Cell Number, Velocity, Cell Aspect Ratio and Directionality Under Different Experimental Conditions in Normal Respiring Lungs. **a.** Quantitative analysis of diameters of microvessels in response to i.v. injection of PBS or norepinephrine (Norepi). **b.** Recruited PMN number at times before/after i.t. inhalation of PBS and i.v. injection of PBS/norepinephrine (Norepi). Solids lines indicate polynomial fitting results. **c.** Quantitative analysis of recruited PMN number; n=5 (FOVs) in i.t. PBS i.v. PBS/Norepi. **d.** Quantitative analysis of velocity, cell aspect ratio, and directionality before and after i.t. PBS at i.p. LPS3h/24h condition; n=900-1200 (cell) for i.p. LPS3h, n=1300-1800 (cell) for i.p. LPS24h. Statistical analysis was performed using one-way ANOVA Tukey test in a, c, d.

**Supplementary figure 8.**
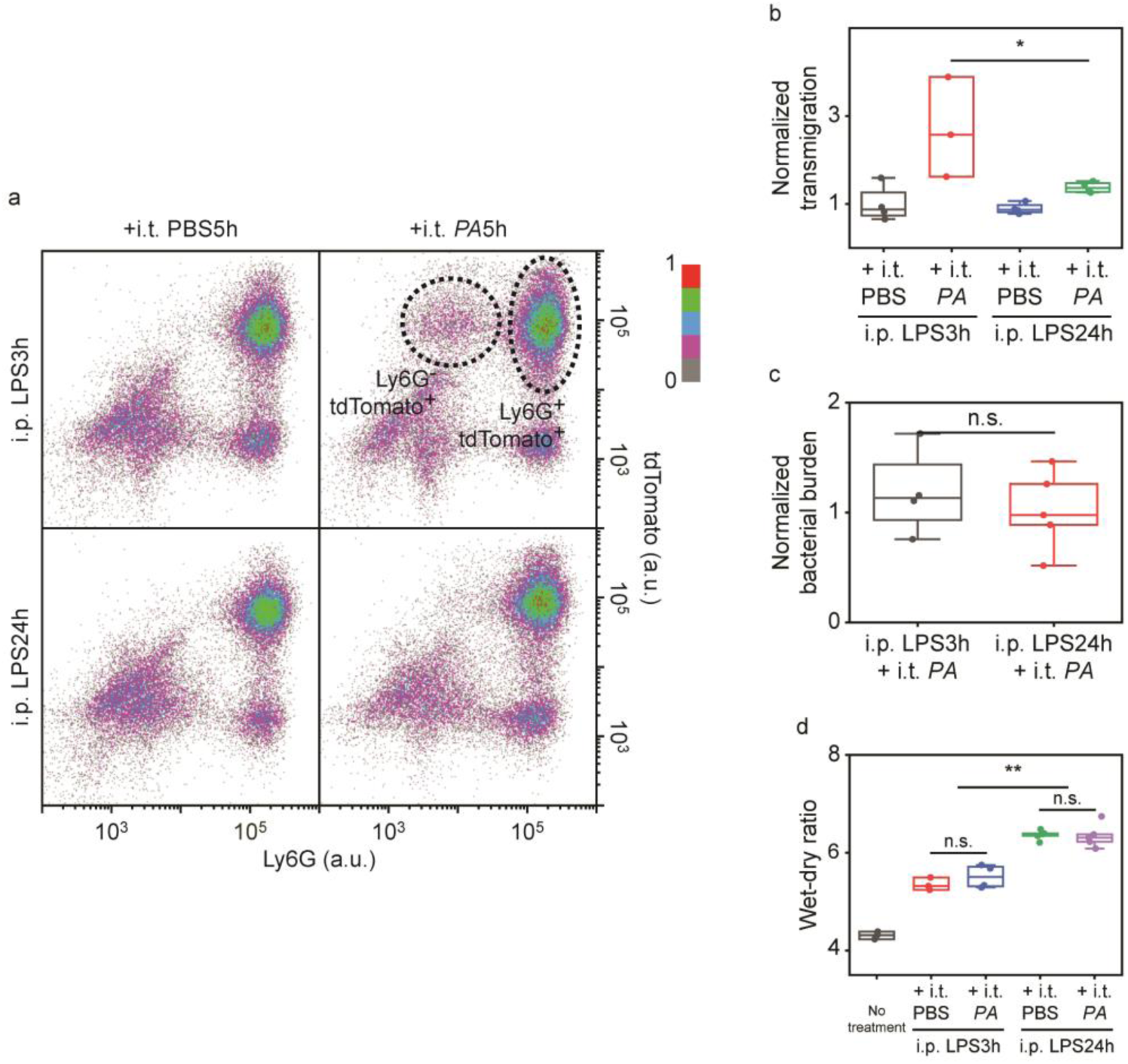
Whole Lung Analysis of PMN Transmigration and Bacterial Burden and Wet/Dry Ratio. **a.** FACS analysis of PMN transmigration occurrence in response to i.t. *PA* injection at different time points (3h or 24h) after i.p. LPS; n=27.453-32,080 (Cell) for all cases. **b.** Quantitative analysis of PMN transmigration in response to i.t. *PA* injection at different time points (3h or 24h) after i.p. LPS; n=4 (mice) for i.p. LPS3h + i.t. PBS, i.p. LPS24h + i.t. PBS and i.p. LPS24h + i.t. *PA*, n=3 (mice) for i.p. LPS3h + i.t. *PA*. Transmigration occurrences were calculated by transmigrated PMN number (tdTomato^+^Ly6G^-^ cell number) / total PMN number (tdTomato^+^ cell number). **c.** Quantitative analysis of bacterial burden of mice lungs in response to i.t. *PA* injection at different time points (3h/24h) after i.p. LPS; n=4 (mice) for i.p. LPS3h + i.t. *PA*, n=5 (mice) for i.p. LPS24h + i.t. *PA*. **d.** Quantitative analysis of wet-dry ratio of mice lungs in response to i.t. *PA* injection at different time points (3h or 24h) after i.p. LPS; n=3 (mice) for no treatment and i.p. LPS3h + i.t. PBS, n=4 (mice) for i.p. LPS3h + i.t. *PA*, n=5 (mice) for i.p. LPS24h + i.t. PBS and i.p. LPS24h + i.t. *PA*. Statistical analysis was performed using one-way ANOVA Tukey test in b-d; *P<0.05, **P<0.01.

**Supplementary movie1. Motion Artifacts in the Lung Optical Window.** Movie of pulmonary microvascular structure (Left). Movie of pulmonary microvessels with feature points (Right). Lung motion is acquired at video-rate and the motion of feature points are tracked by optical flow analysis. Scale bar; 1 mm. Time stamp; msec.

**Supplementary movie2. Stabilization Algorithm in CASTii.** Pulmonary microvessels are labeled by intravenously injected CD31 antibody. Shi and Tomasi method is used to detect feature points in reference image (red feature points in Motion tracking movie). These feature points are tracked using Lucas-Kanade optical flow method, where red feature points are derived from reference image and green feature points are tracked (Motion tracking image movie). Estimated motion is corrected using homography matrix calculated from matched feature points (Motion correction movie). Finally, the motion from respiratory cycle is averaged for each cycle (Respiratory cycle averaging movie).

Scale bar; 20 μm. Time stamp; msec. All two-photon movies are single z slices.

**Supplementary movie3. Compartmentalized Lung PMN Imaging after Intra-tracheal LPS Using CASTii.** Scale bar; 40 μm. Time stamp; min:sec. All two-photon movies are single z slices.

**Supplementary movie4. Representative lung PMN Images with Conventional Method and CASTii.** Left images are original images with conventional methods. Right images are stabilized images with CASTii.

Scale bar; 50 μm. All two-photon movies are single z slices.

**Supplementary movie5. Comparison of Existing Image Registration Algorithms with CASTii and Comparison Among Motion Correction Algorithms in CASTii.** Our computer vision image registration algorithm (feature-based methods) was compared with the existing image registration algorithms, diffeomorphic demons as an example of the existing intensity-based methods. Motion correction algorithm in CASTii is compared with translational transform, where each of averaged XY motion displacements were horizontally moved back, perspective transform algorithm and non-rigid transform. Left image is original image before motion correction.

Scale bar; 20 μm. Time stamp; msec. All two-photon movies are single z slices.

**Supplementary movie6. Passage of PMNs in narrow pulmonary microvessels.** Scale bar; 40 μm. Time stamp; min:sec. All two-photon movies are single z slices.

**Supplementary movie7. PMN circulation following intratracheal LPS challenge.** PMN turnover is observed in basal state and i.t. LPS. Scale bar; 40 μm. Time stamp; min:sec. All two-photon movies are single z slices.

**Supplementary movie8. PMN Dynamics during Endotoxemia Induced with Intraperitoneal LPS.** PMN turnover completely disappears following i.p. LPS. Scale bar; 40 μm. Time stamp; min:sec. All two-photon movies are single z slices.

**Supplementary movie9. PMN Trafficking in Lungs during Endotoxemia induced by Intraperitoneal LPS.** In the condition of i.p. LPS, almost all PMNs attach microvessel endothelial cells and PMNs slowly move along microvessels. Migration velocity is progressively reduced after i.p. LPS. Scale bar; 40 μm. Time stamp; min:sec. All two-photon movies are single z slices.

**Supplementary movie10. Effects of Norepinephrine on Dynamics of Lung PMN Marinated Pool.** After i.v. norepinephrine, PMN number is reduced and turnover rate of PMN increases. Scale bar; 40 μm. Time stamp; min:sec. All two-photon movies are single z slices.

**Supplementary movie11. Effects of Norepinephrine on PMN Recruitment Dynamics Induced by fMLP Inhalation.** PMN dynamics after i.t. fMLP and i.v. PBS/norepinephrine, is visualized. Compared with i.t. fMLP or i.v. PBS, PMN recruitment is reduced and delayed by i.v. norepinephrine. Scale bar; 60 μm. Time stamp; hr:min:sec. All two-photon movies are single z slices.

**Supplementary movie12. Effects of fMLP in 3h vs. 24h endotoxemia models.** After i.t. fMLP, PMN changes their migratory behavior at i.p. LPS3h condition but not i.p. LPS24h. Scale bar; 40 μm. Time stamp; min:sec. All two-photon movies are single z slices.

