## Supplemental movies for "Computer-Vision Stabilized Intravital Imaging Reveals Lung Capillary Neutrophil Dynamics Crucial for Lung Host-Defense Function"

Asrar B. Malik, Ph.D.
Schweppe Family Distinguished Professor
Head of Department of Pharmacology and
Regenerative Medicine
University of Illinois College of Medicine
835 South Wolcott Ave, E403 MSB

Chicago, IL 60612


### Figures and legends

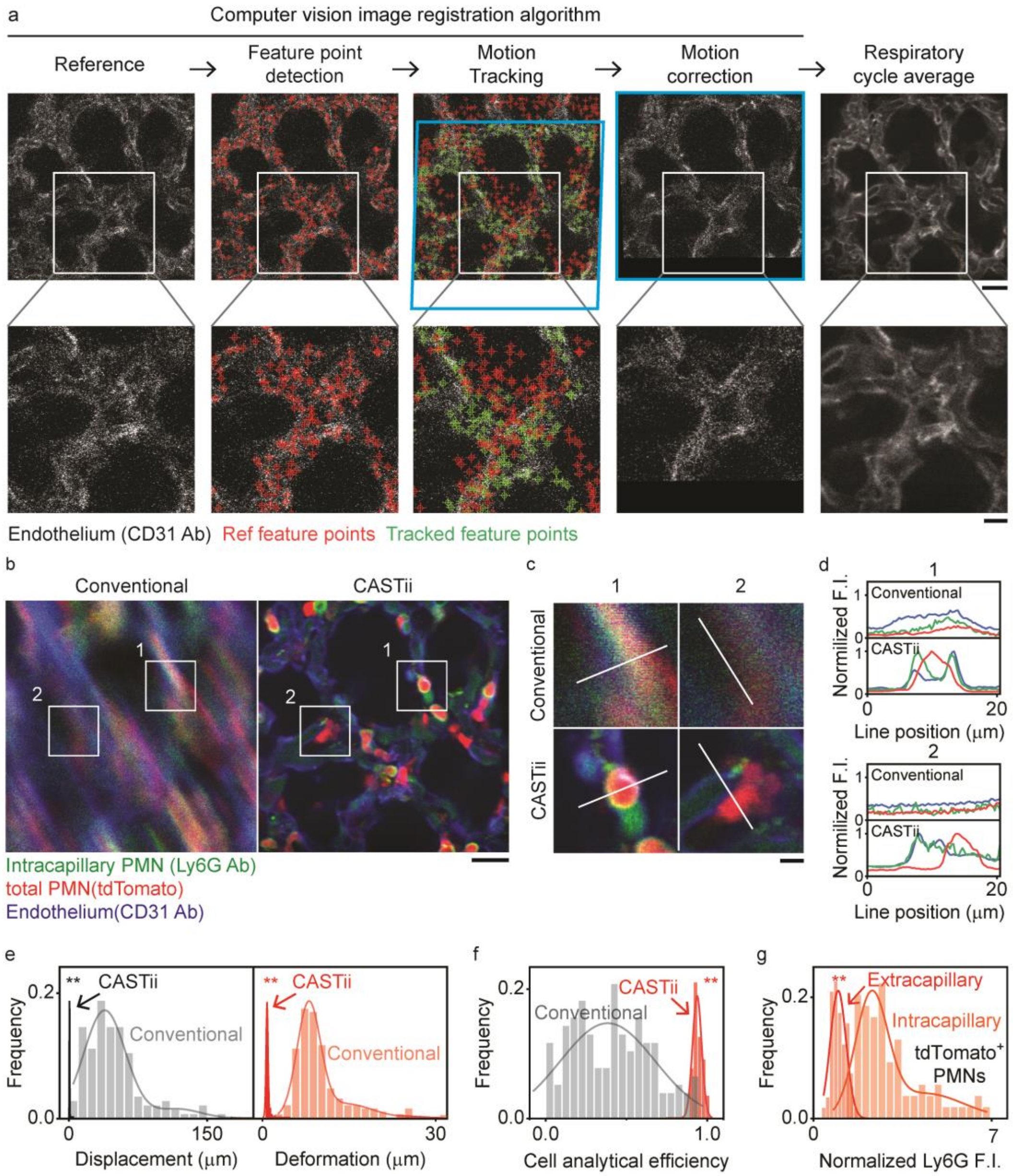


**Figure 1. Lung intravital imaging in normally respiring lungs using CASTii.**

**a.** Flow chart of the stabilization processing of CASTii using an example set of lung images. Pulmonary microvessels are imaged; feature points are detected algorithmically in designated reference image (red crosses indicate the feature points). Feature points are tracked to calculate their deviation from reference (green crosses indicate tracked feature points). This estimated motion is corrected and followed by averaging over the respiratory cycle. Inset white boxes in the top row are magnified at the bottom row. Scale bar; 20 μm (Top) and 10 μm (Bottom).

**b.** PMN imaging using CASTii after 4h of i.t. LPS demonstrates clear gain in precise stabilization. Left image: Conventional, right image: using CASTii. Scale bar; 20 μm.

**c.** Magnified view from inset white boxes in b. Scale bar; 5 μm. d, Line profile analysis of PMNs and fine lung structures along white lines drawn in c.

**e.** Quantification of motion displacement and deformation via peak-to-peak movements using conventional methods and CASTii; n=143 (FOV).

**f.** Quantification of cell analytical efficiency with conventional methods and CASTii; n=143 (FOV).

**g.** Compartmental PMN localization with Ly6G fluorescent intensity; n=58 (Cell, Extracapillary PMN) and n=77 (Cell, Intracapillary PMN).

Statistical analysis was performed using two-sample t-test in e-g. All two-photon images are single z slices.

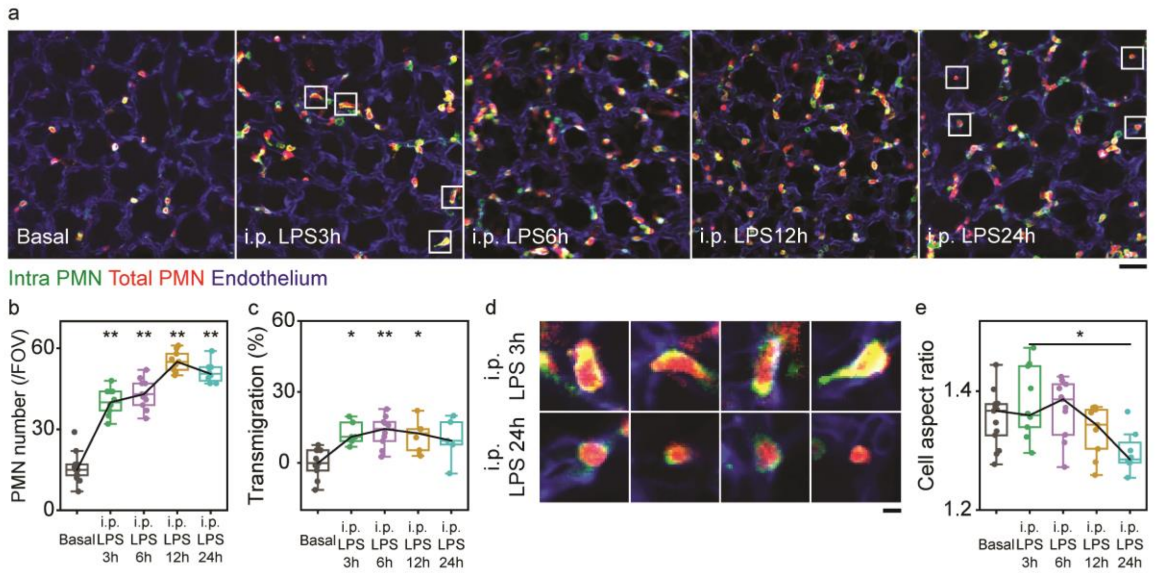


**Figure 2. PMN morphological changes and lung localization in endotoxemia.**

**a.** Two-photon fluorescence images of PMNs and lung microvessels at different time points after i.p. LPS challenge. 10-seconds movies were averaged for each image. Scale bar; 40 μm.

**b.** Quantitative analysis of PMN number per FOV; n=13 (FOV) for basal, n=10 (FOV) for i.p. 3h LPS, n=11 (FOV) for i.p. 6h LPS, n=9 (FOV) for i.p. 12h LPS, n=8 (FOV) for i.p. 24h LPS.

**c.** Quantitative analysis of PMN transmigration; n=203 (cell) in 11 FOVs for basal, n=234 (cell), n=473 (cell) in 6 FOVs for i.p. 3h LPS, n=504 (cell) in 10 FOVs for i.p. 6h LPS, n=497 (cell) in 6 FOVs for i.p. 12h LPS, n=379 (cell) in 5 FOVs for i.p. 24h LPS.

**d.** Magnified PMN morphology after i.p. 3h LPS and i.p. 24h LPS. Inset white boxes in a are magnified. Scale bar; 5 μm.

**e.** Quantitative analysis of cell aspect ratio of PMN; n= 6092 (cells) in 13 FOVs for basal, n= 10841 (cell) in 8 FOVs for i.p. 3h LPS, n= 13784 (cell) in 10 FOVs for i.p. 6h LPS, n= 12277 (cell) in 8 FOVs for i.p. 12h LPS, n= 9243 (cell) in 8 FOVs for i.p. 24h LPS.

Statistical analysis was performed using one-way ANOVA Tukey test for b, c and e; *P<0.05, **P<0.01. All two-photon images are single z slices.

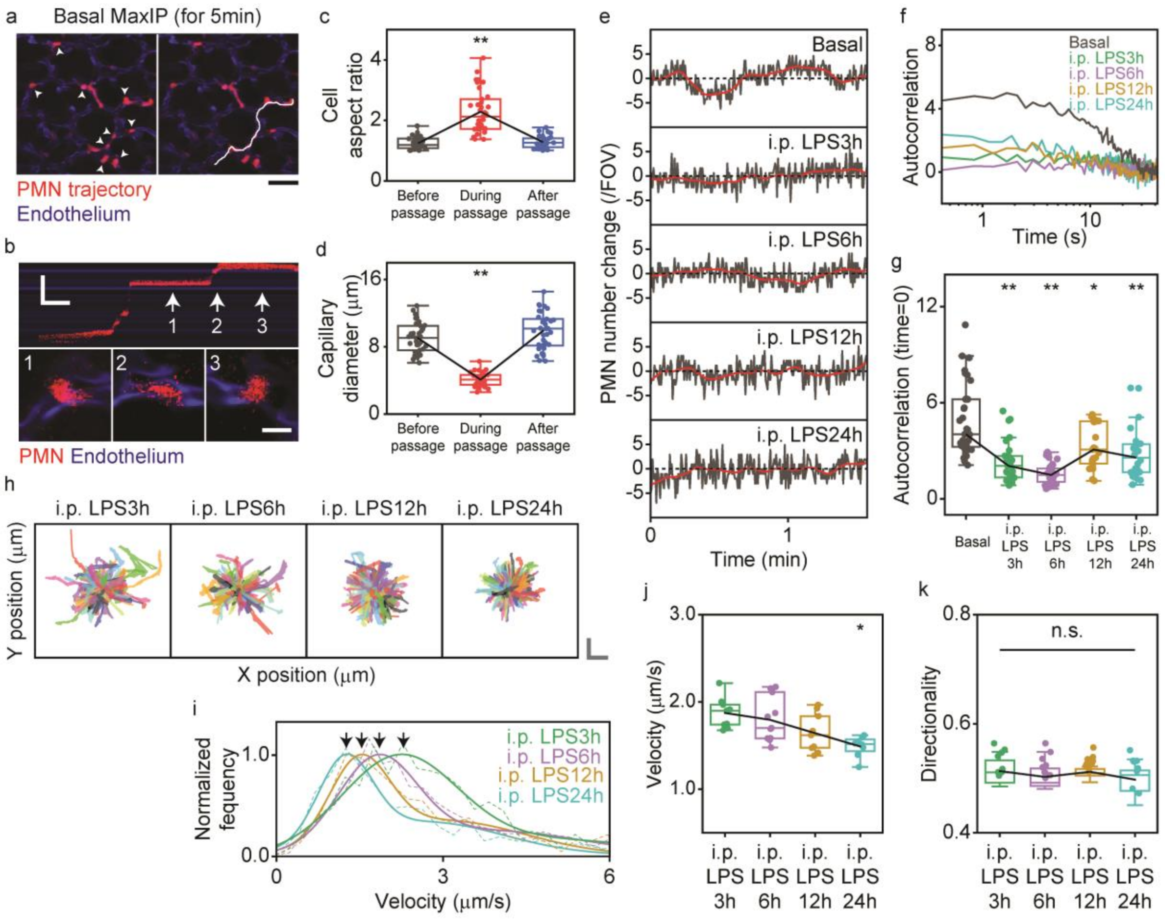


**Figure 3. PMN dynamics at rest and in response to endotoxemia in respiring lungs.**

**a.** PMN trajectory at baseline. Maximum intensity projection (MaxIP) over 5 min is shown as well as PMN trajectory in white (right panel); lung microvessels in which transiently trapped PMNs are labeled (arrow heads). Scale bar; 40 μm.

**b.** Kymograph showing PMN passage in lung microvessels. The white trajectory line was examined via kymograph to demonstrate the “catch and release” kinetics of PMNs. Scale bar; 50 μm (vertical) and 10 s (horizontal). Morphology of PMNs at 3 time points (1,2,3) within the vessel, as indicated by white arrow, is shown at the bottom. Scale bar; 10 μm.

**c.** Quantitative analysis of cell aspect ratio during PMN passage; n=35 (cell) in 12 FOVs.

**d.** Quantitative analysis of diameter of lung microvessels during PMN passage.

**e.** Alterations in total number of PMNs within a FO over time recorded under a variety of conditions.

**f.** Autocorrelation analysis of PMN number fluctuations at baseline and after i.p. LPS challenge. PMN number fluctuations over 2.1 min are used for autocorrelation analysis.

**g.** Quantification of autocorrelation values at time lag of 0; n=33 (data set) in 11 FOVs for Basal, n=30 (data set) in 11 FOVs for i.p. 3h LPS, n=27 (data set) in 10 FOVs for i.p. 6h LPS, n=27 (data set) in 9 FOVs for i.p. 12h LPS, n=24 (data set) in 8 FOV for i.p. 24h LPS.

**h.** Trajectory of PMN migration during endotoxemia induced by i.p. LPS. Scale bar; 10 μm (vertical) and 10 μm (horizontal); n=304 (cell) for i.p. 3h LPS, n=329 (cell) for i.p. 6h LPS, n=384 (cell) for i.p. 12h LPS, and n=307 (cell) for i.p. 24h LPS.

**i.** Changes in velocity distribution over time after i.p. LPS. Arrowheads indicate the first peak obtained by double Gaussian fitting; n=898 (cell) for i.p. 3h LPS, n=692 (cell) for i.p. 6h LPS, n=805 (cell) for i.p. 12h LPS, and n=1171 (cell) for i.p. 24h LPS.

**j.** Quantitative analysis of PMN velocity at different times after i.p. LPS.

**k.** Quantitative analysis of directionality of PMN motility at different times after i.p. LPS; n=10841 (cell) in 8 FOVs for i.p. 3h LPS, n= 13784 (cell) in 10 FOVs for i.p. 6h LPS, n= 12277 (cell) in 8 FOVs for i.p. 12h LPS, n= 9243 (cell) in 8 FOVs for i.p. 24h LPS. Statistical analysis was performed using one-way repeated measures ANOVA Tukey test in c and d, and one-way ANOVA Tukey test in g, j, k; *P<0.05, **P<0.01. All two-photon images are single z slices.

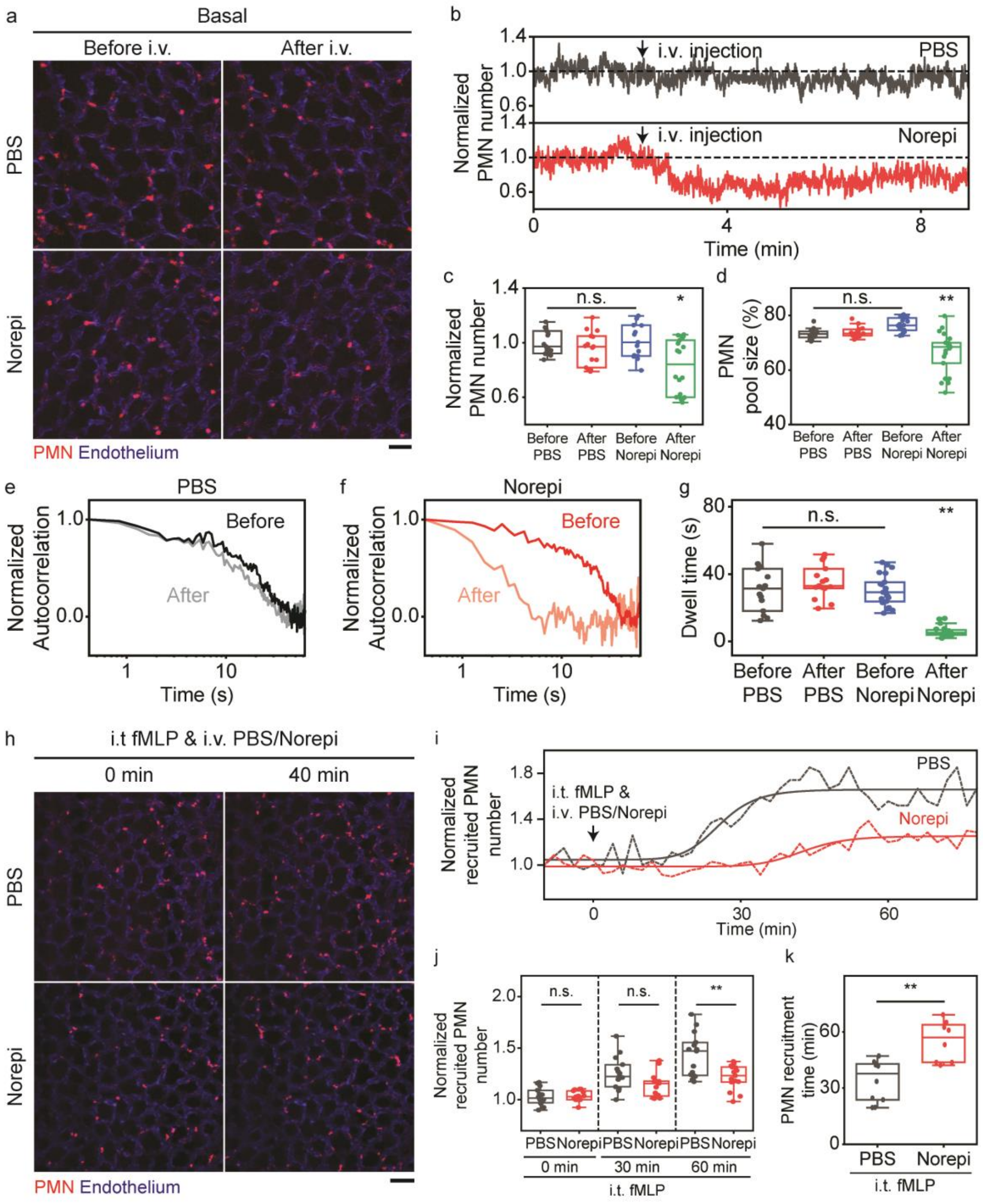


**Figure 4. Dynamics of marginated PMN pool in lung vessels.**

**a.** Two-photon fluorescence images of PMNs and pulmonary microvessels before and after i.v. injection of PBS or norepinephrine (Norepi). Scale bar; 40 μm.

**b.** PMN number changes in response to i.v. injection of PBS or norepinephrine.

**c. d.** Quantitative analysis of PMN number (c) and PMN pool size (d) before andafter i.v. injection of PBS/norepinephrine (Norepi). PMN pool size is defined as ratio of stationary and total PMN (see text); n=15 (data set) in 5 FOVs for before PBS, n=15 (data set) in 5 FOVs for after PBS, n=15 (data set) in 7 FOVs for before Norepi, n=18 (data set) in 7 FOVs for after Norepi.

**e. f.** Autocorrelation analysis of PMN number fluctuations before abd after i.v. injection of PBS (e) or norepinephrine (Norepi) (f).

**g.** Quantitative analysis of dwell time from autocorrelation curves; n=15 (data set) in 5 FOVs for before PBS, n=15 (data set) in 5 FOVs for after PBS, n=21 (data set) in 7 FOVs for before Norepi, and n=21 (data set) in 7 FOVs for after Norepi.

**h.** Two-photon fluorescence images of PMNs and pulmonary microvessels after i.t. inhalation of fMLP and i.v. injection of PBS/norepinephrine (Norepi). Scale bar; 60 μm.

**i.** Recruitment of PMNs in response to i.t. inhalation of fMLP and i.v. injection of PBS or norepinephrine (Norepi). Recruited PMNs are defined as PMNs remaining in FOV for more than 4 min.

**j.** Quantitative analysis of PMN recruitment after i.t. inhalation of fMLP and i.v. injection of PBS or norepinephrine (Norepi). n=15 (FOVs) in i.t. fMLP i.v. PBS or Norepi.

**k.** Quantitative analysis of PMN recruitment time. Recruitment time was determined by using logistics fitting only when the raising phase appeared; n= 10 (FOVs) in i.t. fMLP i.v. PBS and n=8 (FOVs) in i.t. fMLP i.v. Norepi.

Statistical analysis was performed using one-way ANOVA Tukey test in c and d and g and j and two-sample t-test in k; *P<0.05, **P<0.01. All two-photon images are single z slices.

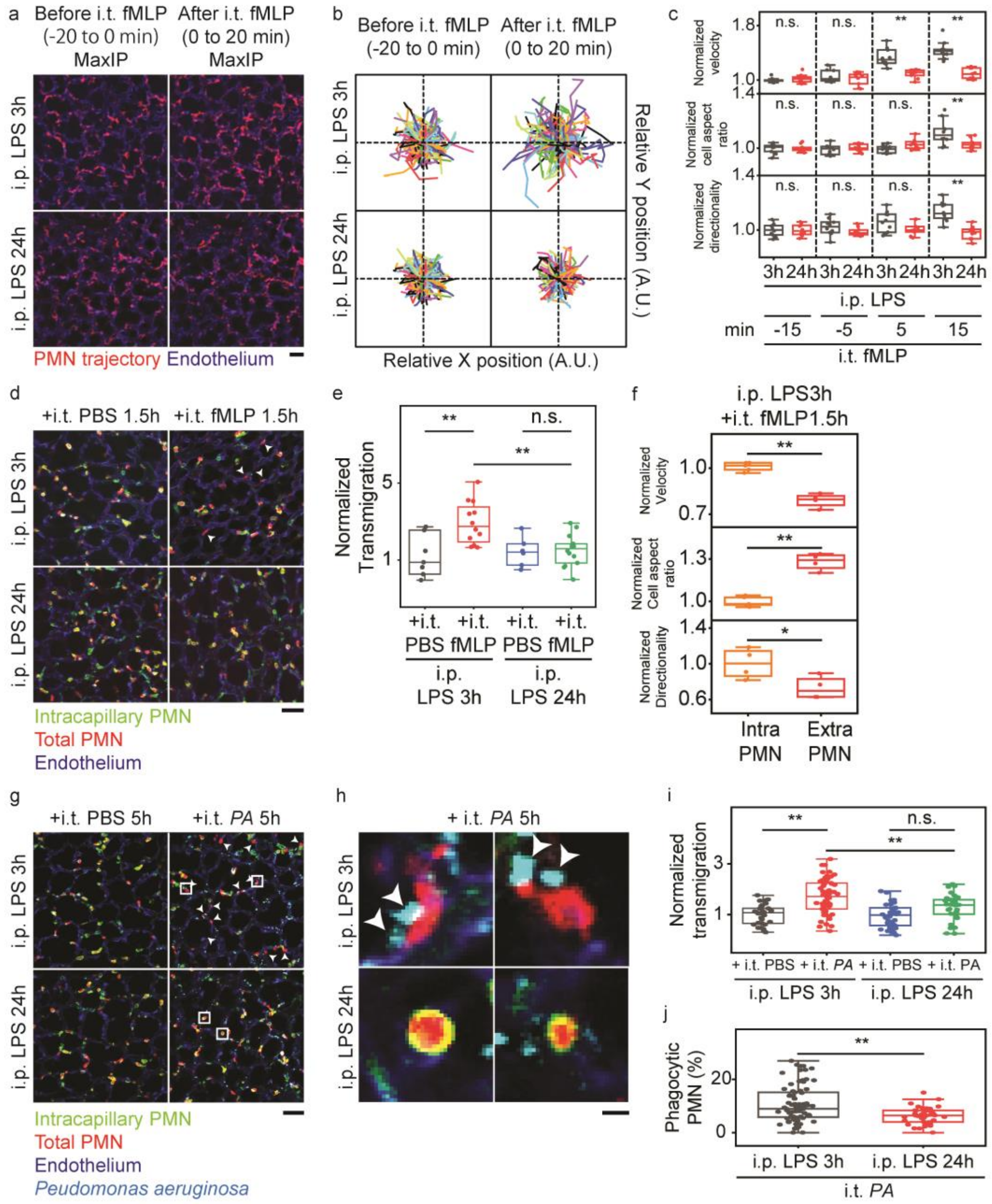

**Figure 5. Time dependent defects in PMN transmigration and phagocytosis in lungs post-endotoxemia**

**a.** Two-photon fluorescent images of PMN trajectory and pulmonary microvessels before and after i.t. fMLP in mice challenged i.p. LPS for 3h or 24h. Results show maximum intensity projection over 20 min as PMN trajectory. Scale bar; 40 μm.

**b.** Trajectory of PMN migration before and after i.t. fMLP in i.p. 3h vs. 24h LPS in mice; n=178 (cell) for before i.t. fMLP at i.p. 3h LPS condition, n=216 (cell) for after i.t. fMLP at i.p.

3h LPS condition, n=261 (cell) for before i.t. fMLP at i.p. 24h LPS condition, and 240 (cell) for after i.t. fMLP at i.p. 24h LPS condition.

All two-photon images are single z slices.

| Methods | Algorithm | Parameter set | | Motion correction accuracy (μm) | Computational time/20 frames (s) | Estimated computational time/25,596,000 frames† (days) |
|---|---|---|---|---|---|---|
| Intensity-based image registration | Diffeomorphic demons[12] | Pyramid Levels 6, Iteration Number 100, Accumulated Field Smoothing 2 | | **0.520 ± 0.038** | 61.602 ± 0.700 (21.5) | 625.636 |
| Feature-based image registration | Feature point detection, Motion tracking, Translation transform | Feature point detection, Motion tracking (Feature point number 500, Pyramid level 6, Max. Iteration number 20, Window size 20-30, Displacement threshold 0.1) | - | 1.253 ± 0.147 | **2.802 ± 0.065 (0.97)** | 28.457 |
| | Feature point detection, Motion tracking, Perspective transform | | - | **0.618 ± 0.061** | **2.870 ± 0.043 (1)** | **29.148** |
| | Feature point detection, Motion tracking, Non-rigid transform | | Geometric transformation fit (Polynominal 4) | **0.563 ± 0.052** | 4.785 ± 0.041 (1.67) | 48.597 |

**Supplementary table 1. Comparison of Computer Vision Image registration algorithms with existing image registration algorithms and comparison of transforms in computer vision image registration algorithms.** Remaining motion after stabilization and computational time are compared between intensity-based and feature-based image registration algorithms. Furthermore, these are also compared among different transforms in feature-based image registration. Remaining motion after stabilization indicates the standard deviation of motion displacement in stabilized lung movie analyzed by optical flow analysis; remaining motion after stabilization (n=3), computational time (n=3). Red text shows the algorithm we adopted. Bold text indicates best spec among algorithms. † Computational time was estimated based on movies with total 25,596,000 frames used for paper's analysis. Additional color channel was assumed to be half computational time.

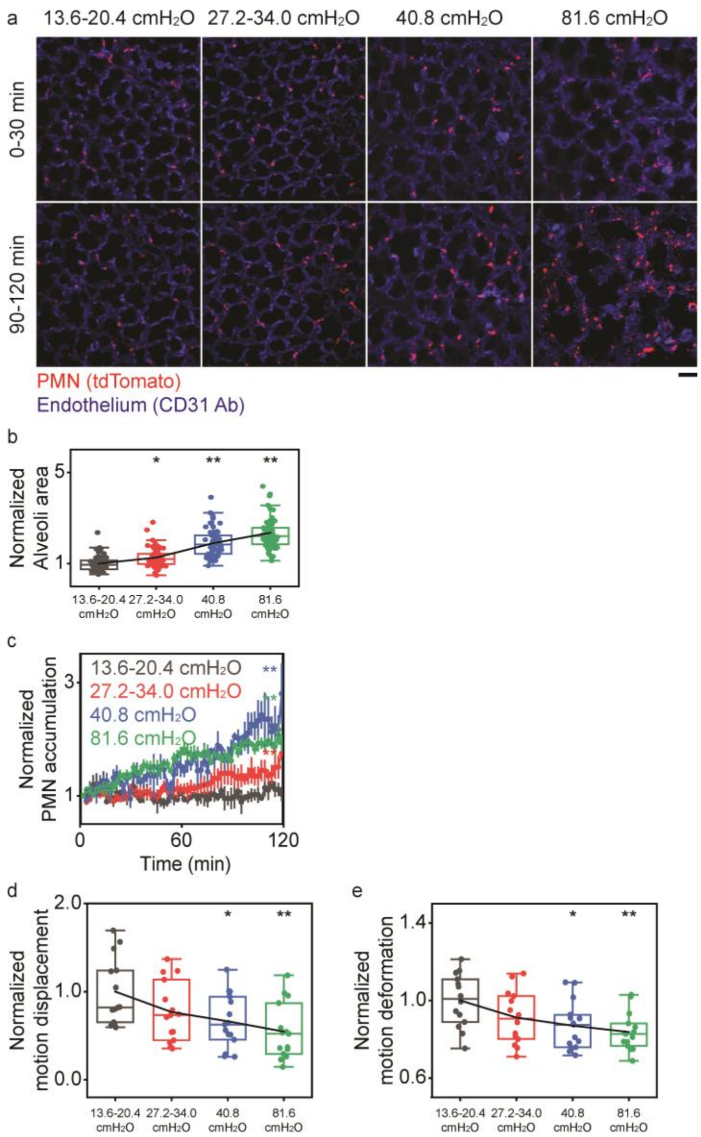


**Supplementary figure 1. Lung Injury and Motion Artifacts During Lung Intravital Imaging using Mechanical Stabilization of Lungs.**

**a.** PMN accumulation as well as alveolar structural alterations in response to applied suction pressure at optical window. Scale bar; 40 $\mu$m.

**b.** Increase in alveoli area in response to suction pressure; n=60 (alveoli), 5 FOVs for each suction pressure.

**c.** Quantification of PMN accumulation over time; n=3 (FOV) for pressure of 13.6-20.4 $cmH_2O$, n=3 (FOV) for pressure of 27.2-34.0 $cmH_2O$, n=4 (FOV) of 40.8 and 81.6 $cmH_2O$ pressure.

**d. e.** Normalized motion displacement (d) and deformation (e) in relation to suction pressure; n=14 (FOV) for 13.6-20.4 and 27.2-34.0 $cmH_2O$ and n=15 (FOV) for other pressures. Statistical analysis was performed using one-way ANOVA Tukey test in a, c-e; *P<0.05, **P<0.01. All two-photon images are single z slices.

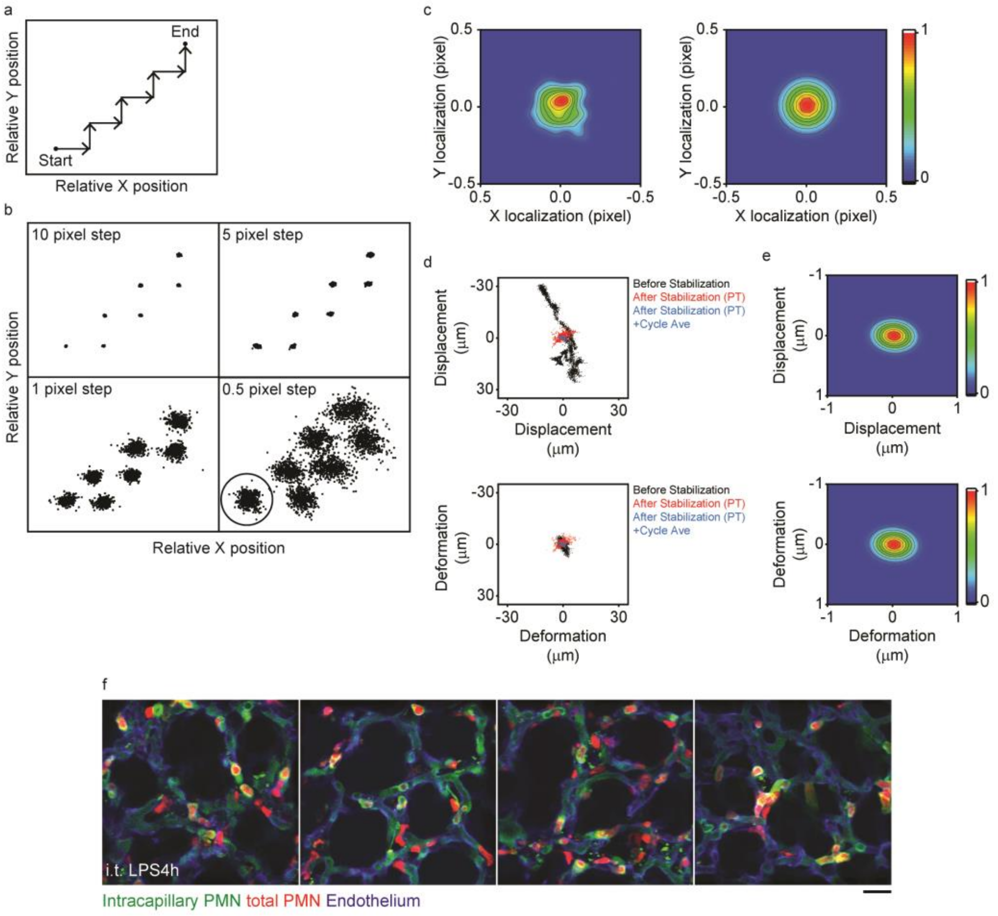


**Supplementary figure 2. Resolution and Precision of CASTii Algorithm.**

**a.** Schematic drawing of experimental procedures. While images of pulmonary vessels (labeled by anti-CD31 antibody) are acquired by video-rate without ventilation, the microscope stage is moved in stepwise manner.

**b.** Demonstration of effective motion tracking by optical flow analysis with a variety of step size motions. Motion tracking of feature points is performed with these points using optical flow analysis; n=3150 (Feature points) for 10-pixel step, n=3374 (Feature points) for 5 pixel step, n=3437 (Feature points) for 1 pixel step, n=3416 (Feature points) for 0.5 pixel step.

**c.** Identification of precision of motion tracking. Plots from 0.5-pixel step circle (n=488, left) is analyzed by 2D Gaussian fitting, resulting in $\sigma$=0.092 pixel (right).

**d.** Scatter plots of lung motion with displacement and deformation; n=4500 (Feature points) for before stabilization, for after stabilization (XY), for after stabilization (PT), for after stabilization (PT) + respiratory cycle averaging. XY indicates XY alignment algorithm, where each of averaged XY motion displacement was horizontally moved back, and PT indicates perspective transform algorithm for motion correction.

**e.** The precision of the CASTii. Displacement and deformation plot after stabilization by CASTii + respiratory cycle averaging were analyzed by 2D Gaussian fitting, resulting in $\sigma_x$=0.19 $\mu$m, $\sigma_y$=0.14 $\mu$m for displacement and deformation, respectively.

**f.** Representative compartmentalized PMN imaging using CASTii after 4h of i.t. LPS. Scale bar; 20 $\mu$m.

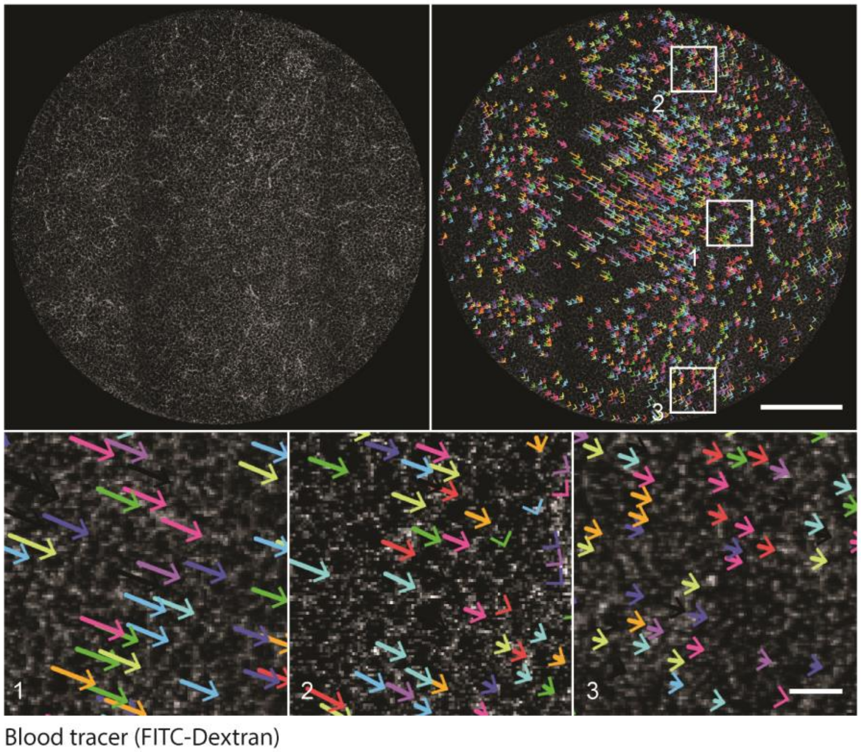


**Supplementary figure 3. Motion Artifacts Within Lung Optical Window.** Image of pulmonary microvessel structure (Top left). Image of pulmonary microvessels with motion vector with colored arrow (Top right). Numbered, inset white boxes in top right are magnified (Bottom). Scale bar; 1 mm (Top) and 100 μm (Bottom).

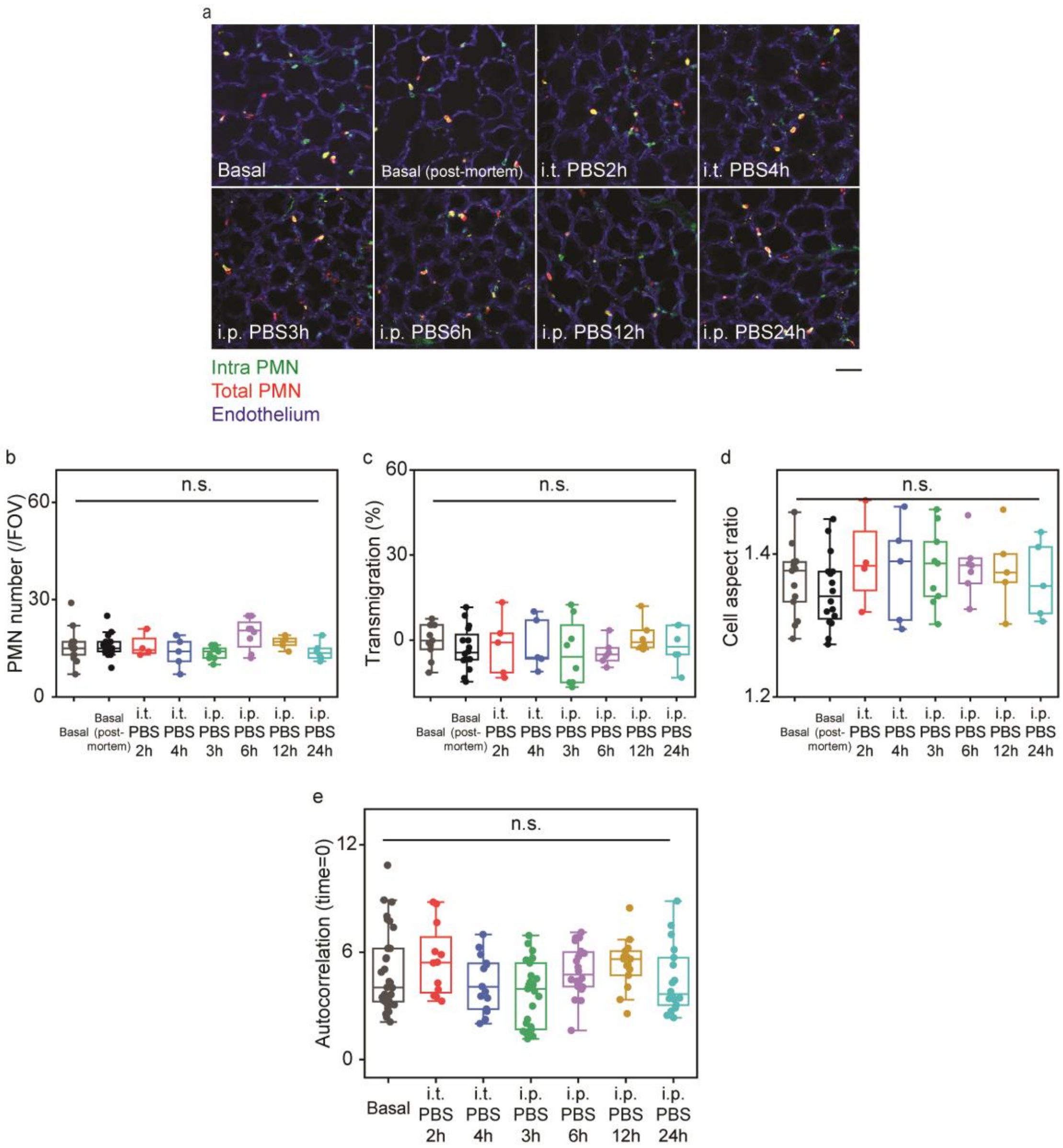


**Supplementary figure 4. PMN Localization, Cell Number, Morphology and Autocorrelation Analysis During Control Condition in Respiring Lungs.**

**a.** Two-photon fluorescent images of PMNs and pulmonary microvessels under a variety of control experimental conditions. Basal (post-mortem) indicates lung in basal condition after euthanizing mice without surgery to insert optical windows. 10-seconds movies were averaged for each image. Scale bar; 40 μm.

**b.** Quantitative analysis of PMN number per FOV; n=13 (FOV) for basal, n=16 (FOV) for basal (post-mortem), n=4 for i.t. PBS2h, n=5 (FOV) for i.t. PBS4h, n=9 (FOV) for i.p. PBS3h, n=18 (FOV) for i.p. PBS6h, n=5 (FOV) for i.p. PBS12h, n=6 (FOV) for i.p. PBS24h.

**c.** Quantitative analysis of PMN transmigration; n=203 (cell) in 11 FOVs for basal, n=252 (cell) in 16 FOVs for basal (post-mortem), n=63 (cell) in 5 FOVs for i.t. PBS2h, n=68 (cell) in 5 FOVs for i.t. PBS4h, n=123 (cell) in 8 FOVs for i.p. PBS3h, n=185 (cell) in 6 FOVs for i.p. PBS6h, n=84 (cell) in 6 FOVs for i.p. PBS12h, n=74(cell) in 6 FOVs for i.p. PBS24h.

**d.** Quantitative analysis of cell aspect ratio of PMNs; n= 6092 (cell) in 13 FOVs for basal, n= 252 (cell) in 16 FOVs for basal (post-mortem), n= 2337 (cell) in 4 FOVs for i.t. PBS2h, n= 2336 (cell) in 5 FOVs for i.t. PBS4h, n= 3859 (cell) in 9 FOVs for i.p. PBS3h, n= 4640 (cell) in 8 FOVs for i.p. PBS6h, n= 2762 (cell) in 5 FOVs for i.p. PBS12h, n= 1958 (cell) in 6 FOVs for i.p. PBS24h.

**e.** Quantitative analysis of autocorrelation value at time lag of 0 in a variety of control experimental conditions; n=33 (data set) in 11 FOVs for basal, n=12 (data set) in 4 FOVs for i.t. PBS2h, n=15 (data set) in 5 FOVs for i.t. PBS4h, n=27 (data set) in 9 FOVs for i.p. PBS3h, n=24 (data set) in 8 FOVs for i.p. PBS6h, n=15 (data set) in 5 FOVs for i.p. PBS12h, n=18 (data set) in 6 FOVs for i.p. PBS24h.

Statistical analysis was performed using one-way ANOVA Tukey test in b-e. All two-photon images are single z slices.

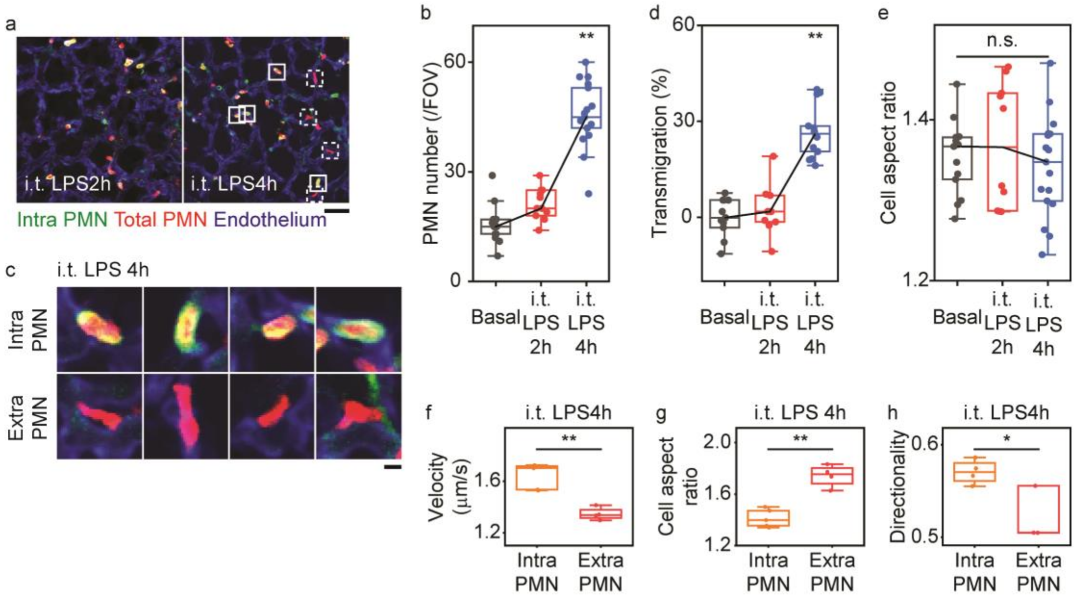


**Supplementary figure 5. PMN Localization, Morphology and Dynamics in Response to intra-tracheal (i.t.) LPS Challenge.**

**a.** Two-photon fluorescent images of PMNs and pulmonary microvascular structure at different times after i.t. LPS condition. Scale bar; 40 μm.

**b.** Quantitative analysis of PMN number per FOV; n=13 (FOV) for Basal, n=11 (FOV) for i.t. LPS2h, n=17 (FOV) for i.t. LPS4h.

**c.** Magnified images of intravascular PMN and extravascular PMN at i.t. LPS4h. Inset solid and dotted white boxes are magnified. Scale bar; 5 μm.

**d.** Quantitative analysis of PMN transmigration; n=203 (cell) in 11 FOVs for basal, n=234 (cell) in 10 FOVs for i.t. LPS2h, n=773 (cell) in 14 FOVs for i.t. LPS4h.

**e.** Quantitative analysis of cell aspect ratio of PMN; n= 6092 (cell) in 13 FOVs for basal, n= 5391 (cell) in 10 FOVs for i.t. LPS2h, n= 15554 (cell) in 16 FOVs for i.t. LPS4h.

**f, g, h**. Change of velocity (f) and cell aspect ratio (g) and directionality (h) of PMN migration between intravascular and extravascular PMN after 4h of i.t. LPS; n=471 (cell) in 5 (data set) and 16 FOVs for intravascular PMN (Velocity), n=302 (cell) in 4 (data set) and 16 FOVs for extravascular PMN (Velocity), n=317 (cell) in 5 (data set) and 16 FOVs for intravascular PMNs (Cell aspect ratio), n=269 (cell) in 4 (data set) and 16 FOVs for extravascular PMN (Cell aspect ratio), n=365 (cell) in 4 (data set) and 16 FOVs for intravascular PMN (Directionality), n=242 (cell) in 3 (data set) and 16 FOVs for extravascular PMN (Directionality).

Statistical analysis was performed using one-way ANOVA Tukey test in b, d, and e, and two-sample t-test for f-h; *P<0.05, **P<0.01. All two-photon images are single z slices.

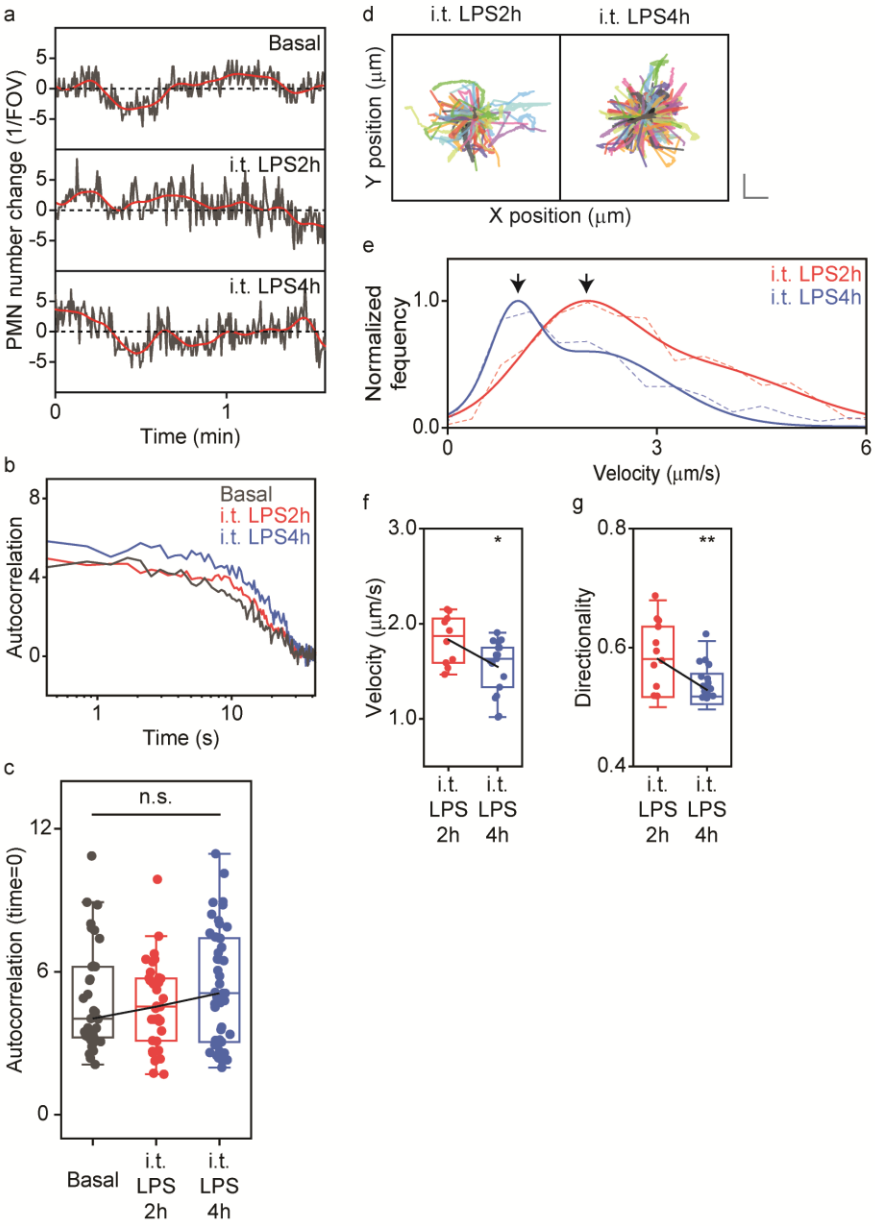


**Supplementary figure 6. PMN Dynamics in Lungs Following intra-tracheal (i.t.) endotoxin.**

**a.** Total number of PMN within a FOV changes over time after i.t. LPS.

**b.** Autocorrelation analysis of PMN number fluctuation at basal state and after i.t. LPS. PMN number fluctuations over 2.1 min are used for autocorrelation analysis.

**c.** Quantification of autocorrelation value at time lag of 0; n=33 (data set) in 11 FOVs for basal, n=33 (data set) in 11 FOVs for i.t. LPS2h, n= 51 (data set) in 17 FOVs for i.t. LPS4h.

**d.** Trajectory of PMN migration after i.t. LPS. Scale bar; 10 μm (vertical) and 10 μm (horizontal); n=351 (cell) for i.t. LPS2h, n=344 (cell) for i.t. LPS4h.

**e.** Changes of velocity distribution over time after i.t. LPS. Arrowheads show the first peak obtained by double Gaussian fitting; n=503 (cell) for i.t. LPS2h, n=1068 (cell) for i.t. LPS4h.

**f, g.** Quantitative analysis of velocity (f) and directionality (g) of PMN motility at different times after i.t. LPS; n= 5391 (cell) in 10 FOVs for i.t. LPS2h, n= 15554 (cell) in 16 FOVs for i.t. LPS4h.

Statistical analysis was performed using one-way ANOVA Tukey test in c and two-sample t-test in f and g; $*P<0.05$, $**P<0.01$.

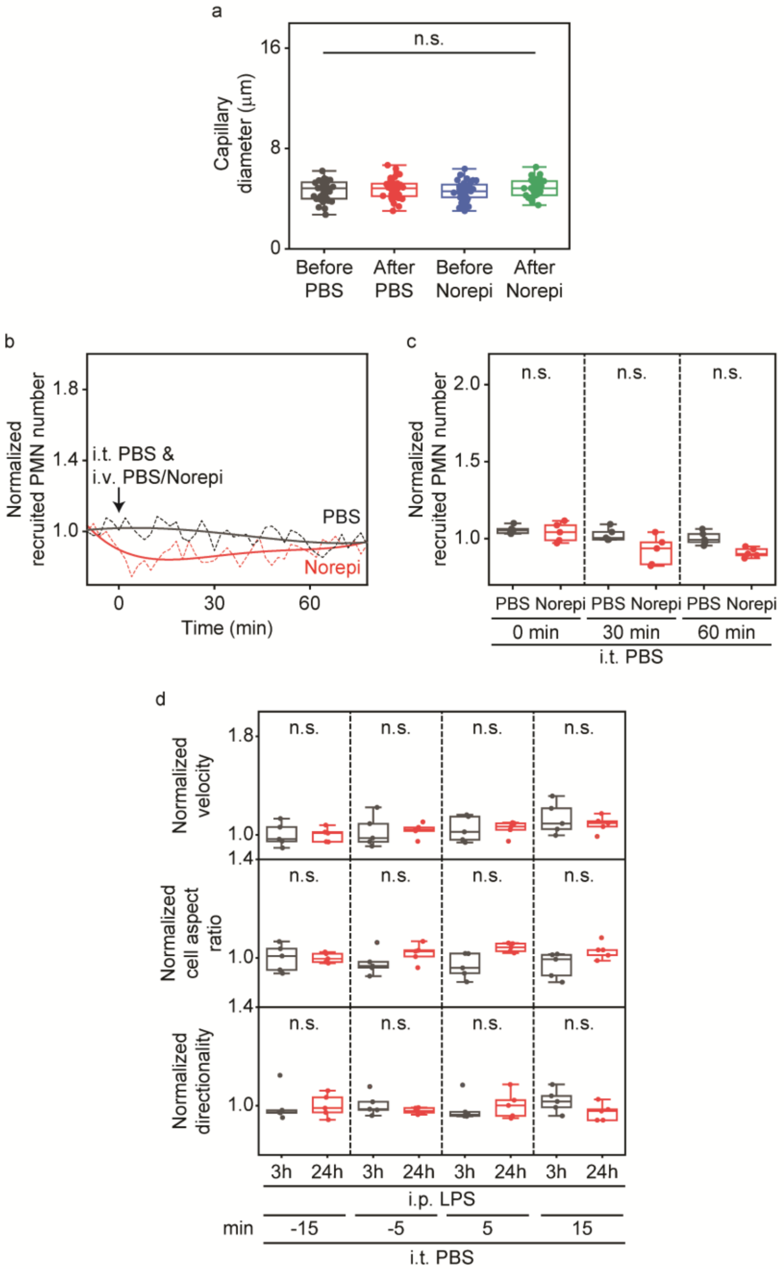


**Supplementary figure 7. Capillary Diameters in Response to PBS or Norepinephrine and PMN Cell Number, Velocity, Cell Aspect Ratio and Directionality Under Different Experimental Conditions in Normal Respiring Lungs.**

**a.** Quantitative analysis of diameters of microvessels in response to i.v. injection of PBS or norepinephrine (Norepi).

**b.** Recruited PMN number at times before/after i.t. inhalation of PBS and i.v. injection of

PBS/norepinephrine (Norepi). Solids lines indicate polynomial fitting results.

**c.** Quantitative analysis of recruited PMN number; n=5 (FOVs) in i.t. PBS i.v. PBS/Norepi.

**d.** Quantitative analysis of velocity, cell aspect ratio, and directionality before and after i.t. PBS at i.p. LPS3h/24h condition; n=900-1200 (cell) for i.p. LPS3h, n=1300-1800 (cell) for i.p. LPS24h.

Statistical analysis was performed using one-way ANOVA Tukey test in a, c, d.

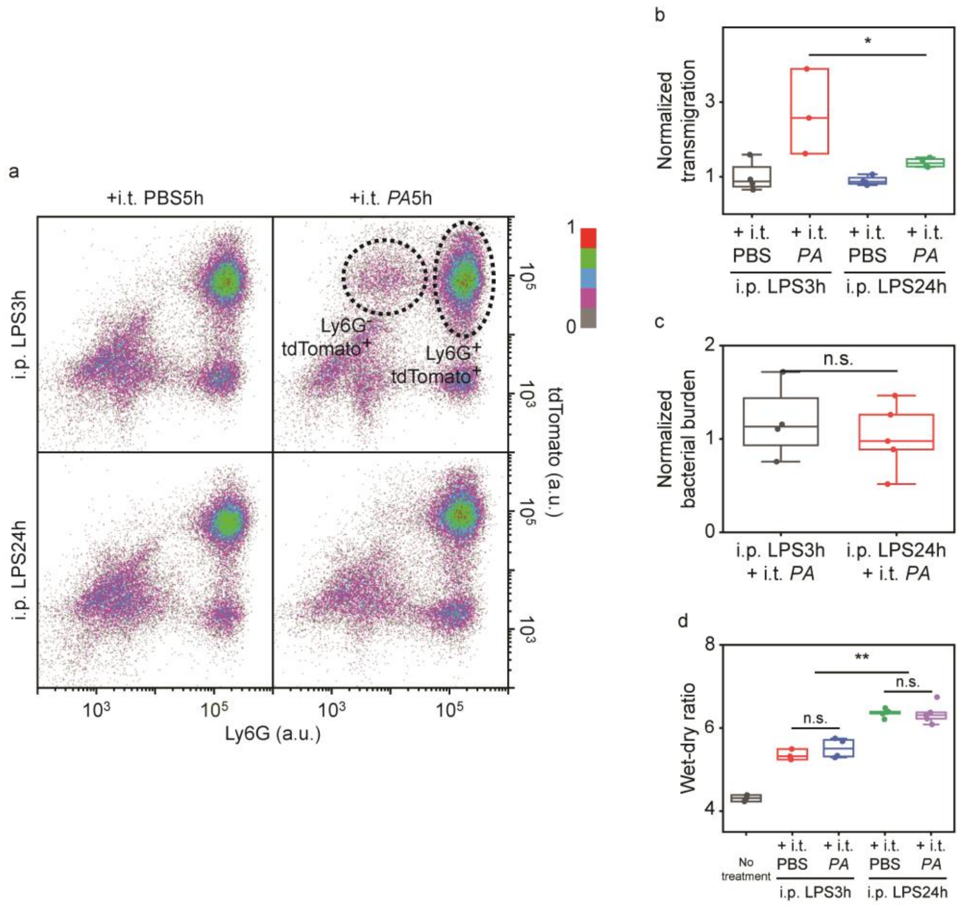


**Supplementary figure 8. Whole Lung Analysis of PMN Transmigration and Bacterial Burden and Wet/Dry Ratio.**

**a.** FACS analysis of PMN transmigration occurrence in response to i.t. *PA* injection at different time points (3h or 24h) after i.p. LPS; n=27.453-32,080 (Cell) for all cases.

**b.** Quantitative analysis of PMN transmigration in response to i.t. *PA* injection at different time points (3h or 24h) after i.p. LPS; n=4 (mice) for i.p. LPS3h + i.t. PBS, i.p. LPS24h + i.t. PBS and i.p. LPS24h + i.t. *PA*, n=3 (mice) for i.p. LPS3h + i.t. *PA*. Transmigration occurrences were calculated by transmigrated PMN number (tdTomato$^{+}$Ly6G$^{-}$ cell number) / total PMN number (tdTomato$^{+}$ cell number).

**c.** Quantitative analysis of bacterial burden of mice lungs in response to i.t. *PA* injection at different time points (3h/24h) after i.p. LPS; n=4 (mice) for i.p. LPS3h + i.t. *PA*, n=5 (mice) for i.p. LPS24h + i.t. *PA*.

**d.** Quantitative analysis of wet-dry ratio of mice lungs in response to i.t. *PA* injection at different time points (3h or 24h) after i.p. LPS; n=3 (mice) for no treatment and i.p. LPS3h + i.t. PBS, n=4 (mice) for i.p. LPS3h + i.t. *PA*, n=5 (mice) for i.p. LPS24h + i.t. PBS and i.p. LPS24h + i.t. *PA*.

Statistical analysis was performed using one-way ANOVA Tukey test in b-d; $*P<0.05$, $**P<0.01$.

**Supplementary movie1. Motion Artifacts in the Lung Optical Window.** Movie of pulmonary microvascular structure (Left). Movie of pulmonary microvessels with feature points (Right). Lung motion is acquired at video-rate and the motion of feature points are tracked by optical flow analysis. Scale bar; 1 mm. Time stamp; msec.

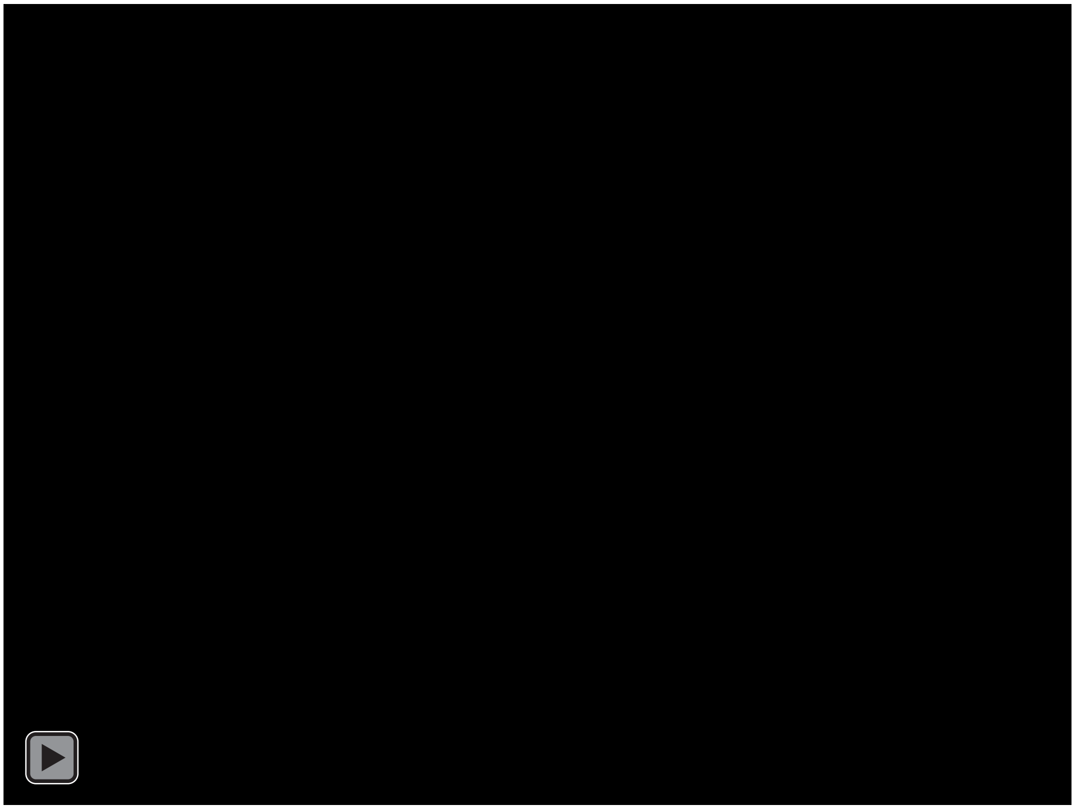

**Supplementary movie2. Stabilization Algorithm in CASTii.** Pulmonary microvessels are labeled by intravenously injected CD31 antibody. Shi and Tomasi method is used to detect feature points in reference image (red feature points in Motion tracking movie). These feature points are tracked using Lucas-Kanade optical flow method, where red feature points are derived from reference image and green feature points are tracked (Motion tracking image movie). Estimated motion is corrected using homography matrix calculated from matched feature points (Motion correction movie). Finally, the motion from respiratory cycle is averaged for each cycle (Respiratory cycle averaging movie).
Scale bar; 20 μm. Time stamp; msec. All two-photon movies are single z slices.

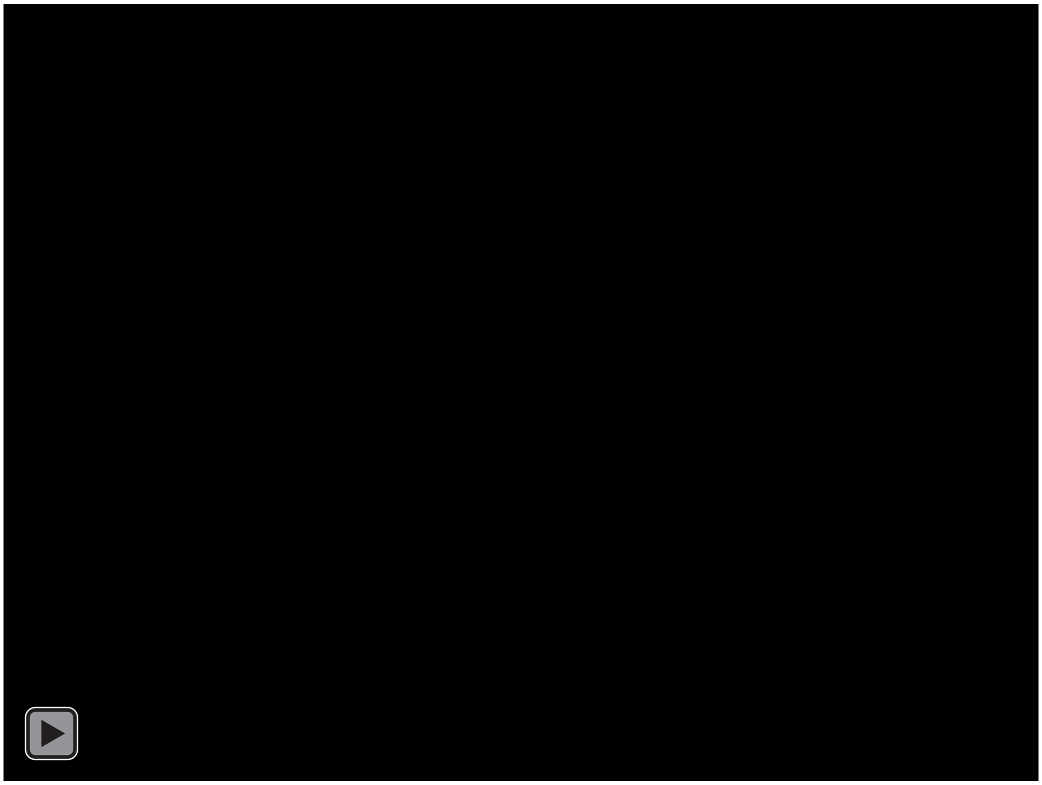

**Supplementary movie3. Compartmentalized Lung PMN Imaging after Intra-tracheal LPS Using CASTii.** Scale bar; 40 μm. Time stamp; min:sec. All two-photon movies are single z slices.

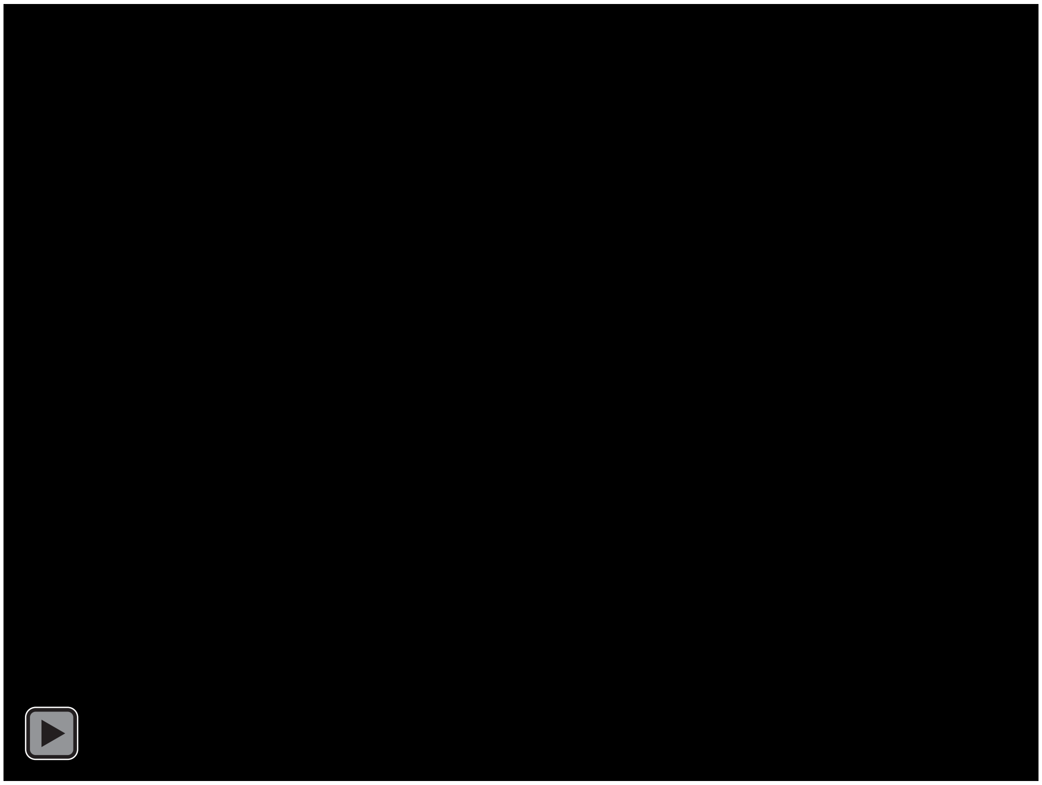

**Supplementary movie4. Representative lung PMN Images with Conventional Method and CASTii.** Left images are original images with conventional methods. Right images are stabilized images with CASTii.

Scale bar; 50 μm. All two-photon movies are single z slices.

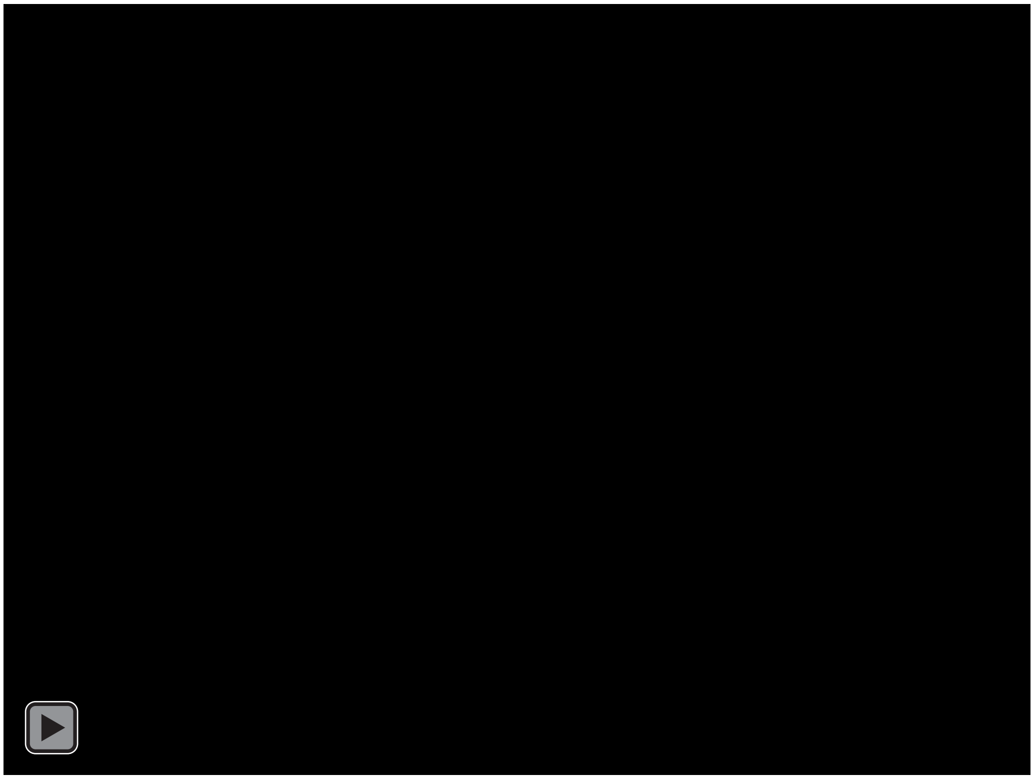

**Supplementary movie5. Comparison of Existing Image Registration Algorithms with CASTii and Comparison Among Motion Correction Algorithms in CASTii.** Our computer vision image registration algorithm (feature-based methods) was compared with the existing image registration algorithms, diffeomorphic demons as an example of the existing intensity-based methods. Motion correction algorithm in CASTii is compared with translational transform, where each of averaged XY motion displacements were horizontally moved back, perspective transform algorithm and non-rigid transform. Left image is original image before motion correction.

Scale bar; 20 μm. Time stamp; msec. All two-photon movies are single z slices.

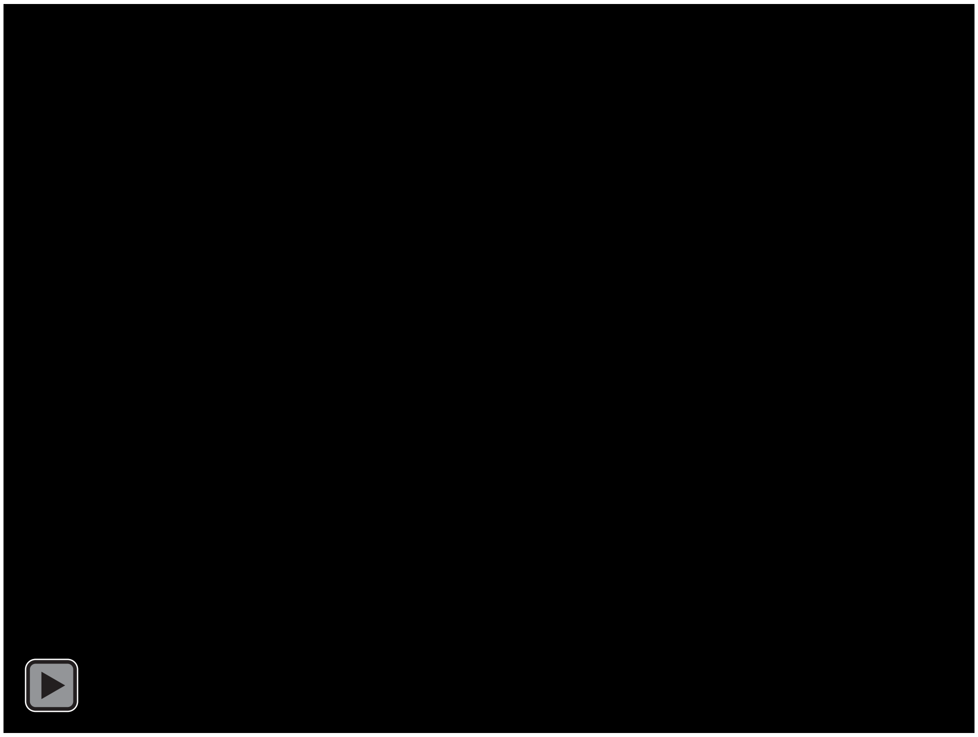

**Supplementary movie6. Passage of PMNs in narrow pulmonary microvessels.** Scale bar; 40 μm. Time stamp; min:sec. All two-photon movies are single z slices.

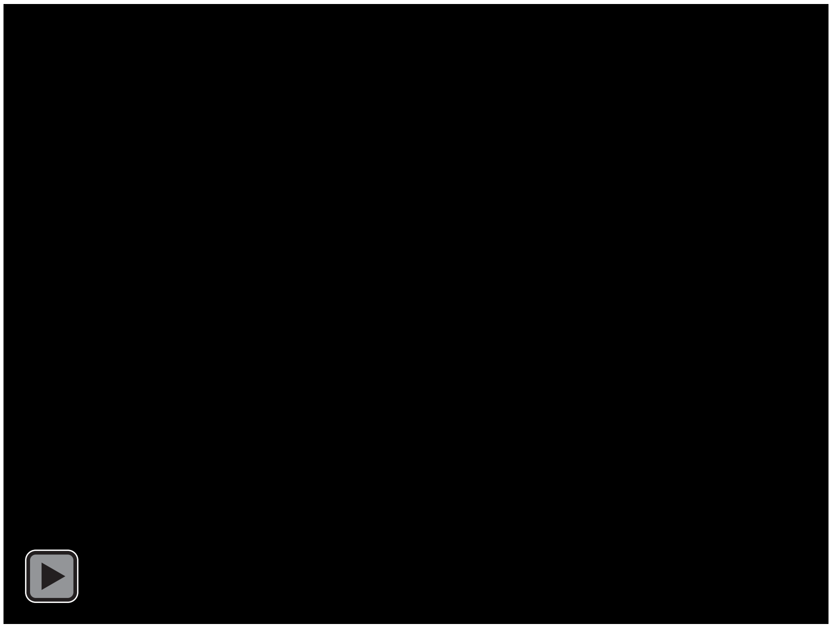

**Supplementary movie7. PMN circulation following intratracheal LPS challenge.** PMN turnover is observed in basal state and i.t. LPS. Scale bar; 40 μm. Time stamp; min:sec. All two-photon movies are single z slices.

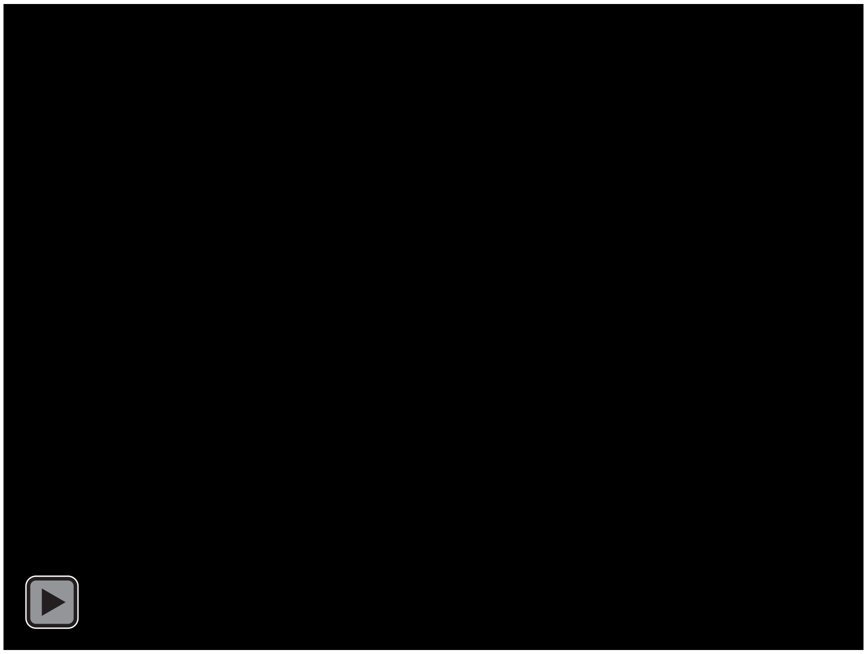

**Supplementary movie8. PMN Dynamics during Endotoxemia Induced with Intraperitoneal LPS.** PMN turnover completely disappears following i.p. LPS. Scale bar; 40 μm. Time stamp; min:sec. All two-photon movies are single z slices.

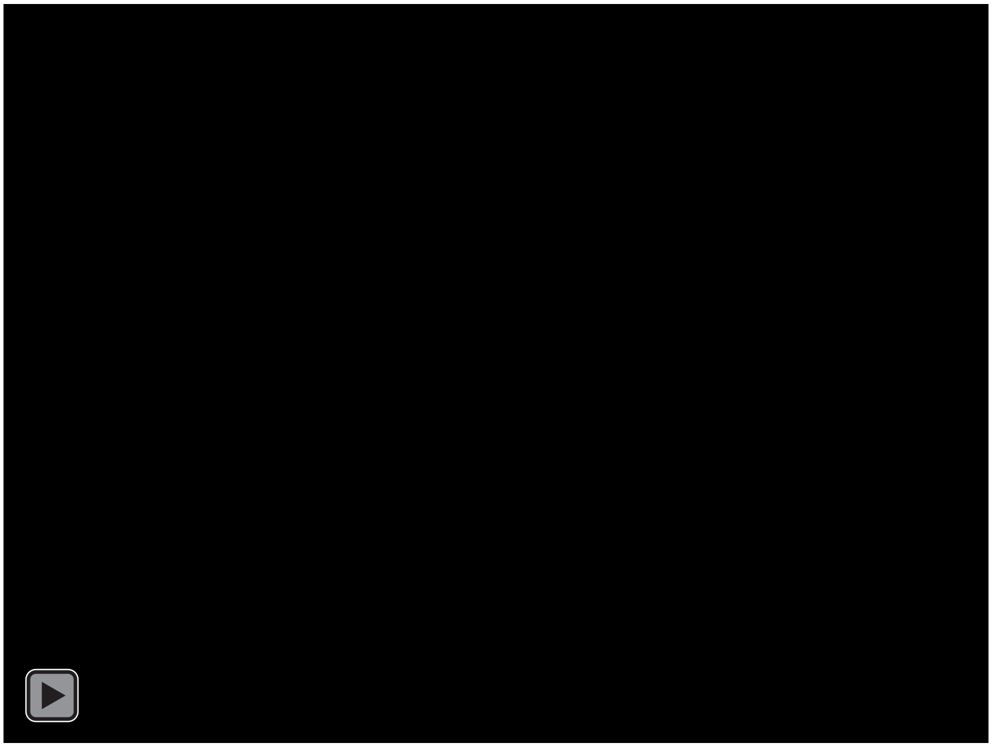

**Supplementary movie9. PMN Trafficking in Lungs during Endotoxemia induced by Intraperitoneal LPS.** In the condition of i.p. LPS, almost all PMNs attach microvessel endothelial cells and PMNs slowly move along microvessels. Migration velocity is progressively reduced after i.p. LPS.  Scale bar; 40 μm. Time stamp; min:sec. All two-photon movies are single z slices.

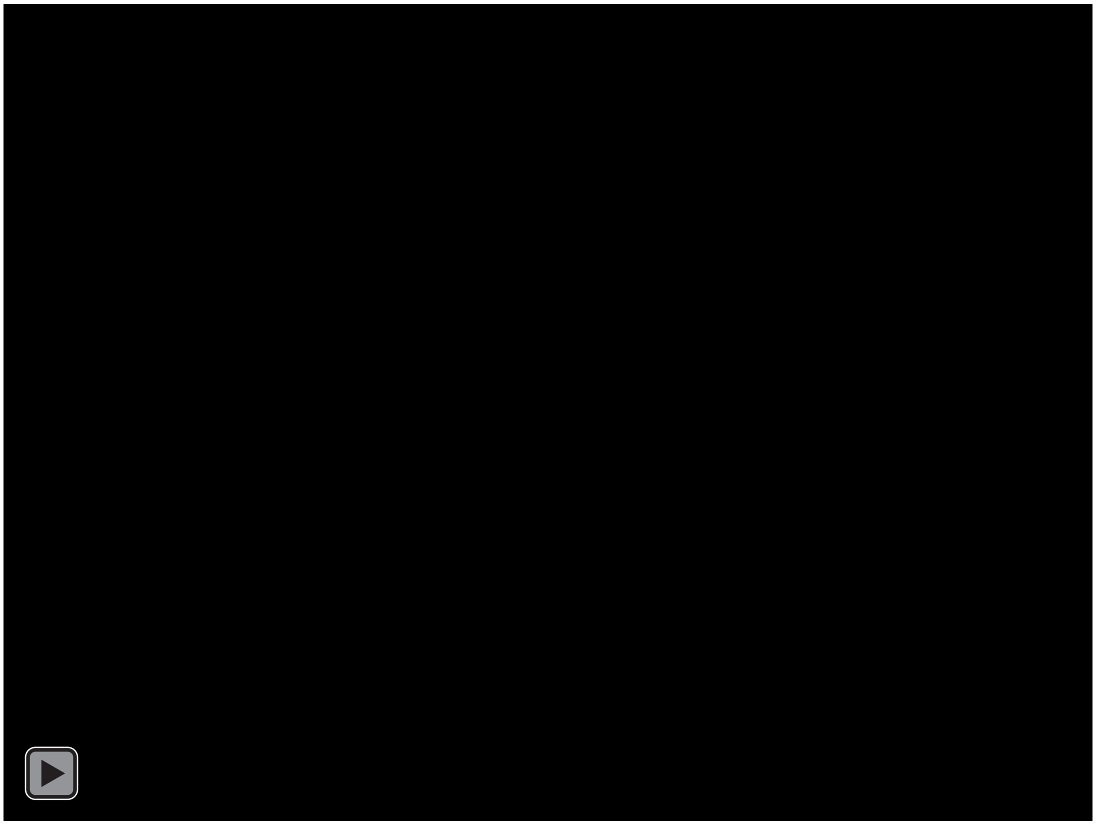

**Supplementary movie10. Effects of Norepinephrine on Dynamics of Lung PMN Marinated Pool.** After i.v. norepinephrine, PMN number is reduced and turnover rate of PMN increases.  Scale bar; 40 $\mu$m. Time stamp; min:sec. All two-photon movies are single z slices.

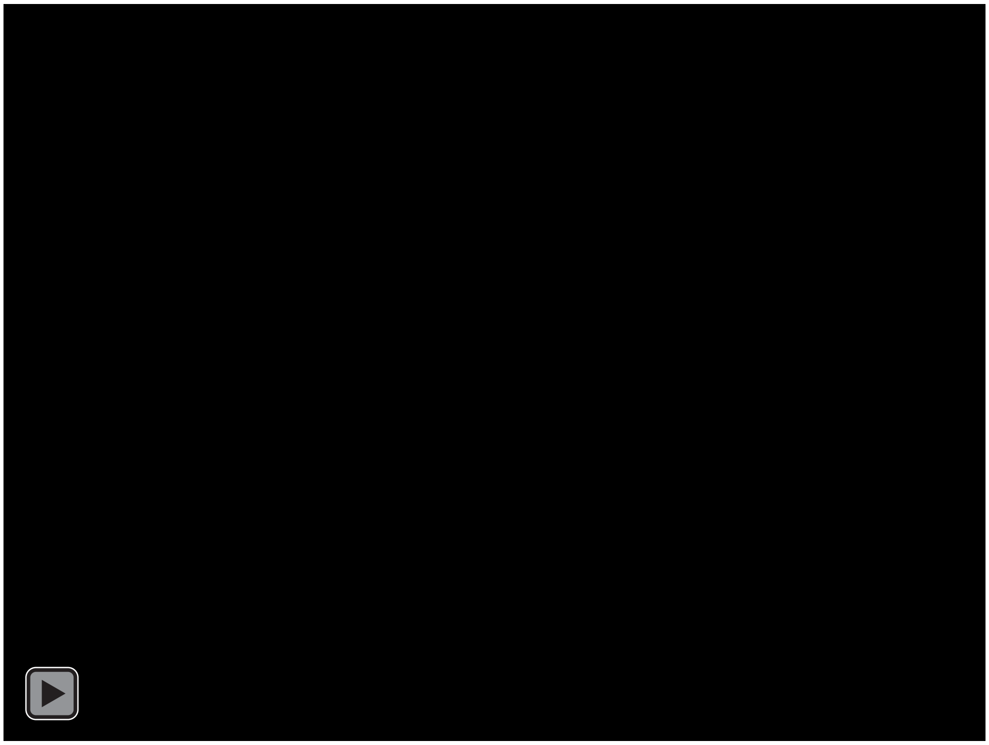

**Supplementary movie11. Effects of Norepinephrine on PMN Recruitment Dynamics Induced by fMLP Inhalation.** PMN dynamics after i.t. fMLP and i.v. PBS/norepinephrine, is visualized. Compared with i.t. fMLP or i.v. PBS, PMN recruitment is reduced and delayed by i.v. norepinephrine. Scale bar; 60 μm. Time stamp; hr:min:sec. All two-photon movies are single z slices.

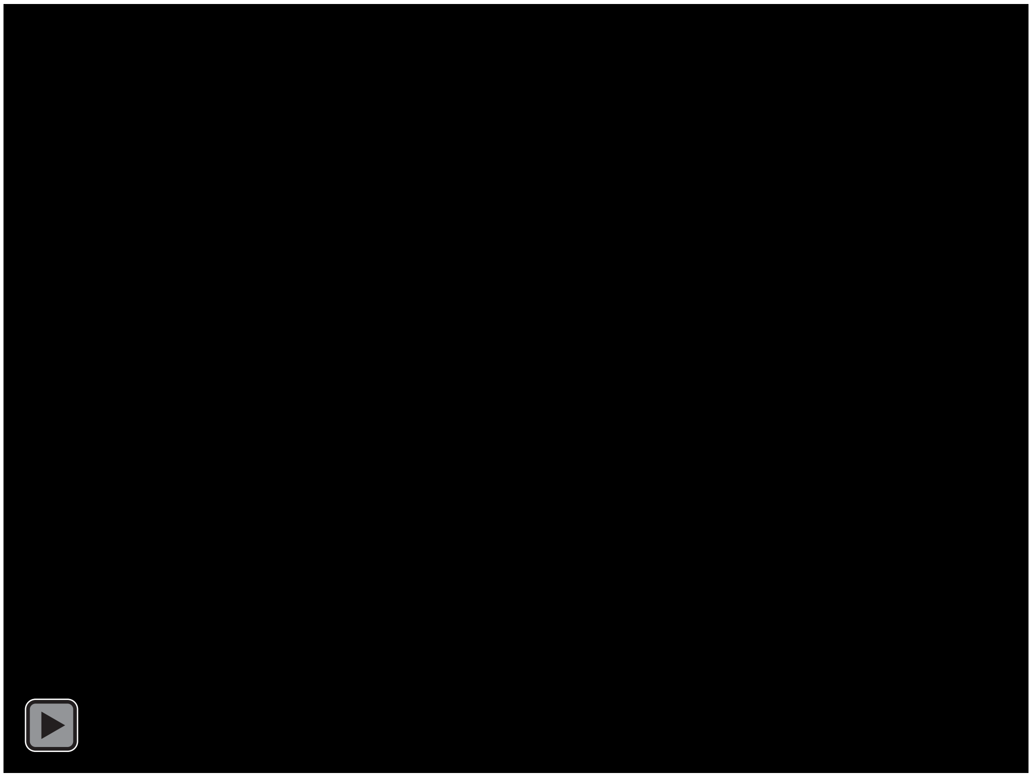

**Supplementary movie12. Effects of fMLP in 3h vs. 24h endotoxemia models.** After i.t. fMLP, PMN changes their migratory behavior at i.p. LPS3h condition but not i.p. LPS24h. Scale bar; 40 μm. Time stamp; min:sec. All two-photon movies are single z slices.

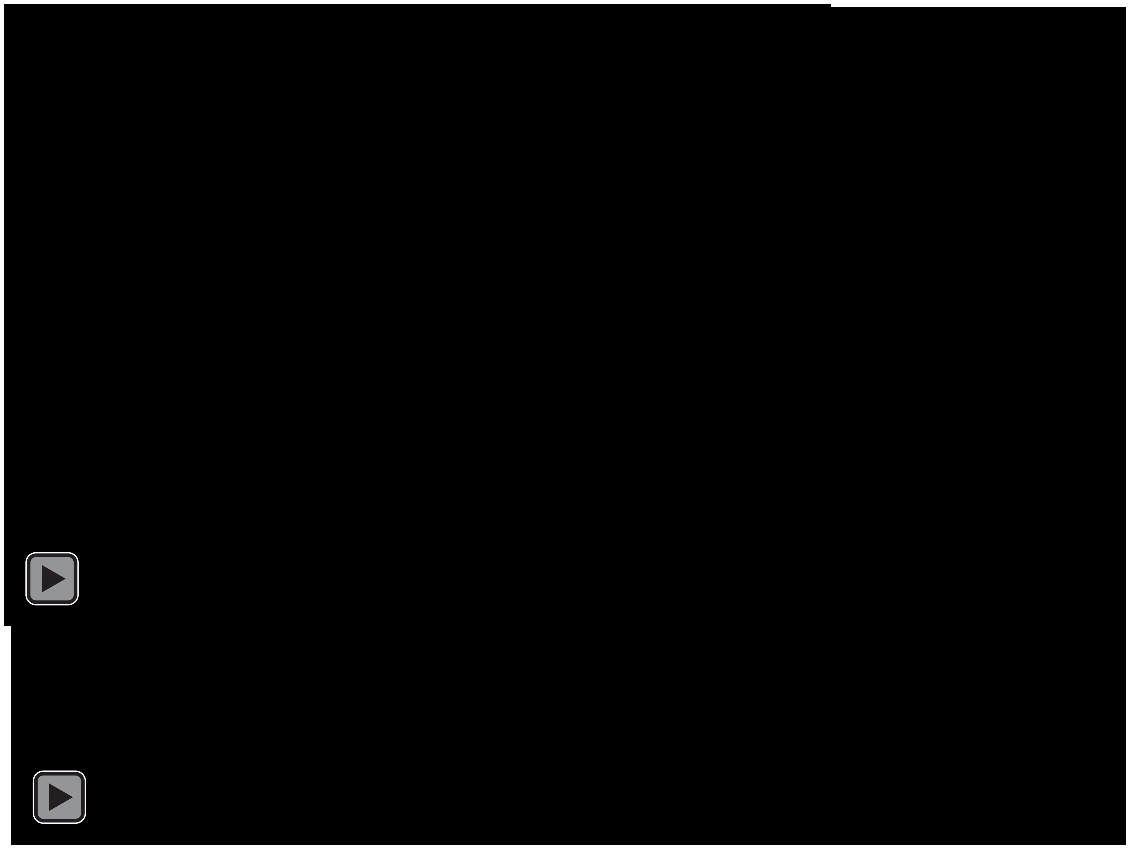
